# Telomere dysfunction impairs intestinal differentiation and predisposes to diet-induced colitis

**DOI:** 10.1101/2022.07.14.499988

**Authors:** Mindy Engevik, Lei Guo, Won-Suk Song, Christopher G Chronowski, Viktor Akhanov, Navish Bosquez, Amy C. Engevik, Numan Oezguen, Cristian Coarfa, Nagireddy Putluri, Andre Catic, Joseph A. Baur, Milton Finegold, Noah Shroyer, Jianhua Gu, Ronaldo P. Ferraris, Jennifer A Halliday, Christophe Herman, Simona Colla, Herman Dierick, Cholsoon Jang, Ergun Sahin

**Affiliations:** Department of Pathology & Immunology, Baylor College of Medicine, Houston, TX 77030, USA; Department of Regenerative Medicine & Cell Biology Medical University of South Carolina, Charleston, SC 29425, USA; Department of Population and Data Sciences, University of Texas Southwestern Medical Center, Dallas, TX 75390; Department of Biological Chemistry, University of California Irvine, School of Medicine, Irvine, CA 92697, USA; Huffington Center On Aging, Baylor College of Medicine, Houston, TX 77030, USA; Department of Pathology, Texas Children’s Hospital, Houston, TX, USA; Dan L Duncan Comprehensive Cancer Center & Center for Precision Environmental Health Baylor College of Medicine, Houston, TX 77030, USA; Department of Molecular and Cellular Biology, Baylor College of Medicine, Houston, TX 77030, USA; Advanced Technology Core, Baylor College of Medicine, Houston, TX 77030, USA; Department of Biochemistry and Biophysics, School of Medicine, University of Pennsylvania, PA 19104, USA; Molecular and Human Genetics, Baylor College of Medicine, Houston, TX 77030, USA; Department of Medicine, Section of Gastroenterology and Hepatology, Baylor College of Medicine, Houston, TX 77030, USA; Houston Methodist Research Institute, Electron Microscopy Core, Houston, TX 77030, USA; New Jersey Medical School, Rutgers University, Newark, NJ 07103, USA; Department of Leukemia, Division of Cancer Medicine, The University of Texas MD Anderson Cancer Center, Houston, TX 77030, USA; Department of Physiology and Biophysics, Baylor College of Medicine, Houston, TX 77030, USA

## Abstract

Intestinal epithelium dysfunction causes barrier defects, malabsorption and dysbiosis, predicting local and systemic disease, morbidity and mortality in humans. However, the underlying causes are not well understood. Here we show that telomere shortening is a host intrinsic factor that impairs enterocyte differentiation. The presence of such undifferentiated enterocytes is associated with barrier disruption and malabsorption of nutrients, such as fructose. A fructose-rich diet causes increased fructose spillover to the colon and induces colitis in a microbiome-dependent manner. The microbiome uses fructose to synthesize essential metabolites, including NAD precursors, that complement the host’s low NAD pool in the inflamed colon. Thus, telomere shortening drives enterocyte dysfunction and predisposes to diet-induced colitis through barrier disruption, increased nutrient flux to the colon and modulation of the microbiome. This differerentiation defect expands the canonical stem cell failure-centered view of how telomere shortening impacts the intestine and predisposes to intestinal disease in conditions associated with short telomeres.

## Introduction

The single-cell layer of the intestinal epithelium (IE) consists of several cell types – enterocytes, goblet cells, Paneth cells, enteroendocrine cells and M cells - that are regenerated from Lgr5(+) intestinal stem cells (ISCs) through a tightly controlled process under homeostatic conditions^1^. The IE is essential for the formation of the intestinal barrier that prevents the translocation of harmful bacterial products and antigens from the lumen into the intestinal tissue. The IE is also responsible for the digestion and absorption of nutrients, most of which are efficiently absorbed in the proximal intestine although indigestible nutrients such as complex carbohydrates or nutrients consumed in excess can reach the colonic microbiome more readily and change disease predisposition^2–4^.

Studies in humans and animal models have established the integrity of the IE as an important determinant for local and systemic disease predisposition and predictor of morbidity and mortality^5^. Barrier dysfunction is a hallmark of patients with inflammatory bowel disease, Crohn’s Disease (CD) and Ulcerative Colitis (UC) and it is a prognostic marker for increased risk^6,7^ and relapse^8,9^ in these patients. Patients with celiac disease have increased barrier permeability and healthy first-degree relatives of these patients show increased permeability, indicating that barrier dysfunction precedes disease manifestations^10^. Intestinal barrier integrity also plays an important role in the development of extraintestinal disorders, including type I diabetes and liver disease^11^. In critically ill patients and in the elderly, barrier dysfunction and intestinal inflammation predict mortality and frailty^12,13^. Besides barrier dysfunction, malabsorption is another important clinical consequence of patients with IE defects that can lead to systemic deficiencies of nutrients as well as an increased spillover of nutrients into the colon where it can impact microbiome composition and metabolism^14^. These human studies are supported by a large body of animal studies that indicate the importance of intestinal barrier as a regulator of intestinal and systemic inflammation and predictor of age-onset mortality^15^. However, a major challenge in this rapidly evolving field is our poor understanding of the underlying factors that drive barrier dysfunction. This would be important from a pathogenic view but also for the identification of at-risk patient patients.

A critical component of the IE are enterocytes, terminally differentiated cells that are essential for nutrient absorption and barrier integrity^16^. The execution of these different enterocyte functions depends on the formation of distinct functional membrane domains. The apical domain carries a brush border formed by microvilli, which carry digestive enzymes and transporters and increase the intestinal surface area for efficient nutrient absorption^17–19^. The basolateral domain contains junction proteins that form the selectively permeable barrier of the intestine^20,21^.

Impaired enterocyte differentiation can lead to barrier disruption, nutrient malabsorption, intestinal inflammation, and systemic growth retardation as shown in patients with loss-of-function mutations in genes essential for enterocyte maturation^22,23^. Enterocyte dysfunction is also observed in patients with UC^21,24,25^ and CD^7,26–28^ as well as in the elderly^29–31^ and is believed to be an important pathogenetic factor in IBD and age-related inflammation. A recent study in CD patients demonstrated that enterocytes are not fully matured and have abnormal microvilli as well as areas where microvilli are lost^32^. The underlying mechanisms for enterocyte and barrier dysfunction in IBD patients and in the elderly are not well understood.

The integrity of telomeres, the repetitive DNA sequences at chromosome ends, is critical for maintaining intestinal homeostasis. This is demonstrated directly in patients with severe telomere shortening due to loss-of-function mutations in the telomere-synthesizing enzyme telomerase (telomere biology disorders, TBD)^33–35^. A subset of these patients develops regenerative failure and several intestinal pathologies, of which inflammation of the colon (colitis) is most recognized^36^. Advanced telomere shortening is also a characteristic of patients with longstanding inflammatory bowel disease and implicated in disease progression and complications, including dysplasia and colorectal cancer^37,38^. Studies in human IBD tissues and organoids have further demonstrated the importance of dysfunctional telomeres for inflammation through activation of the DNA damage response^39^. Furthermore, telomere shortening is a hallmark of the intestine in older people and implicated in age-related diseases including colorectal cancer^40^.

Mechanistic studies in telomerase knockout mice with short telomeres (TKO hereafter), which recapitulate the predisposition to intestinal inflammation, have suggested that these pathologies are due to progressive intestinal stem cell (ISC) decline caused by several non-exclusive mechanisms^41,42^. Recent studies in intestinal organoids carrying patient-relevant mutations in telomerase have further substantiated the stem cell compromise^43,44^. Together, these studies have served as a basis to interpret the observed intestinal inflammation as a consequence of ISC decline, although how this - other than through near-total regenerative failure - causes intestinal inflammation is less clear. In contrast to the well-recognized stem cell compromise, the impact of telomere shortening on differentiation is not well understood.

Here, we report that telomere integrity is an important determinant of appropriate enterocyte differentiation as demonstrated by the formation of immature and functionally compromised enterocytes in TKO mice. The presence of these immature enterocytes is associated with a barrier defect and malabsorption that predisposes to diet-induced colitis in a microbiome-dependent manner. We identify NAD precursors as novel fructose-derived microbial metabolites that cross into the inflamed colon where NAD levels are low and critical for colitis. Thus, our findings indicate a novel role of telomere integrity in maintaining intestinal cell differentiation and predisposition to colitis in the context of a fructose-rich diet through modulation of the microbiome.

## Results

### Telomere shortening impairs enterocyte maturation and function

Previous studies have shown that telomere shortening in the intestine predisposes to inflammation through pathways that are incompletely understood and mainly attributed to ISC compromise^33,45,46^. To determine the impact of telomere dysfunction on the intestinal epithelium (IE), we performed RNA-seq analysis on the IE from the proximal intestine of young WT and TKO mice. Surprisingly, despite the absence of any apparent clinical or histological pathologies, a global comparison of differentially expressed genes segregated the two genotypes (**Fig. 1a, Supplemental Tables S1 and S2**). Gene ontology analysis indicated that nutrient transporters are among the most repressed genes in TKO mice (**Fig. 1b and sFig. 1a**). These repressed transporters cover a wide range of substrates including sugars, amino acids, and dipeptides (**Fig. 1c**). In addition, other enterocyte-specific genes encoding microvilli components, digestive enzymes, junction proteins, and signaling proteins important for cell polarization were also repressed in TKO mice. We validated some of these key transcriptomic changes in an independent set of mice by RT-qPCR and immunohistochemistry (**Fig. 1d, e**).

**Fig. 1.**
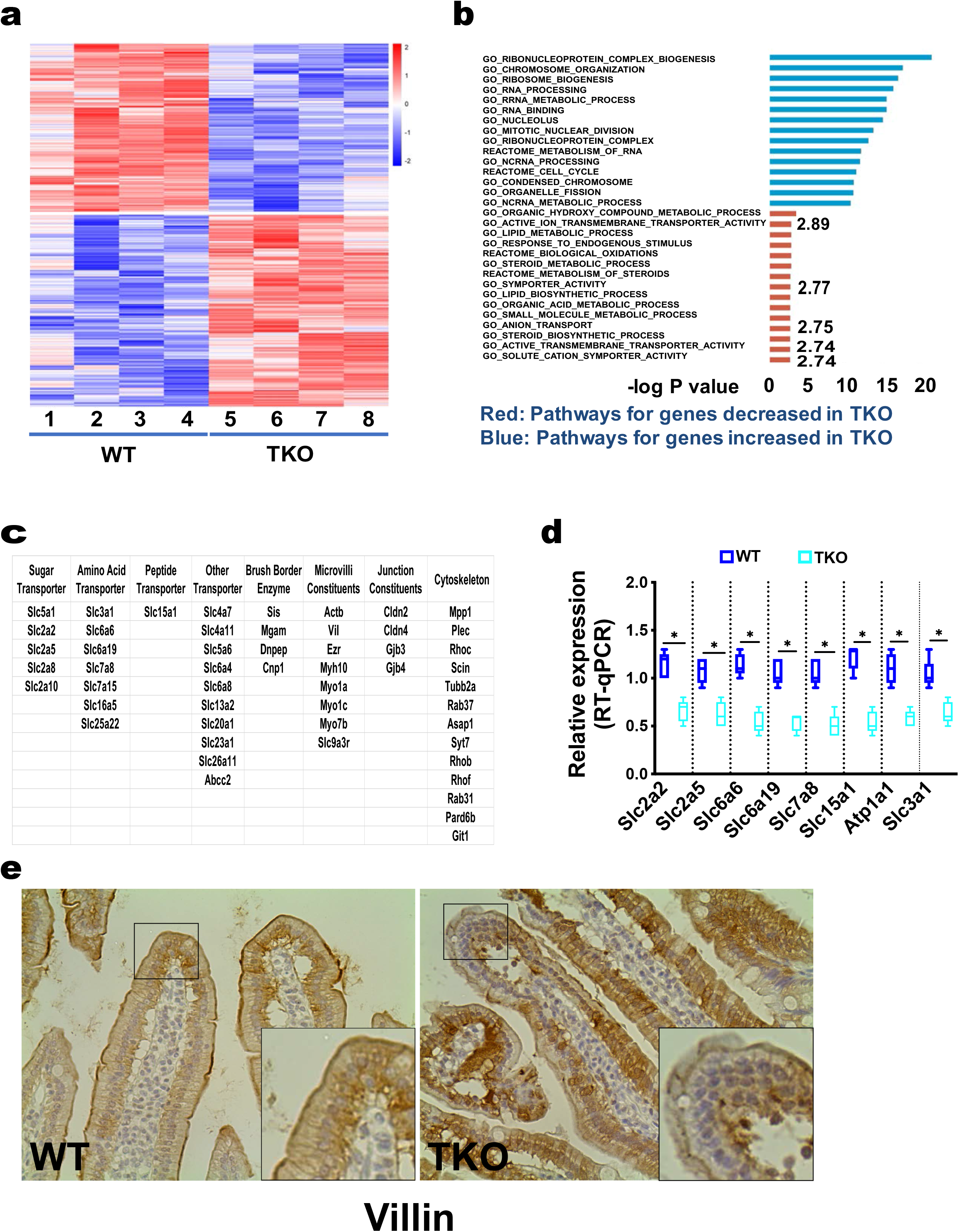
Telomere dysfunction decreases expression of transporters, enzymes, and junction proteins in the small intestine. RNA-seq was carried out on intestinal epithelial scraps from four 14-week-old male wild type (WT) and telomerase deficient mice with short telomeres (TKO); n= 4 per group. (a) Differential expression analysis shows that all four TKO intestines have very distinct transcriptional profile compared to WT intestines; differentially expressed genes defined as fold change > 2 & adjusted p < 0.05. (b) Top enriched GO terms of differentially expressed genes (DEG; Red: Enriched GO terms of the upregulated DEGs in TKO; Blue: Enriched GO terms of the downregulated DEGs in TKO). Transporters are among the most significantly downregulated genes in TKO mice: Active Ion Transmembrane Transporter (-log P: 2.89); Symporter (-log P: 2.77); Anion Transport (-log P: 2.75); Transmembrane Transporter (-log P: 2.74) and Solute Cation Symporter (-log P: 2.74). (c) Partial list of genes repressed in TKO mice, including sugar, amino acid, dipeptide transporters, brush border enzymes, microvilli constituents, junction components and cytoskeleton. (d) RT-qPCR validation of the indicated transporters in an independent set of TKO and WT mice; n= 5 per group; Student’s t-test with p < 0.05 considered to be statistical different. (e) Representative immunohistochemistry image of Villin shows its dysregulation in TKO small intestines; n= 3 per group.

To identify morphological changes in enterocytes, we analyzed enterocytes at different sections of the villus axis by scanning and transmission electron microscopy (SEM and TEM). SEM revealed discreet patches of microvilli loss in TKO enterocytes (yellow arrows in **Fig. 2a**). At higher magnification, the center of these areas is devoid of microvilli while the edges harbored irregular microvilli with alterations in microvilli packing and structure compared to the very uniform microvilli in WT. Such microvilli loss and abnormalities in TKO mice were observed along the entire villus axis (**sFig. 2a, b**). These changes were also observed in the jejunum and colon (**sFig. 2c and data not shown**). TEM imaging corroborated these findings; the patchy loss of microvilli and abnormal packing at low magnification and pearling instability at high magnification (**Fig. 2b and sFig. 2d**), similar to what has been reported in mice deficient for key differentiation-relevant transcription factors in the intestine^47^. Quantitation of microvilli length demonstrated significantly decreased length in TKO mice (**Fig. 2c**).

**Fig. 2.**
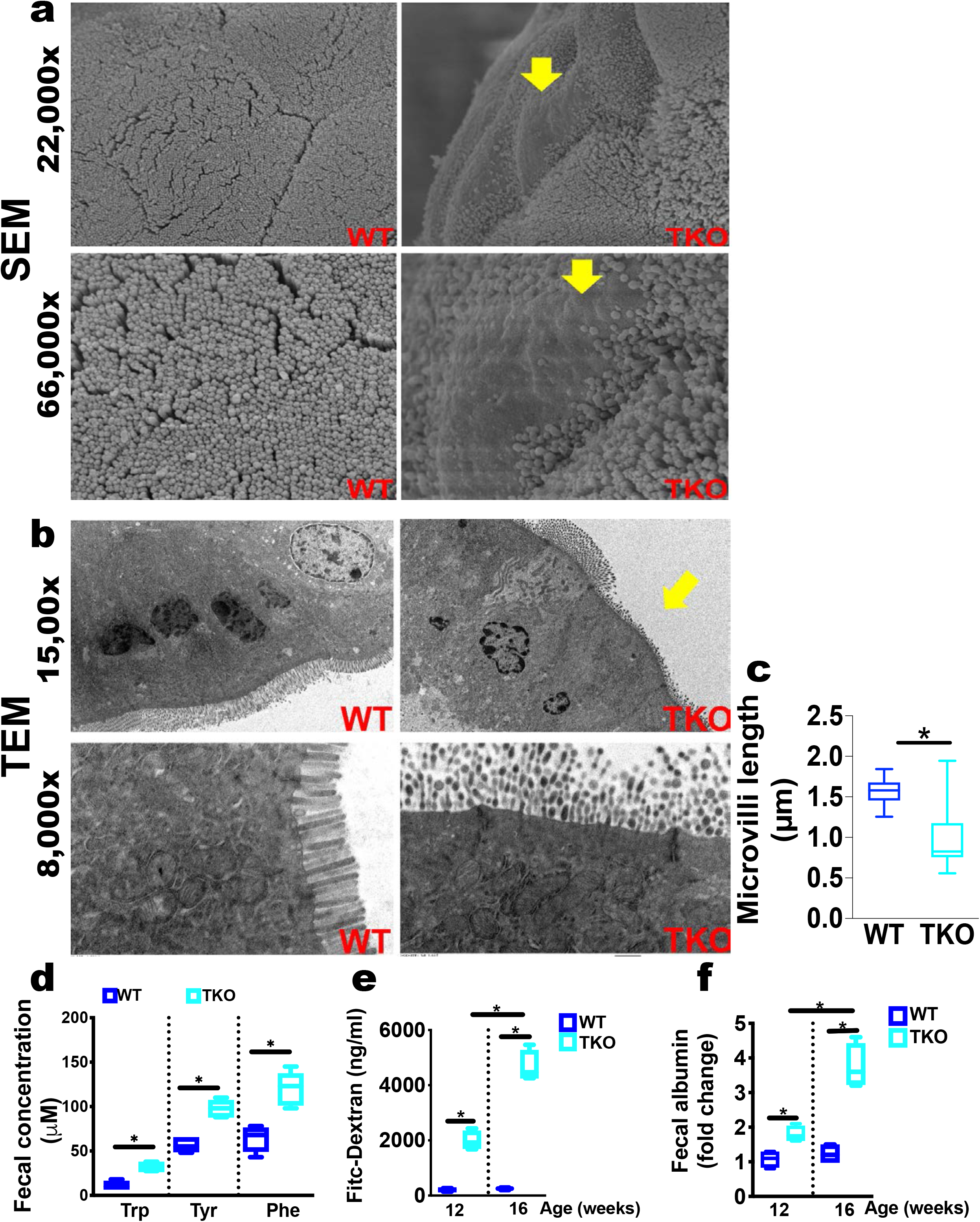
Telomere dysfunction impairs enterocyte maturation with loss of microvilli, malabsorption and leaky barrier. The small intestines of 14-16-week-old male WT and TKO mice were analyzed to assess morphology, absorption, and barrier function. (a) Scanning electron microscopy (SEM) analysis of the proximal small intestine at low (22,000x) and high magnification (66,000x) reveals patchy loss of microvilli in TKO mice. Yellow arrows point to areas with abnormal and diminished microvilli; n= 3 per group; representative images. (b) Transmission electron microscopy (TEM) analysis of the proximal intestine at low (1,500x) and high (8,000x) magnification shows abnormal microvilli (short, bent and pearl-like appearance) and microvilli-deficient areas in TKO mice; n= 3 per group; representative images. (c) Quantification of microvillus length indicates shorter microvilli in TKO mice; n= 3 per group; 10 images per mouse were analyzed. (d) Mass spectrometry-based quantification of tryptophan (Trp), tyrosine (Tyr) and phenylalanine (Phe) in feces shows their increased levels in TKO mice; n= 3 per group. (e) Blood Fitc-Dextran levels after oral delivery indicates abnormally elevated barrier permeability in TKO Mice; n= 3 per group. (f) Fecal albumin concentration indicates abnormally elevated barrier permeability in TKO mice; n= 3 per Group. Statistical differences were calculated using Student’s t-test with *p < 0.05 considered to be statistical different.

To determine the functional consequences of the transcriptional and structural changes, we measured intestinal nutrient absorption and barrier integrity. The absorption of amino acids was markedly impaired in TKO mice, reflected by increased levels of amino acids and peptides in feces (**Fig. 2d and data not shown**). Serum Fitc-Dextran and fecal albumin levels indicated barrier defects in young TKO mice (**Fig. 2e, 2f**). These phenotypes became progressively worse as mice aged. Together, these transcriptomic, ultrastructural, and functional analyses demonstrate that short telomeres are associated with defective enterocyte differentiation and functional compromise, including barrier dysfunction and malabsorption.

### Dietary fructose induces colitis in telomerase-deficient mice

We next sought to determine whether the presence of these immature enterocytes altered the ability of the intestine to process specific diets. The decreased expression of the main fructose transporter Glut5 (Slc5a1) (**Fig. 1c**) and the clinical relevance of increased fructose intake worldwide for colitis and colon cancer^48–51^ prompted us to investigate the effects of fructose in the setting of short telomeres. We found a significant reduction in fructose absorptive capacity in TKO mice (**Fig. 3a**). After gavage of ^13^C-fructose (with unlabeled glucose at 1:1 ratio to mimic high fructose corn syrup or sucrose intake), WT mice showed no fructose in feces, indicating complete absorption. In contrast, TKO mice showed high fecal fructose levels, which were almost ~50% of those in Glut5 knockout mice that showed impaired fructose absorption^52^.

**Fig. 3.**
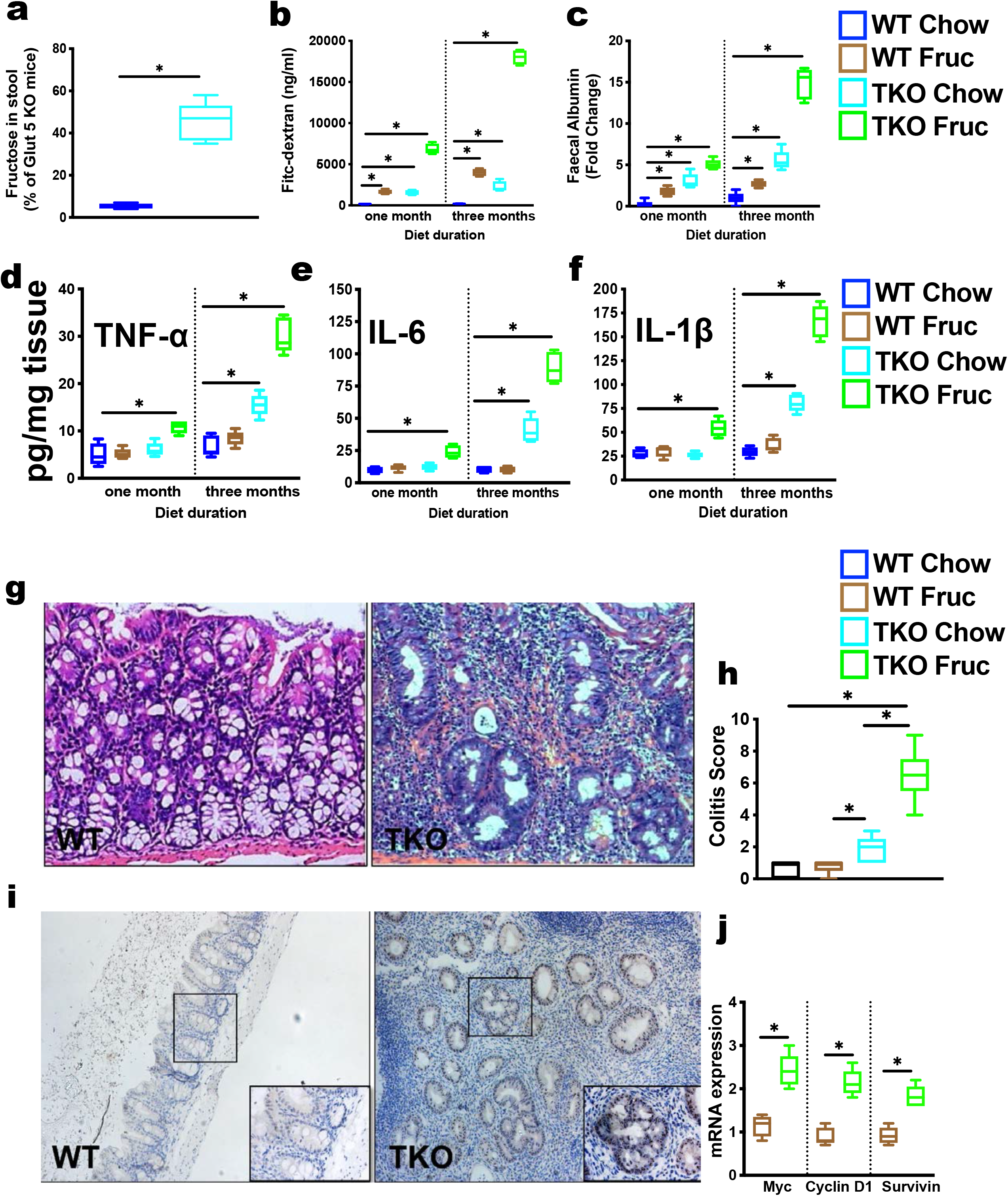
Telomere dysfunction impairs fructose absorption and predisposes to fructose-induced colitis and dysplasia. TKO and WT mice were fed a regular chow with normal drinking water or fructose -enriched water and analyzed after one and three months. (a) High fructose in stool after ^13^C-fructose gavage indicates fructose malabsorption in TKO mice. Results are expressed as percentage of fructose levels in Glut 5 knockout mice; n= 5 per group. (b, c) High blood Fitc-Dextran (a) and fecal albumin concentration (b) indicates fructose-induced severe barrier dysfunction in TKO mice in a time-depended manner; n= 5-8 per group. (d-f) Time-dependent increase of the indicated inflammatory cytokines in colonic tissue reflects fructose-induced colitis in TKO mice; n= 5 per group. (g, h) Representative H&E images of WT and TKO colons (g) after 3-month fructose feeding show severe colitis (h, colitis core); n= 10 per group. (i,j) Immunohistochemistry of c-Myc (i) and RT-qPCR of c-Myc targets (j) in TKO colons show dysplasia; n= 3 per group for IHC and n= 5 for RT-qPCR. Statistical differences were calculated using Student’s t-test with *p < 0.05 considered to be statistical different.

To assess the chronic effects of fructose consumption in the context of short telomeres, we fed TKO and WT mice with a fructose-rich diet (30% (w/v) in drinking water) commonly used to mimic high soft drink consumption^53,54^. TKO and WT mice showed similar fructose consumption and calorie intake as determined by the Comprehensive Lab Animal Monitoring System (CLAMS, data not shown). While fructose feeding modestly affected the barrier integrity in WT mice as previously reported^55,56^, it induced severe and progressive barrier disruption in TKO mice determined by Fitc-Dextran (**Fig. 3b)** and fecal albumin measurement (**Fig. 3c)**. Fructose feeding also worsened nutrient absorption in TKO mice, as reflected by substantially increased levels of amino acids (**sFig. 3a, b**) and dipeptides (**sFig. 3c, d**) in feces and corresponding decreases in blood.

These fructose-induced changes were paralleled by a time-dependent induction of colitis in TKO mice indicated by the progressive increase of proinflammatory cytokines TNFa, IL-6 and IL-1b in colonic tissue (**Fig. 3d-f**). At the histological level, fructose-induced colitis in TKO mice was also evident by erosion of the epithelial layer, ulcerations, and infiltration of inflammatory cells, reflected in a significantley increased colitis score (**Fig. 3g, h**). These changes coincided with the induction of known colitis and dysplasia markers, including the induction of c-Myc and its target genes (**Fig. 3i, j**), as well as nuclear β-catenin and high cell proliferative rate (**sFig. 3e**). Telomere shortening conferred a pronounced susceptibility to fructose as the colitis incidence increased significantly during the 2-, 3- and 4-months fructose diet (40%, 66%, and 75%) compared to the regular chow diet (0%, 10% and 30%, **sFig. 3f, g**). This pronounced pro-inflammatory effect of fructose in the setting of short telomeres contrasted with the absence of colitis in WT mice even after long-term fructose feeding **(sFig. 3f, g)**.

To determine whether other dietary sugars are also colitogenic in the setting of short telomeres, we carried out a side-by-side comparative study with fructose and glucose. Glucose and fructose consumption was comparable by CLAMS between TKO and WT mice (data not shown). In contrast to fructose, glucose had little effect on barrier function (**sFig. 3h, i**) or colitis incidence, which was similar to that in TKO mice on a regular chow diet (**sFig. 3j**). Given the importance of microbiome in fructose catabolism and its pathological roles^57–59^, we next tested whether the microbiome contributes to colitis and dysplasia. To this end, we treated TKO mice with an established combination of antibiotics to deplete the microbiome^60^. Notably, antibiotics treatment substantially suppressed barrier leakage (**sFig. 3k, l**) and colitis (**sFig. 3m, n**). Thus, poor fructose absorption and subsequently increased exposure of the colonic microbiome to fructose induces colitis and dysplasia in the setting of short telomeres. Together these data suggest that the microbiome is critical for fructose enhanced TKO colitis.

### NAD metabolism is dysregulated in fructose-induced colitis in TKO mice

To gain insights into the mechanisms through which fructose induces colitis when telomeres are dysfunctional, we performed a combined metabolomic and RNA-seq analysis on colonic tissue after 4-month fructose feeding. Both analyses revealed clear differences between WT and TKO mice. Metabolite Set Enrichment analysis (**Fig. 4a**) revealed that metabolites related to NAD and Tryptophan (Trp) were significantly altered in TKO colons. Indeed, metabolites in the kynurenine pathway - Trp, kynurenine and 3-hydroxykynurenine (3-HK) – were significantly changed in TKO colons (**Fig. 4b**). Consistent with the essential role of the kynurenine pathway in generating NAD from Trp catabolism (**sFig. 4a**), NAD, NADH and the NAD precursor nicotinamide (NAM) were all decreased in the inflamed colon (**Fig. 4b, c**). Strikingly, RNA-seq analysis (**sFig. 4b**) performed on the same colonic tissues subjected to metabolomic analysis showed upregulation of the enzymes involved in NAD synthesis, including the rate-limiting enzyme Ido1 (52-fold increase-top increased gene in the dataset) and Nicotinamide phosphoribosyltransferase (Nampt), the rate-limiting enzyme in the NAD salvage pathway that generates NAD from the NAM (2-fold, **Fig. 4d**). However, the expression of NAD-consuming enzymes, CD38 and eight of the known 17 PARPs, were also highly increased (**Fig. 4e**), suggesting that the observed NAD decline is due to increased NAD consumption that is not sufficiently supplemented by NAD biosynthesis. To understand whether these NAD-related changes occur prior to the development of colitis, we performed metabolomics (**sFig. 4c**) and expression analysis of Ido1 (**sFig. 4d**) at earlier time points. These analyses demonstrate that changes in NAD metabolism in TKO colons are an early event and occur prior to colitis indicating that they are pathogenetic drivers.

**Fig. 4.**
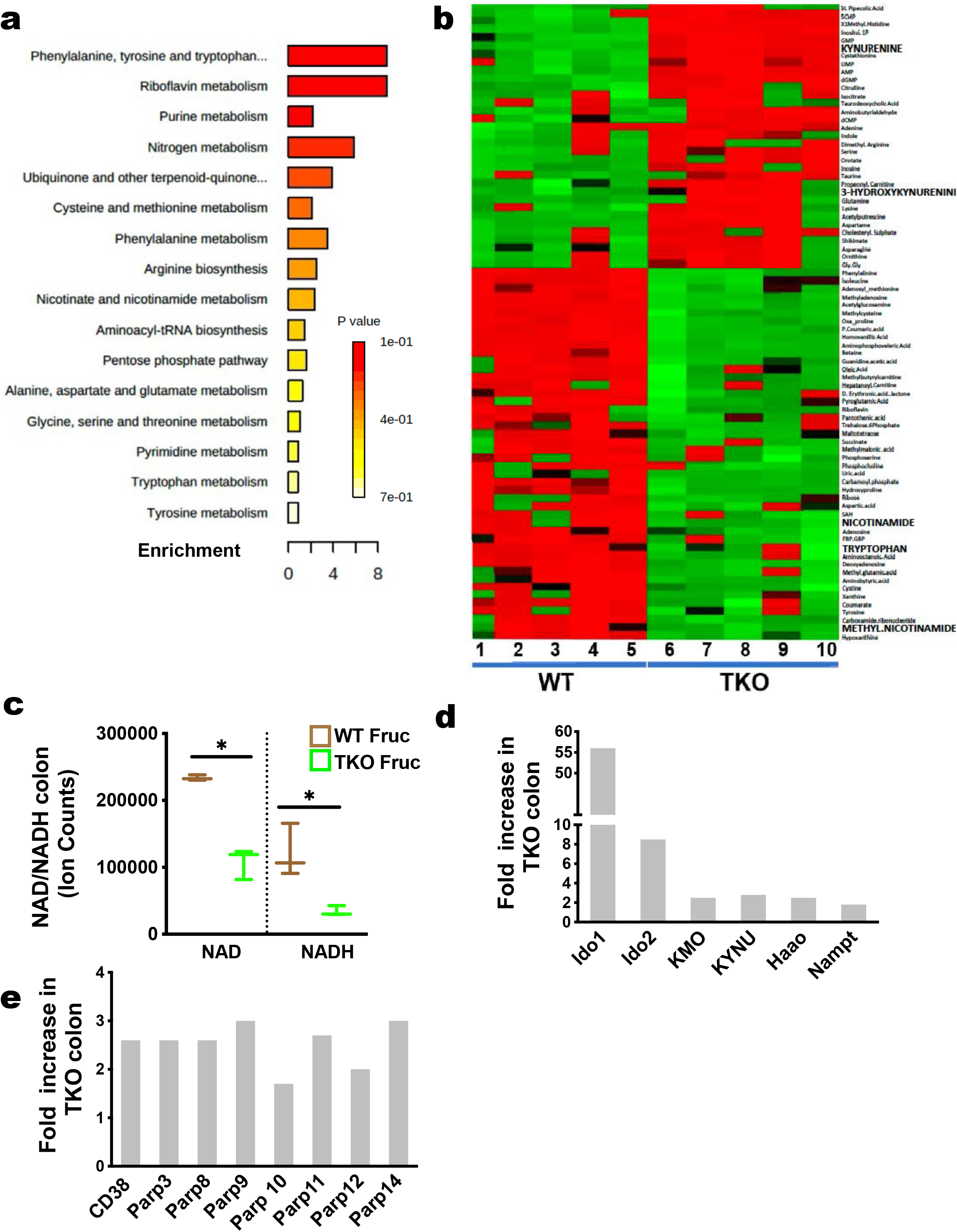
Fructose-induced colitis in TKO mice is associated with disrupted NAD homeostasis. Metabolomics and RNA-seq analysis was performed on colonic tissue in WT and TKO mice after feeding fructose in the drinking water for 3 months. (a, b) Metabolomics pathway enrichment analysis indicates altered tryptophan and its downstream pathways (kynurenine and NAD) in TKO colons; n= 5 per group. (c) LC-MS measurements of NAD and NADH levels show their depletion in TKO colons; n= 3 per group. (d) RNA-seq analysis indicates upregulated expression of tryptophan catabolism and NAD synthesis enzymes in TKO colons; n= 4 per group. (e) RNA-seq analysis indicates upregulated expression of NAD-consuming enzymes in TKO colons; n= 4 per group Statistical differences were calculated using Student’s t-test with *p < 0.05 considered to be statistical different. For RNAseq, differentially expressed genes are defined as fold change > 2 and p<0.05.

### The microbiome synthesizes NAD from fructose

Given the marked changes in NAD metabolism in TKO colons after a fructose diet and the improvement of colitis by antibiotics, we hypothesized that microbial fructose catabolism contributes to the colitis phenotype. Specifically, we wondered whether the microbiota uses fructose to generate NAD to supplement its deficiency in TKO colons. Like mammals, bacteria can generate NAD from Trp and, in addition, through an energetically simpler pathway that uses aspartate (Asp) (**Fig. 5a**)^61^. NAD is then used by several enzymes that release NAM, which can be reused to generate NAD via nicotinamide mononucleotide (NMN). In bacteria, NMN can be also converted to nicotinic acid (NA) and further converted to the NAD precursor, nicotinic acid mono nucleotide (NAMN) (**Fig. 5a**)^61^.

**Fig. 5.**
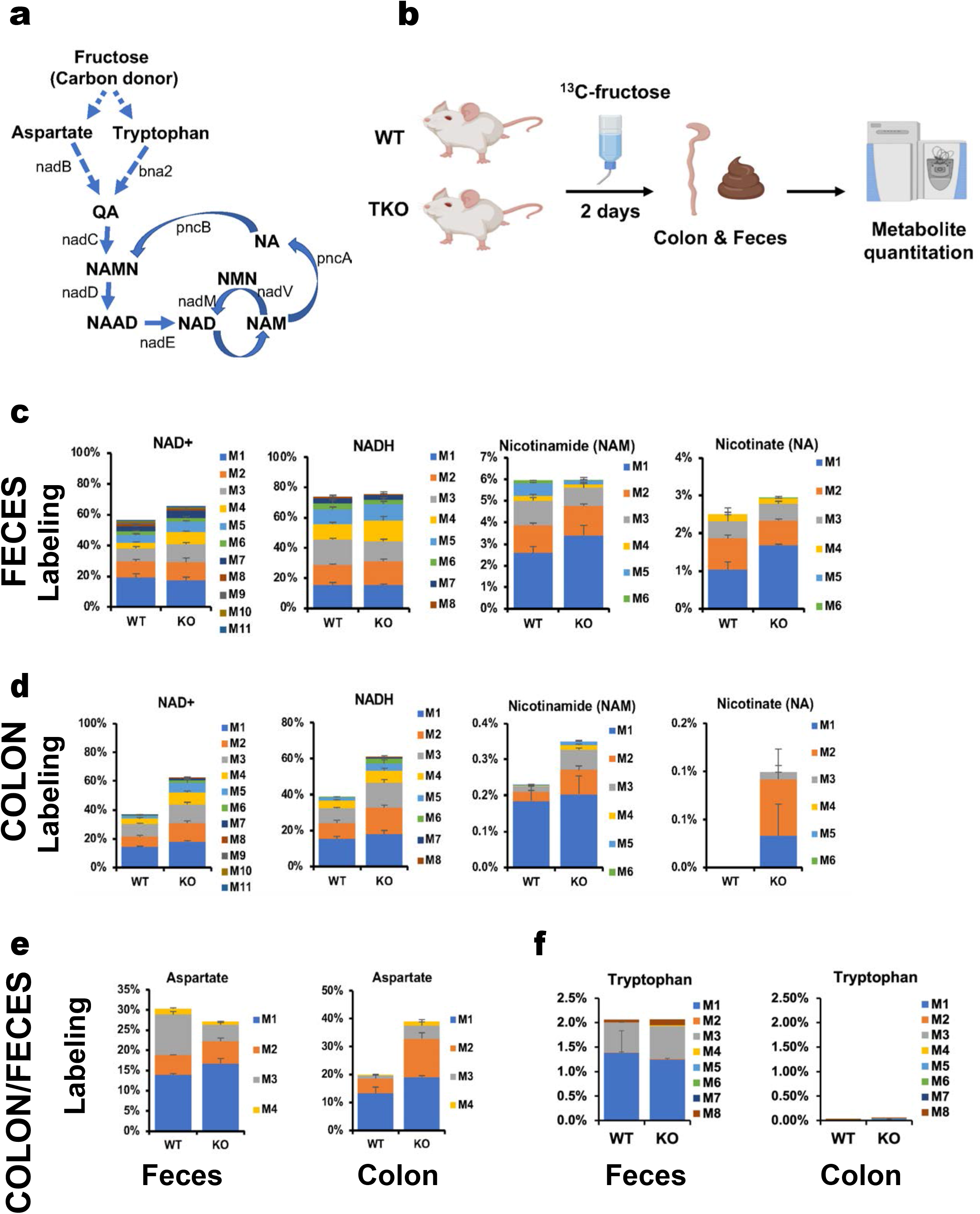
Fructose is used to generate NAD precursors and NAD by microbiota. WT and TKO mice were fed [U-^13^C]-labeled fructose in the drinking water for 2 days and metabolites in feces and colon were analyzed by LC-MS. (a) Scheme describing NAD-biosynthesis pathways in bacteria. (b) Experimental scheme for in vivo fructose isotope tracing in colon and feces. (c, d) Labeled isotopomer distribution of NAD, NADH and the indicated NAD precursors in feces (c) or colon (d) indicates their synthesis from fructose by gut microbiota; n = 3 per group. (e, f) Labeled isotopomer distribution of aspartate (e) and tryptophan (f) in feces and colon indicates aspartate as a major precursor of microbial NAD synthesis; n= 3 per group. Results are expressed as average ± s.e.

To determine whether the microbiome uses fructose to generate NAD and its precursors, we performed in vivo stable isotope tracing coupled with targeted metabolomics^62^. We fed mice universally ^13^C-labeled fructose in drinking water for 2 days to achieve steady-state labeling in feces and colon (**Fig. 5b**). Consistent with a significant contribution of fructose to NAD synthesis in gut microbiota, feces showed up to 70% of labeling in NAD and NADH in both genotypes (**Fig. 5c**). Moreover, free NAM and NA, both of which contain the pyridine ring de-novo synthesized from Asp or Trp, were also significantly labeled in feces, supporting their bacterial origin. Despite lower labeling, these labeled metabolites were also detected in the colon, suggesting that they are taken up by the colon (**Fig. 5d**). Importantly, all these metabolites showed higher labeling in TKO mice, reflecting enhanced NAD synthesis hinted from RNA-seq data (**Fig. 4d**). For example, approximately 60% of NAD and NADH in colonic tissue from TKO mice were labeled compared with 40% labeling in WT mice (**Fig. 5d**). Such a trend was also evident for NAM and NA (**Fig. 5d**). The labeling of NAM and NA was much higher in feces than those in the colon, further implicating their microbial origin and subsequent transfer to the colon (**Fig. 5d**).

To determine the relative contribution of the Asp versus Trp pathway to NAD generation in bacteria, we also measured Asp and Trp labeling **(Fig. 5e, f)**. Asp showed much higher labeling than Trp, suggesting that fructose is much more readily used to generate Asp, the simpler and energetically preferred pathway in bacteria. The labeling of Asp and Trp was also detectable in colonic tissue, with the latter only detectable at very low levels **(Fig. 5e, f)**.

We also performed a similar stable isotope tracing in *Drosophila* by feeding ^13^C-fructose as a sole carbon source to assess NAD production by the fly microbiome. Similar to what we found in mice, we detected high labeling of Trp, Asp, and other NAD precursors both in fly head and body, indicating that the microbiome contributes to the host’s NAD precursor pools in flies as well (**sFig. 5a**). Furthermore, as in mice, Asp (~ 20%) was more significantly labeled than Trp (~ 1.5%). The low labeling of Trp was in line with the low labeling of most essential amino acids in mice and flies, except for methionine, which is significantly labeled in feces (~ 7%) and colon (~ 3%, **sFig. 5b** and data not shown for flies). Together, our tracing study revealed microbial production of NAD precursors from fructose and its transfer into the colon with potential functional relevance for colitis in TKO mice.

### NAD is critical for ISC survival in the context of short telomeres

We next sought to identify which specific cell type carries this altered NAD metabolism. We first focused on ISCs given their importance for regeneration and transformation^63^. To this end, we generated TERT/Lgr5-Cre-IRES-GFP mice with varying degrees of telomere length to analyze Lgr5(+) cells with intact (WT) and short (TKO) telomeres. Lgr5(+) cells from TKO mice, but not those from WT mice, showed highly induced NAD-producing as well as NAD-consuming enzymes upon a high-fructose diet (**Fig. 6a, b**). This was paralleled by decreased NAD and NADH levels on TKO Lgr5(+) cells (**Fig. 6c**).

**Fig. 6.**
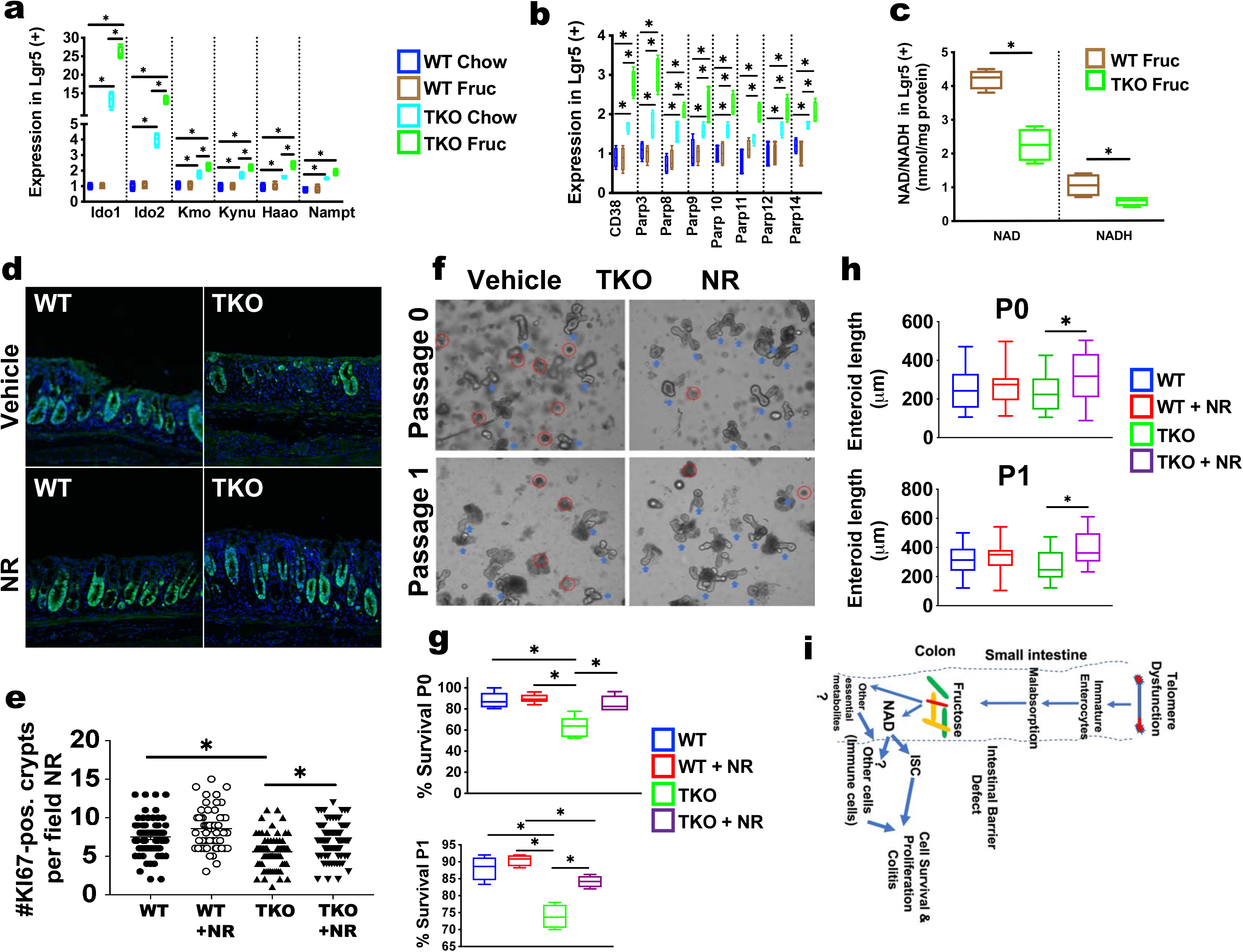
Increasing NAD levels via NR supplementation improves intestinal stem cell regeneration in TKO mice and enteroids. (a) NAD and NADH levels are decreased in TKO Lgr5(+) ISCs; n= 3 per group. (b) RT-qPCR analysis indicates upregulated expression of tryptophan catabolism and NAD synthesis enzymes in fructose-fed TKO Lgr5(+) ISCs; n = 3 per group. (c) RT-qPCR analysis indicates upregulated expression of NAD-consuming enzymes in fructose-fed TKO Lgr5(+) ISCs; n = 3 per group. (d, e) Representative KI67 immunofluorescence staining in colons (d) and quantification of regenerating crypts in 10 longitudinal images taken at 40x per mouse (e) shows increased colonic crypt numbers in TKO mice with NR supplementation; n = 8 per group. (f-h) Representative images of TKO and WT mice-derived enteroids (f) and quantitation of survival (g) and length (h) indicates increased survival and length of enteroids from TKO mice at isolation (P0) and after first passage (P1) with NR supplementation. Red circles indicate dying enteroids, n= 3 mice per group. (i) Proposed model by which short telomeres predispose to colitis following a fructose-rich diet. Telomere dysfunction induces enterocyte differentiation defect that leads to fructose malabsorption in the small intestine, resulting in spillover to the colonic microbiome. There, the microbiome (indicated by colored rods) use fructose to generate NAD/NAD precursors and other essential metabolites, which cross the defective barrier in the setting of short telomeres and enhance ISC survival and proliferation. Together with the potential impact of bacterial NAD on other cells such as immune cells, this contributes to chronic inflammation in colon. Statistical differences were calculated using Student’s t-test with *p < 0.05 considered to be statistical different.

To assess the effects of NAD restoration on ISC functions, we used a whole-body irradiation assay, a reliable and quantifiable experimental model system for studying ISC-mediated crypt regeneration in vivo^64^. Before irradiation, we treated TKO and WT mice with the NAD precursor, nicotinamide riboside (NR), or vehicle for 6 weeks to supplement NAD^65^. We then quantitated regenerating crypts 4 days post-irradiation. The colon of vehicle-treated TKO mice displayed a significant reduction in crypt numbers and extended sections of nearly empty regenerating crypts (**Fig. 6d, e**). In contrast, TKO mice treated with NR showed a significant increase in proliferating crypts (**Fig. 6d, e**). A trend towards an increase of regenerating crypts was also observed in WT mice treated with NR, although this did not reach statistical significance. This NR-mediated increased crypt regeneration was also evident in the small intestine (**sFig. 6a, b**).

The effect of increased NAD levels was further examined with an intestinal organoid model. The survival of crypts was significantly impaired in crypts derived from TKO mice at isolation and after passage (**Fig. 6f, g**). As we observed in vivo, the addition of NR increased the survival (**Fig. 6f, g**) and the size of organoids (**Fig. 6h**) from TKO mice while it did not have any significant effect on WT organoids’ survival or size (**Fig. 6f-h and sFig. 6c, d**). Thus, ISCs with short telomeres have lower levels of NAD and the increase of NAD levels improve their survival and regeneration.

Together, our studies reveal that telomere dysfunction induces differentiation defects of enterocytes, which are functionally compromised as reflected by the increased barrier permeability and nutrient malabsorption. Such malabsorption leads to increased spillover of nutrients such as fructose, which, in the setting of short telomeres and impaired barrier, drives colitis in a microbiome dependent manner. Fructose is utilized by the microbiome to synthesize and provide NAD and NAD precursors to the the inflamed colon with low NAD levels. The survival of IScs with short telomeres is improved when NAD is reconstituted and this maintains a pool of damaged ISCs that sustain the inflammatory process and can undergo malignant transformation (**Fig. 6i**).

## Discussion

Our multi-omics, structural and functional analysis demonstrates that dysfunctional telomeres cause a differentiation defect of enterocytes. This defect is observed even in young mice with preserved intestinal architecture, indicating that it is an early consequence of short telomeres, which precedes ISC failure. This finding expands our current stem cell failure centered view and provides an alternative, albeit non-exclusive, view on how telomere shortening leads to intestinal inflammation.

The presence of immature enterocytes explains the observed barrier defect in mice and, potentially, in human patients with intestinal inflammation, in which a barrier defect has been suggested to drive intestinal inflammation^36,45,66^. This barrier defect predisposes to inflammation through a pathological interaction with the microbiome based on the significant amelioration of inflammation when the microbial load is reduced after a fructose-rich diet or in aged TKO mice^45,46^. The importance of the intestinal epithelium as a driver of intestinal inflammation is further strengthened by the finding that intestinal inflammation is reduced when telomerase is reactivated specifically in the intestinal epithelium^45^. In this scenario, the disruption of the intestinal barrier is an early telomere shortening-associated event that allows the continous translocation of bacterial products to fuel the inflammatory process. It is possible that, with aging and increased inflammation, the enterocyte defect becomes more pronounced and, combined with stem cell failure at later stages, contributes to disease progression and complications.

Another important consequence of the enterocyte maturation defect is impaired nutrient absorption, as reflected in the increased spillover of nutrients into the colon. Malabsorption is a well-recognized feature in patients with intestinal inflammation, due to telomerase mutations, and in IBD patients^14,36^. While the systemic deficiency of nutrients can be addressed with parenteral substitution, the functional consequences of the increased flux of nutrients into the colon are not well understood in these patients^14^. In this regard, our studies provide important insights into how the enterocyte defect in the small intestine and subsequent malabsorption can drive large intestinal disease. Fructose consumption has risen significantly over the last decades worldwide and is linked to increased risk of intestinal (colitis, colorectal cancer) and systemic diseases^67,68^. We find a pronounced colitogenic effect of fructose in TKO mice with a pronounced barrier defect, but not in WT mice; indicating that short telomeres are an inherent host-specific risk factor for colitis through the combined effect of increased spillover of fructose in the colon, severe barrier defect and the microbiome ^55,56^. Glucose did not cause colitis but rather provided benefits^69^, likely due to the high glucose-absorptive capacity of the small intestine.

Previous studies have shown the significant impact of fructose on the microbiome^48,49,56^. Our in vivo stable isotope tracing revealed for the first time that fructose is actively utilized by the microbiome, in both mice and flies, to synthesize NAD and NAD precursors. Importantly, these metabolites readily cross the intestinal barrier and feed the colonic tissue in TKO mice. The significant increase of fructose-derived NAD/NAD precursors in the colon of TKO mice suggests that the flux of microbial metabolites is facilitated by the pronounced barrier leakage caused by critically short telomeres. Although the functional relevance of the microbiome-derived NAD/NAD precursors needs to be tested in future works, they could become quantitatively important for the host in conditions with high NAD consumption, such as inflammation^70^ or increased DNA damage due to short telomeres^71^. This view is supported by increased ISC survival when NAD levels are restored via NR supplementation. While we focused on the effect of NAD in ISCs as the cellular pool for malignant transformation^72^, NAD can be critical for immune cell survival and activity, which is also likely involved in colitis development^70^ (**Fig. 6i**).

The mechanisms underlying the differentiation defect are currently unknown. However, the molecular pathways, through which short telomeres control ISC survival, can provide some clues. Such examples include the DNA damage response, p53-dependent pathways^41,73,74^ and dysregulated Wnt signaling^43,66,75^. Interestingly, p53 activation impairs hepatocyte maturation in human organoids with short telomeres through direct repression of the hepatocyte differentiation transcription factor HNF4α^76^, which also plays roles in enterocyte differentiation^77^. Alternative mechanisms can involve mitochondrial dysfunction, as mice deficient for the mitochondrial transcription factor, TFAM, show enterocyte maturation defects and TKO mice have compromised mictochondrial biogenesis and function^78,79^. Future single-cell analyses will be instrumental in the identification of transcription factors that underlie the short telomere-induced enterocyte maturation defect.

Our findings have clinical implications for understanding the enterocyte and barrier defects in broad gastrointestinal pathologies, including IBD, celiac disease^80^, graft versus host^81^ and in the aged^82^. IBD patients develop accelerated telomere shortening because of increased cell turnover and insufficient telomerase activity, which is implicated in disease progression and complications including colorectal cancer^37^. Enterocyte dysfunction and increased intestinal permeability are prominent features and a major pathogenic factor in these patients, as demonstrated by the the predictive value for disease development and recurrence^8,26^. A recent study in CD patients demonstrated that enterocytes from non-affected areas are immature and show decreased expression of differentiation markers genes, including the ones that regulate microvilli structure and function^32^. Indeed, ultrastrucrual analysis showed microvilli abnormalities, such as microvilli loss or reduced length^32^, which we observed in young TKO mice. Thus, our findings raise the possibility that this could be a downstream consequence of dysfunctional telomeres. Similar enterocyte and barrier defects have been reported for celiac disease^83^ and chronic graft versus host disease^84^.

Telomere shortening is also a hallmark of the aged intestine and predisposes to age-related intestinal disease. While studies in organoids derived from the elderly have shown reduced formation efficiency^85^, indicative of a stem cell compromise, the effect of telomere shortening on differentiation and function of the intestine is not known. In particular, a region-specific analysis would be important, as heterogeneity in telomere length is observed in the intestine, including shorter telomeres in neoblastic adenomas that are invariably characterized by inflammation^86^. It remains to be established whether the shorter telomeres in the neoblastic adenomas is accompanied by differentiation and barrier defect. Establishing the role of short telomeres in human intestinal aging is important, given the growing body of evidence that an altered microbiome and barrier dysfunction are important factors for local and systemic aging decline and disease^15,31,87^.

Looking forward, our findings in the gut have potential implications for understanding the pathogenesis of other organ disorders associated with telomere shortening. TEM and SEM analysis reveal ultrastructural abnormalities in TKO enterocytes. Microvilli are crucial for different cell types including kidney epithelial cells, lymphocytes and tuft cells in the lungs^88^. Whether the role of telomere integrity in the formation of microvilli holds true for these cells or, more generally, other essential cellular structures in different types of cells remains to be established. Of particular interest will be to determine whether the differentiation of epithelial cells in other organs - such as the lung and the skin - is compromised by telomere shortening given that these organs are affected in TBD patients and in the aged^89^. Of note, lung fibrosis is a common manifestation in TBD patients, and a perturbed lung barrier may predispose to continued inflammation and long-term fibrosis^90^.

## Acknowledgments

This work was in part supported by the Ted Nash Long Life Foundation, Edward Mallinckrodt Jr. Foundation and NIA R01 grant (R01AG047924) to E. S., AASLD Foundation Pinnacle Research Award in Liver Disease, the Edward Mallinckrodt, Jr. Foundation Award, and NIH R01 grant (R01AA029124) to C. J. W.S.S was supported by the Korea Health Technology R&D Project through the Korea Health Industry Development Institute funded by the Ministry of Health & Welfare of Korea (HI19C1352). R.P.F. was supported by IH Grant R01-AT010243 and NSF Grant No. IOS 1754783. This study was in part supported by NIH grants DP1-AI152073 (C.H). Histological services were supported by PHS grant P30DK056338 to the Texas Medical Center Digestive Diseases Center. This project was also supported by the Cytometry and Cell Sorting Core at Baylor College of Medicine with funding from the CPRIT Core Facility Support Award (CPRIT-RP180672), the NIH (CA125123 and RR024574) and the assistance of Joel M. Sederstrom. Imaging for this project was supported by the Integrated Microscopy Core at Baylor College of Medicine with funding from NIH (DK56338, and CA125123), CPRIT (RP150578, RP170719), the Dan L. Duncan Comprehensive Cancer Center, and the John S. Dunn Gulf Coast Consortium for Chemical Genomics. The metabolomics core was supported by the CPRIT Core Facility Support Award RP170005 “Proteomic and Metabolomic Core Facility,” NCI Cancer Center Support Grant P30CA125123, intramural funds from the Dan L. Duncan Cancer Center (DLDCC).

## Author Contributions

E.S., M.E. and C.Y. developed the general concept, ideas, and research strategies. E.S. M.E., W.S.S., C.G.C., V.A., N.B. and A. C. E. carried out studies. L.G., A.C. and S.C. performed RNA-seq analysis. C.J., W.S.S., N.O., C.C and N.P. carried out metabolite and stable isotope tracing studies. J.G. helped with TEM and SEM studies. M.F. and E.S. performed histological analysis. J.A.H. and C.H. were involved in microbiome studies. H.D. performed fly studies. J.A.B., R.P.F. and N.S. contributed intellectually and N.S. helped with enteroid experiments. E.S. and C.J. wrote the manuscript with all the other authors’ feedback.

## Contact for Reagent and Resource Sharing

Further information and requests for reagents should be directed to and will be fulfilled by the Lead Contact, Ergun Sahin or Cholsoon Jang

## Experimental Model and Subject Details

### Mouse Studies

All animal experiments were performed according to procedures approved by the Institutional Animal Care and Use Committee at Baylor College of Medicine. Mice were maintained in openly ventilated cages with group housing, in a temperature-controlled (20-22°C) facility with 12 h light/12 h dark cycle, with *ad libitum* access to food (PicoLab® Select Rodent Diet 50 IF/6F Diet) and water. Mice were maintained on standard rodent chow with 12-hr light-dark cycles. For all studies age and sex matched mice on a C57/B6 background were used.

### Tert-deficient mice

TERT deficient mice were provided by Ronald A. DePinho and generated by deletion of exon 1, an essential part of the TERT open reading frame^1^. TERT heterozygous mice were continuously interbred to produce successive generations (G1-G3) of TERT deficient mice with progressive telomere shortening in each generation as described^2^. G3 mice were used for all studies and referred to as TKO mice.

### Glu 5 deficient mice

Glut 5 germline deficient mice3 were provided by Ronald Ferraris with permission from Jian Zuo.

### Lgr5-EGFP-IRES-CreERT2

Lgr5-EGFP-IRES-CreERT2 mice^4^ were obtained from Jackson Laboratory.

## Method Details

### RNA isolation, cDNA synthesis and Real-time PCR

Trizol reagent (Thermo Fisher Scientific) was used to extract total RNA from tissues. RNA was digested with DNase I (New England Biolabs) and purified with Rneasy Mini columns (QIAGEN) according to manufacturer’s instructions. 0.5 μg of total RNA was used for cDNA synthesis by reverse transcription reaction with Protoscript II reverse transcriptase (New England Biolabs). Quantitative PCR was performed using SensiFAST™ SYBR kit (BIOLINE). The primer sequences used for qPCR are listed in Table S1. ΔΔ CT method was used to calculate the expression levels of the transcripts.

### RNA sequencing

#### Small intestine

After euthanasia, intestines were longitudinally opened and washed with ice-cold PBS. Using glass slides, epithelial cells were scraped off in the first part of the small intestine (duodenum) and RNA was immediately isolated as described above using Trizol followed by DNAse digestion and column purification.

#### Colon

Approximately 3 cm of tissue distal to the coecum was selected for immediate RNA isolation by a combined Trizol/column purification procedure.

#### Library preparation

RNA processing, library preparation and sequencing was performed by Quick biology, Pasadena, CA. The RNA integrity was checked by Agilent Bioanalyzer 2100 or Tapestation 4200 (Agilent, Santa Clara, CA) and samples with clean rRNA peaks (RIN>6) were used for library preparation. RNA-Seq library was prepared according to KAPA mRNA HyperPrep kit with 200-300 bp insert size (Roche, Wilmington, MA) using 250 ng of total RNAs as input. Final library quality and quantity were analyzed by Agilent Bioanalyzer 2100 or Tapestation 4200 and Life Technologies Qubit 3.0 Fluorometer. 150 bp PE (paired-end) reads were sequenced on Illumina HiSeq sequencer (Illumina Inc., San Diego, CA).

#### RNA sequencing analysis

Quality of the RNAseq data was examined with FastQC v0.11.8. Sequences were then aligned to GRCm38 (mm10) mouse genome using Spliced Transcripts Alignment to a Reference (STAR v2.7.0) software and gene-level read counts were generated using the Subread package v1.6.3. Genes with expression of 0 in more than 30% of all samples were removed; counts were normalized using limma/voom^5^. Sample clustering was performed by Principal Component Analysis (PCA). We performed DE analysis using the R package limma v3.44.3^6^. BH adjusted P value ≤ 0.05 and fold change ≥ 2 were used as the thresholds for detecting DEGs. To understand what biological processes are represented in the DEGs, we performed functional enrichment of gene ontology (GO) and canonical functional pathway gene sets from the Molecular Signatures Database (MSigDB) gene annotation database v6.1^7,8^.

### Histology & Immunohistochemistry/Immunofluorescence

Small and large intestines were removed, washed with PBS, cut open and fixed in 4% paraformaldehyde overnight at 4°C. Tissues were embedded in paraffin, sectioned (5 μm), and stained with hematoxylin and eosin (H&E) for histopathological examination.

For immunohistochemistry, tissue sections were deparaffinized and rehydrated in an ethanol series. Antigen retrieval was performed using citrate buffer (Vector Labs, H-3300) in a pressure cooker for 30 minutes at 109°C. Tissues were then permeabilized with 0.5% Triton X-100/PBS for 1 hour at room temperature and quenched in 3% H_2_O_2_ for 20 minutes at room temperature. Tissue sections were blocked with 2.5% horse serum (Vector Labs, 30021) for 1 hour in a humidity chamber at room temperature and were then incubated at 37°C for 1 hour with primary antibodies diluted in 2.5% horse serum. The antibodies used included c-Myc (Santa Cruz, sc-788), beta-catenin (BD Biosciences, 610153), Ki67 (BD Biosciences, 550609), and villin (Santa Cruz, sc-7672). Tissues were then incubated with ImmPRESS Polymer Reagent (Vector Labs) for 30 min at room temperature in a humidity chamber. DAB Chromogen reagent (Biocare Medical, DB801L) was added onto slides until sufficient stain intensity was reached. All tissues were developed for the same time, followed by counterstaining with hematoxylin, dehydrated in an ethanol series, and mounted with Permount mounting medium (Fisher, SP15-100).

For immunofluorescence, formalin-fixed paraffin-embedded tissues were cut (5 μm) thick sections and slides were deparaffinized and rehydrated. Antigen retrieval with citrate buffer (Vector) was performed in a pressure cooker (30 minutes at 109°C). Autofluorescence was quenched by incubating tissue slides in 50mM NH4Cl/PBS for 30 minutes, followed by permeabilization in 0.5% Triton-X/PBS for 1 hour. Slides were blocked for 1 hour in 5% BSA/TBST in humidified chamber. Primary antibody incubation was carried out in humidified chamber at 37°C for 1 hour, followed by secondary antibody (Alexa Fluor, Thermo Fisher Scientific) for 1 hour at room temperature. Slides were counterstained in DAPI for 10 minutes and mounted in Fluoromount medium.

#### Determination of fructose levels in cecal content

WT, TKO and Glut 5 were gavaged with a mix of 1:1 universally labeled fructose (U-^13^C6, Cambridge Isotope Laboratories) and regular glucose at 1g/kg body weight. Fructose levels were determined two hours later by mass spectrometry using equal amounts of cecal content or as described before^9^.

#### Transmission Electron Microscopy (TEM)

Tissues were fixed in McDowell and Trump (4F:1G) fixative, pH 7.3, for overnight at 4°C. After fixation, samples were washed thoroughly in PBS. For dehydration, samples were sequentially put in 25%, 50%, 75%, 95% and 100% ethanol solution in water for 15 minutes each. Tissues were then infiltrated in a gradient series of Spurr’s Low Viscosity resin (Sigma-Aldrich) and ethanol and then embedded in fresh Spurr’s resin and polymerized for 3 days at 60°C. 55–60 nm thin sections were cut on a Leica (Wezlar, BRD) UC7 ultra-microtome and viewed on a Hitachi (Tokyo, Japan) H7500 transmission electron microscope set to 80 kV. Images were collected using an AMT XR-16 digital camera and AMT Image Capture (Advanced Microscopy Techniques Corp, Woburn, MA), v602.600.51 software.

Microvilli length was determined in 3 independent mice with 60 microvilli per mouse as described^10^.

#### Scanning Electron Microscopy (SEM)

Tissues were fixed in McDowell and Trump (4F:1G) fixative, pH 7.3, for overnight at 4°C. After fixation, samples were washed thoroughly with PBS and dehydrated sequentially in 20%, 40%, 60%, 90% and 100% ethanol. Samples were then put in a critical point dryer (Autosamdri-815 from Tousimis Research Corporation) in ethanol for 1.5 hours and afterwards applied onto a silicon wafer (Ted Pella Inc.) and left to dry at room temperature. Double side Carbon conductive tape (Cat# 16084-7, Ted Pella Inc) was used to immobilize the wafer on an aluminum SEM sample holder. To enhance the SEM image contract, samples were coated with a 5nm thin iridium film using a magnetron sputtering Coater (208HR High Resolution Sputter Coater, Ted Pella Inc.). Images were captured using the Nava Nano SEM 230 system (FEI). The work distance was at 5 mm. All images were captured at room temperature and in a high vacuum (2E^−6^ Torr). In the process of imaging, the e-beam diameter was set at 8 nm and the acceleration high voltage was set at 5KV.

#### Fructose and Glucose diet

WT and TKO mice had free access either to plain tap water, water enriched with 30% Glucose (Sigma) or 30% fructose (Sigma) for indicated durations. Sugar-enriched water was changed twice per week.

#### CLAMS analysis

Comprehensive Lab Animal Monitoring System (CLAMS, Columbus Instruments) was used to measure water, food intake and energy expenditure in WT and TKO mice as described before^11^.

#### Analysis of intestinal permeability

Intestinal permeability was determined in mice that were starved overnight and gavaged with Fluorescein isothiocynate conjugated dextran (FITC-dextran, Sigma Aldrich, FD4) at a concentration of 100mg/ml. FITC concentration in plasma was determined with a fluorescent spectrophotometer (Thermo Fisher) 4 hours post-gavage using serially diluted FITC-dextran as a standard. The background levels were determined in plasma from mice that were not gavaged with FITC-dextran and subtracted from individual values.

#### Determination of albumin concentration in mice feces

Albumin concentration in stool was determined using a commercial kit (Albumin ELISA Kit, Genway, GWB-282C17) according to the manufacturer’s instructions. 100 mg of freshly collected stool was sonicated in PBS, centrifuged and the supernatant was collected and diluted 1:500 with dilution buffer. The diluted samples were used for the ELISA following the manufacturer’s instructions and the albumin concentration was determined from triplicate wells. Blank wells were used to determine the background and an Albumin standard dilution series was used to calculate the concentration.

#### Quantification of pro-inflammatory cytokines by ELISA in the colon

Cytokine levels were determined using commercially available enzyme-linked immunosorbent assay (ELISA) kit (Quantikine Murine; R&D Systems). Absorbance values from each ELISA is normalized using a Bradford protein assay respective to each sample and is expressed in units of pg/mg of protein.

#### Histological Colitis Score

Formalin-fixed and paraffin-embedded intestinal tissue was sectioned and stained with hematoxylin and eosin (H&E). A modified semi-quantitative scoring system that assigned degree of inflammation, epithelial changes and mucosal architecture alterations score values from 0 to 5 was used to assess colitis severity^12^. Inflammation (scores 0-5; 0, no inflammation; 1, minimal, restricted to mucosa; 2, mild, occasional submucosa; 3, moderate, mucosa and submucosa involved; 4, marked, mucosa and submucosa; 5, marked, transmural), epithelial changes (each assigned subcategory assigned 0 (absent) or 1 (present); Hyperplasia, Goblet cell loss, Cryptitis, Crypt abscess, Erosion) and Mucosal Architecture (0 or 1 for each subcategory: ulceration, irregular crypts, crypt loss, granulation tissue)

#### Antibiotic treatment

To reduce bacterial load in colon a combination of 4 different antibiotics including ampicillin (1 g/L), vancomycin (500 mg/L), neomycin sulfate (1 g/L), and metronidazole (1 g/L) in drinking water as previously described^13^. Antibiotic-containing water was changed twice a week. Reduction of bacterial load was confirmed by 16S. Antibiotics were started 2 weeks prior to fructose treatment and continued throughout the diet duration.

### Metabolomic Analysis of colonic tissue

Metabolites were extracted from colonic tissue using previously described^14,15^. Briefly the extracted samples analyzed using high performance liquid chromatography (HPLC) coupled to Agilent 6490 QQQ mass spectrometry using ESI positive and negative ionization. Column was waters X-bridge amide 3.5 μm, 4.6 × 100 mm (Waters <u>Milford, MA</u>). In ESI positive, mobile phase A and B were 0.1% formic acid in water and acetonitrile respectively. In ESI negative, mobile phase A and B were 20 mM ammonium acetate in 95% acetonitrile and 5% water (pH 9.0) and 100% acetonitrile respectively. Gradient:0 – 3 min-85% B, 3–12 min-30% B, 12–15 min 2% B, 15–16 min 85% B. The data was processed using Mass Hunter Quantitate sofetware. The data was normalized with internal standard and log2-transformed per-sample, per-method basis. For every metabolite in the normalized dataset, two sample t-tests were conducted to compare expression levels between different groups. Differential metabolites were identified by adjusting the p-values for multiple testing at an FDR threshold of <0.25 and generated a heat map.

### Metabolite enrichment analysis (MSEA)

MSEA was performed according to Xia et al.^16^ using the webtool data base MetaboAnlyst (https://www.metaboanalyst.ca/). Over Representation Analysis was performed to identify enriched metabolites.

### Stable Isotope Tracing

#### Mice

TKO and WT mice that had been on a fructose diet for 2 months were provided universally labeled fructose dissolved in drinking water at 15% for two days after which fecal content and colon were subjected to MS analysis (see below).

##### Drosophila melanogaster

Fruit flies were provided 30% either unlabeled or universally labeled fructose (dissolved in water) through capillaries for 3 days after which head and body were separated and subjected separately to MS analysis.

#### Metabolomic

Serum (5 μl) was mixed with 150 μl 4°C 40:40:20 methanol:acetonitrile:water (extraction solvent), vortexed and immediately centrifuged at 16,000g for 10 minutes at 4 °C. The supernatant (~ 100 μl) was collected for liquid chromatography-mass spectrometry (LC-MS) analysis. Frozen feces, mouse tissues or fly samples were ground at liquid nitrogen temperature with a CryoMill (Retsch). The resulting tissue powder was weighed and then extracted by adding 4°C extraction solvent, vortexed and centrifuged at 16,000g for 10 minutes at 4°C. The volume of the extraction solution (μl) was 40x the weight of tissue (mg) to make an extract of 25 mg of tissue per ml of solvent. The supernatant (40 μl) was collected for LC-MS analysis. For metabolite measurements using LC-MS, a quadrupole orbitrap mass spectrometer (Q Exactive Plus; Thermo Fisher Scientific) operating in negative or positive ion mode was coupled to a Vanquish UHPLC system (Thermo Fisher Scientific) with electrospray ionization and used to scan from m/z 70 to 1,000 at 1 Hz, with a 140,000 resolution. LC separation was achieved on an XBridge BEH Amide column (2.1 × 150 mm^2^, 2.5 μm particle size, 130 Å pore size; Waters Corporation) using a gradient of solvent A (95:5 water: acetonitrile with 20 mM of ammonium acetate and 20 mM of ammonium hydroxide, pH 9.45) and solvent B (acetonitrile). The flow rate was 150 μl/min. The LC gradient was: 0 min, 85% B; 2 min, 85% B; 3 min, 80% B; 5 min, 80% B; 6 min, 75% B; 7 min, 75% B; 8 min, 70% B; 9 min, 70% B; 10 min, 50% B; 12 min, 50% B; 13 min, 25% B; 16 min, 25% B; 18 min, 0% B; 23 min, 0% B; 24 min, 85% B; and 30 min, 85% B. The autosampler temperature was 5°C and the injection volume was 3 μl. Data were analyzed using the MAVEN software (build 682, http://maven.princeton.edu/index.php). Natural isotope correction was performed with AccuCor R code (https://github.com/lparsons/accucor) as described previously by us^9^.

#### Lgr5 (+) ISC analysis

Lgr5 (+) cells were isolated as described before^17^. Lgr5 (+) high cells were sorted and analyzed.

#### Nicotinamide riboside (NR) treatment

NR (ChromaDex, 00014315) was mixed into chow at a concentration of 400mg/kg by a commercial provider (Research Diets Inc.) and mice were fed either chow or NR-enriched chow for 8 weeks prior to irradiation and clonogenic assay (see below).

#### Enteroid Studies and Clonogenic Assay

*Intestinal crypt isolation and culture* Crypts were isolated and cultured according to established procedures^18^. Briefly, a 15 cm segment of the proximal intestine was dissected and flushed with 10 mL of sterile cold PBS using a curved tip syringe. The segment was cut open longitudinally and swirled in sterile PBS to wash off remaining chyme and mucus. Next, the segment was cut into 0.5 cm pieces and transferred to a 15mL conical tube filled with 10mL of sterile PBS (without Ca/Mg). The pieces were pipetted up and down 15 times using a 10 mL serological pipette, the PBS was replaced, and the washing was repeated three times. Cleaned pieces were transferred to 10mL of 1mM EDTA/PBS (without Ca/Mg) and rocked at 4°C for 30 minutes. After chelation with EDTA, tissue pieces were moved to a new tube containing 5mL of shaking buffer (2g sorbitol, 3g sucrose per 50mL of sterile PBS). In shaking buffer, the pieces were pipetted up and down 10 times with a serological pipette to dissociate crypts. To further purify the suspension, villi were filtered using a 70mM mesh (Elko, 03-70/33) and the remaining crypts pelleted at 150g for 10 minutes. Pelleted crypts were resuspended in 100-200μL crypt media (Advanced DMEM/F12 supplemented with 1x Glutamax, 10mM HEPES, 10% R-spondin conditioned media, 10% Noggin conditioned media, 50ng/mL EGF, 10mM nicotinamide, 1x B27, and 1x N2) and a 10μL drop was used to count the number of crypts. Volumes were adjusted to equalize the concentration of crypts across samples. Crypts were plated in 48-well plates at a density of 40-60 crypts in 20μL of Matrigel with or without NR (2mM). Plates were incubated at 37°C for 15 minutes to allow the Matrigel to polymerize and subsequently filled with 200μL of crypt medium with and without NR (2mM).

*Enteroid Survival and Size* were analyzed as described before^18,19^. Briefly, growth was followed using a Nikon Eclipse TE300 microscope equipped with a motorized stage (Prior Scientific) that allows tracking by saving sample coordinates. Pictures were taken immediately after seeding and again at day 3 to assess survival. Survival was defined as crypts showing defined borders and growth. Additionally, budding was indicative of survival whereas a conglomeration of cells without defined lumen or borders indicated a dead crypt. Plotted counts represent the average percentage of viable organoids per mouse. Per well, 3 fields were analyzed at a minimum of two wells per mouse.

*Clonogenic Assay* TKO and WT mice were fed either a regular chow or NR-enriched chow for 8 weeks. Mice were then irradiated with 12Gy and sacrificed after 4 days for immunohistological analysis^20^. For every mouse, 8-10 images were taken of the proximal intestine starting at the proximal duodenum using Brunner’s glands as reference and of the colon starting at the distal colon using the anus as reference. Pictures were taken using a Nikon A1R-s confocal microscope and Nikon Elements software. Images of the intestine were taken using a 10x Plan Fluor 0.3 NA objective, while a 20x Plan Fluor Apo 0.75NA objective was used for colon pictures. A total of 5-8 WT and TKO mice were analyzed in two independent set of experiments. Ki67 staining (Abcam, ab15580) was performed and crypts containing 5 or more Ki67 positive cells were counted as viable. Counts represent the number of viable crypts per image field.

## Quantification and Statistical Analysis

### Statistics

Statistical analysis was performed with GraphPad Prism (GraphPad Software, La Jolla, CA, USA). Results are presented as mean values ± s.e.m. Student’s t-test was used to determine the statistical differences between two groups. P-values less than 0.05 were considered statistically significant. Statistical parameters can be found in the figure legends. The exact n-values and statistical significance are reported in Figure Legends.

### KEY RESOURCES TABLE

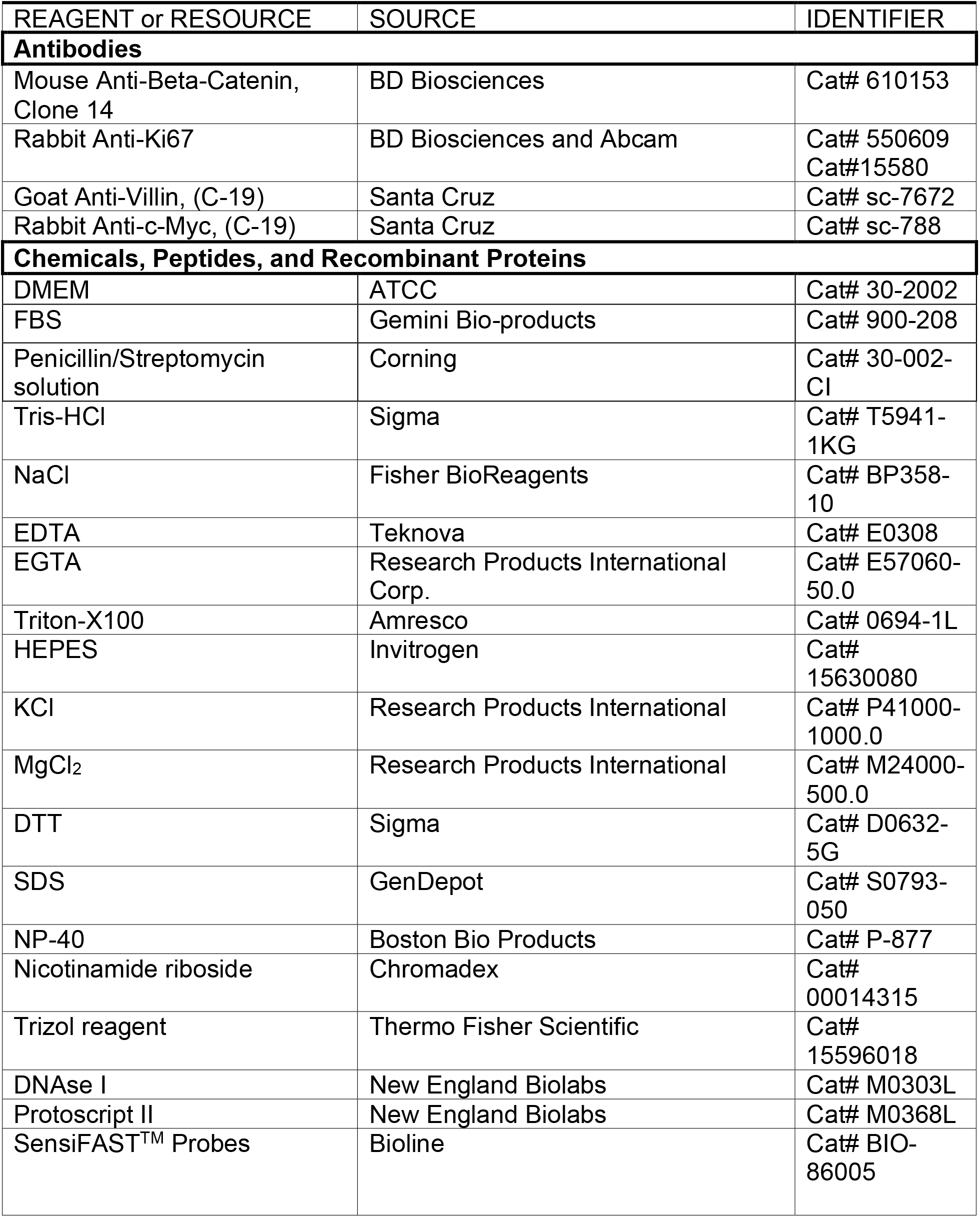
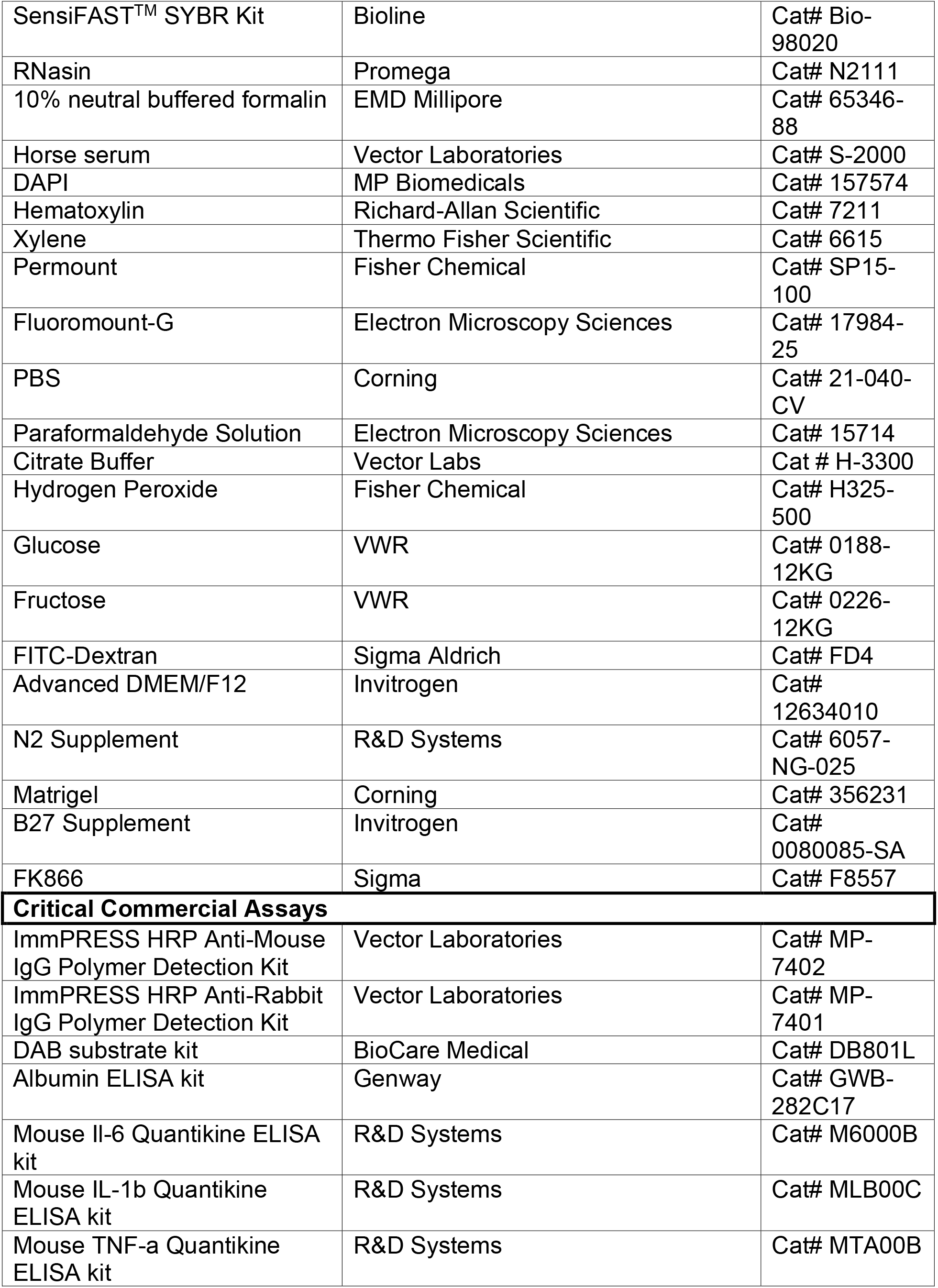
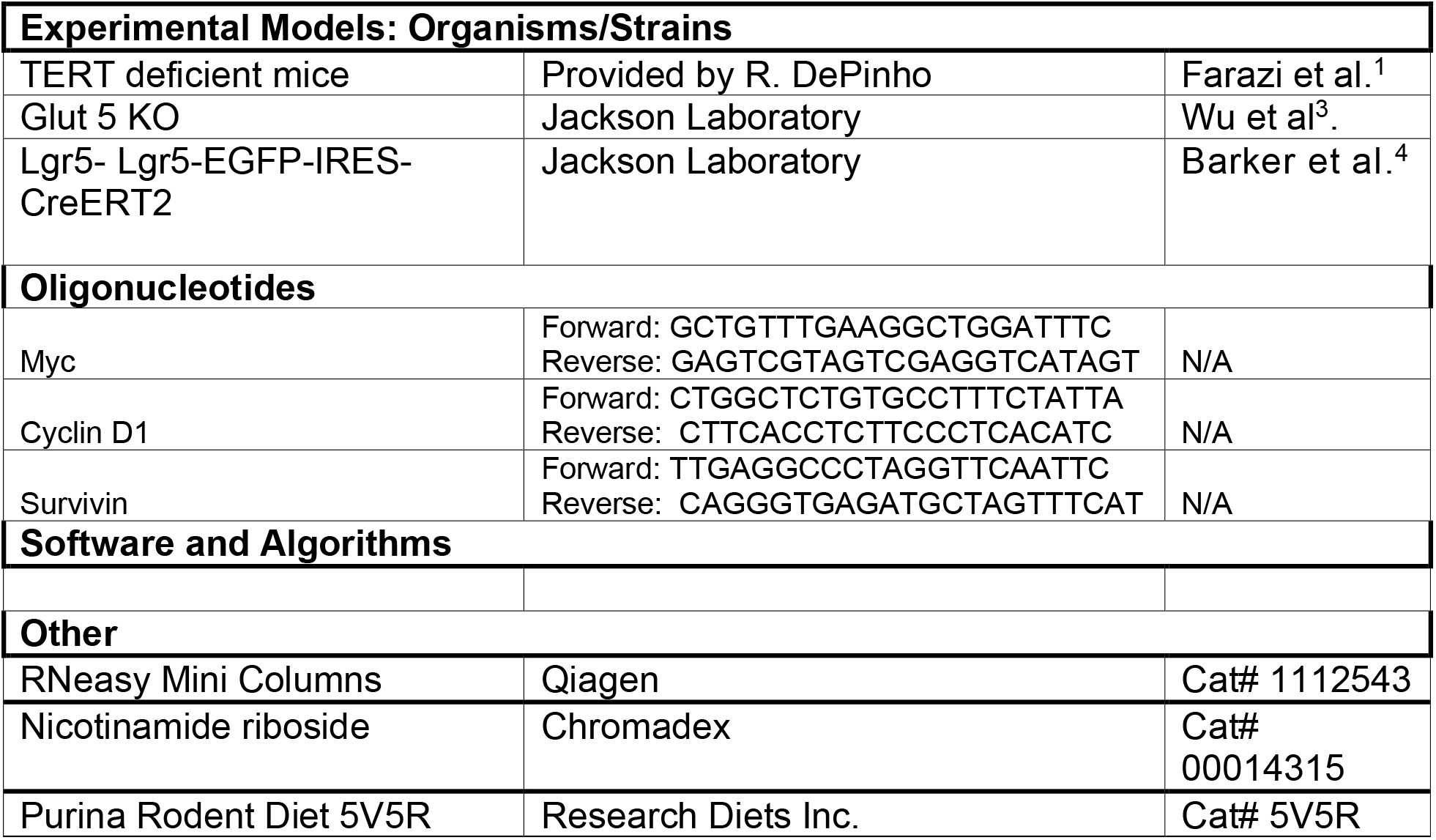

## References

1 Farazi, P. A., Glickman, J., Horner, J. & Depinho, R. A. Cooperative interactions of p53 mutation, telomere dysfunction, and chronic liver damage in hepatocellular carcinoma progression. Cancer Res 66, 4766–4773, doi:10.1158/0008-5472.CAN-05-4608 (2006).

2 Sahin, E. et al. Telomere dysfunction induces metabolic and mitochondrial compromise. Nature 470, 359–365, doi:nature09787 [pii] 10.1038/nature09787 (2011).

3 Wu, X. et al. Glucose transporter 5 is undetectable in outer hair cells and does not contribute to cochlear amplification. Brain Res 1210, 20–28, doi:10.1016/j.brainres.2008.02.094 (2008).

4 Barker, N. et al. Identification of stem cells in small intestine and colon by marker gene Lgr5. Nature 449, 1003–1007, doi:10.1038/nature06196 (2007).

5 Law, C. W., Chen, Y., Shi, W. & Smyth, G. K. voom: precision weights unlock linear model analysis tools for RNA-seq read counts. Genome Biology 15, R29, doi:10.1186/gb-2014-15-2-r29 (2014).

6 Ritchie, M. E. et al. limma powers differential expression analyses for RNA-sequencing and microarray studies. Nucleic Acids Res 43, e47, doi:10.1093/nar/gkv007 (2015).

7 Liberzon, A. et al. Molecular signatures database (MSigDB) 3.0. Bioinformatics (Oxford, England) 27, 1739–1740, doi:10.1093/bioinformatics/btr260 (2011).

8 Subramanian, A. et al. Gene set enrichment analysis: A knowledge-based approach for interpreting genome-wide expression profiles. Proceedings of the National Academy of Sciences 102, 15545, doi:10.1073/pnas.0506580102 (2005).

9 Jang, C. et al. The small intestine shields the liver from fructose-induced steatosis. Nat Metab 2, 586–593, doi:10.1038/s42255-020-0222-9 (2020).

10 Engevik, A. C. et al. Editing Myosin VB Gene to Create Porcine Model of Microvillus Inclusion Disease, With Microvillus-Lined Inclusions and Alterations in Sodium Transporters. Gastroenterology 158, 2236–2249 e2239, doi:10.1053/j.gastro.2020.02.034 (2020).

11 Lee, C. S. et al. A chemical chaperone improves muscle function in mice with a RyR1 mutation. Nat Commun 8, 14659, doi:10.1038/ncomms14659 (2017).

12 Erben, U. et al. A guide to histomorphological evaluation of intestinal inflammation in mouse models. Int J Clin Exp Pathol 7, 4557–4576 (2014).

13 Rakoff-Nahoum, S., Paglino, J., Eslami-Varzaneh, F., Edberg, S. & Medzhitov, R. Recognition of commensal microflora by toll-like receptors is required for intestinal homeostasis. Cell 118, 229–241, doi:10.1016/j.cell.2004.07.002 (2004).

14 Amara, C. S. et al. Serum Metabolic Profiling Identified a Distinct Metabolic Signature in Bladder Cancer Smokers: A Key Metabolic Enzyme Associated with Patient Survival. Cancer Epidemiol Biomarkers Prev 28, 770–781, doi:10.1158/1055-9965.EPI-18-0936 (2019).

15 Vantaku, V. et al. Epigenetic loss of AOX1 expression via EZH2 leads to metabolic deregulations and promotes bladder cancer progression. Oncogene 39, 6265–6285, doi:10.1038/s41388-019-0902-7 (2020).

16 Xia, J. & Wishart, D. S. MSEA: a web-based tool to identify biologically meaningful patterns in quantitative metabolomic data. Nucleic Acids Res 38, W71–77, doi:10.1093/nar/gkq329 (2010).

17 Sato, T. et al. Single Lgr5 stem cells build crypt-villus structures in vitro without a mesenchymal niche. Nature 459, 262–265, doi:10.1038/nature07935 (2009).

18 Mahe, M. M. et al. Establishment of Gastrointestinal Epithelial Organoids. Curr Protoc Mouse Biol 3, 217–240, doi:10.1002/9780470942390.mo130179 (2013).

19 Lo, Y. H. et al. SPDEF Induces Quiescence of Colorectal Cancer Cells by Changing the Transcriptional Targets of beta-catenin. Gastroenterology 153, 205–218 e208, doi:10.1053/j.gastro.2017.03.048 (2017).

20 Metcalfe, C., Kljavin, N. M., Ybarra, R. & de Sauvage, F. J. Lgr5+ stem cells are indispensable for radiation-induced intestinal regeneration. Cell Stem Cell 14, 149–159, doi:10.1016/j.stem.2013.11.008 (2014).

## References

1 Beumer, J. & Clevers, H. Cell fate specification and differentiation in the adult mammalian intestine. Nat Rev Mol Cell Biol 22, 39–53, doi:10.1038/s41580-020-0278-0 (2021).

2 Sommer, F. & Backhed, F. The gut microbiota--masters of host development and physiology. Nat Rev Microbiol 11, 227–238, doi:10.1038/nrmicro2974 (2013).

3 Turnbaugh, P. J. et al. An obesity-associated gut microbiome with increased capacity for energy harvest. Nature 444, 1027–1031, doi:10.1038/nature05414 (2006).

4 Barratt, M. J., Lebrilla, C., Shapiro, H. Y. & Gordon, J. I. The Gut Microbiota, Food Science, and Human Nutrition: A Timely Marriage. Cell Host Microbe 22, 134–141, doi:10.1016/j.chom.2017.07.006 (2017).

5 Odenwald, M. A. & Turner, J. R. The intestinal epithelial barrier: a therapeutic target? Nat Rev Gastroenterol Hepatol 14, 9–21, doi:10.1038/nrgastro.2016.169 (2017).

6 May, G. R., Sutherland, L. R. & Meddings, J. B. Is small intestinal permeability really increased in relatives of patients with Crohn’s disease? Gastroenterology 104, 1627–1632, doi:10.1016/0016-5085(93)90638-s (1993).

7 Hollander, D. et al. Increased intestinal permeability in patients with Crohn’s disease and their relatives. A possible etiologic factor. Ann Intern Med 105, 883–885, doi:10.7326/0003-4819-105-6-883 (1986).

8 Wyatt, J., Vogelsang, H., Hubl, W., Waldhoer, T. & Lochs, H. Intestinal permeability and the prediction of relapse in Crohn’s disease. Lancet 341, 1437–1439, doi:10.1016/0140-6736(93)90882-h (1993).

9 Arnott, I. D., Kingstone, K. & Ghosh, S. Abnormal intestinal permeability predicts relapse in inactive Crohn disease. Scand J Gastroenterol 35, 1163–1169, doi:10.1080/003655200750056637 (2000).

10 Matysiak-Budnik, T. et al. Alterations of the intestinal transport and processing of gliadin peptides in celiac disease. Gastroenterology 125, 696–707, doi:10.1016/s0016-5085(03)01049-7 (2003).

11 Konig, J. et al. Human Intestinal Barrier Function in Health and Disease. Clin Transl Gastroenterol 7, e196, doi:10.1038/ctg.2016.54 (2016).

12 Rea, I. M. et al. Age and Age-Related Diseases: Role of Inflammation Triggers and Cytokines. Front Immunol 9, 586, doi:10.3389/fimmu.2018.00586 (2018).

13 Furman, D. et al. Chronic inflammation in the etiology of disease across the life span. Nat Med 25, 1822–1832, doi:10.1038/s41591-019-0675-0 (2019).

14 Balestrieri, P. et al. Nutritional Aspects in Inflammatory Bowel Diseases. Nutrients 12, doi:10.3390/nu12020372 (2020).

15 Untersmayr, E., Brandt, A., Koidl, L. & Bergheim, I. The Intestinal Barrier Dysfunction as Driving Factor of Inflammaging. Nutrients 14, doi:10.3390/nu14050949 (2022).

16 Snoeck, V., Goddeeris, B. & Cox, E. The role of enterocytes in the intestinal barrier function and antigen uptake. Microbes Infect 7, 997–1004, doi:10.1016/j.micinf.2005.04.003 (2005).

17 Crawley, S. W., Mooseker, M. S. & Tyska, M. J. Shaping the intestinal brush border. J Cell Biol 207, 441–451, doi:10.1083/jcb.201407015 (2014).

18 Shifrin, D. A., Jr. et al. Enterocyte microvillus-derived vesicles detoxify bacterial products and regulate epithelial-microbial interactions. Curr Biol 22, 627–631, doi:10.1016/j.cub.2012.02.022 (2012).

19 Selsted, M. E. & Ouellette, A. J. Mammalian defensins in the antimicrobial immune response. Nat Immunol 6, 551–557, doi:10.1038/ni1206 (2005).

20 Giepmans, B. N. & van Ijzendoorn, S. C. Epithelial cell-cell junctions and plasma membrane domains. Biochim Biophys Acta 1788, 820–831, doi:10.1016/j.bbamem.2008.07.015 (2009).

21 Marchiando, A. M., Graham, W. V. & Turner, J. R. Epithelial barriers in homeostasis and disease. Annu Rev Pathol 5, 119–144, doi:10.1146/annurev.pathol.4.110807.092135 (2010).

22 Muller, T. et al. MYO5B mutations cause microvillus inclusion disease and disrupt epithelial cell polarity. Nat Genet 40, 1163–1165, doi:10.1038/ng.225 (2008).

23 Wiegerinck, C. L. et al. Loss of syntaxin 3 causes variant microvillus inclusion disease. Gastroenterology 147, 65–68 e10, doi:10.1053/j.gastro.2014.04.002 (2014).

24 Schmitz, H. et al. Altered tight junction structure contributes to the impaired epithelial barrier function in ulcerative colitis. Gastroenterology 116, 301–309, doi:10.1016/s0016-5085(99)70126-5 (1999).

25 Prasad, S. et al. Inflammatory processes have differential effects on claudins 2, 3 and 4 in colonic epithelial cells. Laboratory investigation; a journal of technical methods and pathology 85, 1139–1162, doi:10.1038/labinvest.3700316 (2005).

26 Hollander, D. Crohn’s disease--a permeability disorder of the tight junction? Gut 29, 1621–1624, doi:10.1136/gut.29.12.1621 (1988).

27 Pearson, A. D., Eastham, E. J., Laker, M. F., Craft, A. W. & Nelson, R. Intestinal permeability in children with Crohn’s disease and coeliac disease. Br Med J (Clin Res Ed) 285, 20–21, doi:10.1136/bmj.285.6334.20 (1982).

28 Ukabam, S. O., Clamp, J. R. & Cooper, B. T. Abnormal small intestinal permeability to sugars in patients with Crohn’s disease of the terminal ileum and colon. Digestion 27, 70–74, doi:10.1159/000198932 (1983).

29 Cesar Machado, M. C. & da Silva, F. P. Intestinal Barrier Dysfunction in Human Pathology and Aging. Curr Pharm Des 22, 4645–4650, doi:10.2174/1381612822666160510125331 (2016).

30 Britton, E. & McLaughlin, J. T. Ageing and the gut. Proc Nutr Soc 72, 173–177, doi:10.1017/S0029665112002807 (2013).

31 Funk, M. C., Zhou, J. & Boutros, M. Ageing, metabolism and the intestine. EMBO Rep 21, e50047, doi:10.15252/embr.202050047 (2020).

32 VanDussen, K. L. et al. Abnormal Small Intestinal Epithelial Microvilli in Patients With Crohn’s Disease. Gastroenterology 155, 815–828, doi:10.1053/j.gastro.2018.05.028 (2018).

33 Savage, S. A. & Bertuch, A. A. The genetics and clinical manifestations of telomere biology disorders. Genet Med 12, 753–764, doi:10.1097/GIM.0b013e3181f415b5 (2010).

34 Armanios, M. Telomeres and age-related disease: how telomere biology informs clinical paradigms. J Clin Invest 123, 996–1002, doi:10.1172/JCI66370 (2013).

35 Vulliamy, T. et al. The RNA component of telomerase is mutated in autosomal dominant dyskeratosis congenita. Nature 413, 432–435, doi:10.1038/35096585 (2001).

36 Jonassaint, N. L., Guo, N., Califano, J. A., Montgomery, E. A. & Armanios, M. The gastrointestinal manifestations of telomere-mediated disease. Aging Cell 12, 319–323, doi:10.1111/acel.12041 (2013).

37 O’Sullivan, J. N. et al. Chromosomal instability in ulcerative colitis is related to telomere shortening. Nat Genet 32, 280–284, doi:10.1038/ng989 (2002).

38 Risques, R. A. et al. Ulcerative colitis is a disease of accelerated colon aging: evidence from telomere attrition and DNA damage. Gastroenterology 135, 410–418, doi:10.1053/j.gastro.2008.04.008 (2008).

39 Chakravarti, D. et al. Telomere dysfunction instigates inflammation in inflammatory bowel disease. Proc Natl Acad Sci U S A 118, doi:10.1073/pnas.2024853118 (2021).

40 Hastie, N. D. et al. Telomere reduction in human colorectal carcinoma and with ageing. Nature 346, 866–868, doi:10.1038/346866a0 (1990).

41 Chin, L. et al. p53 deficiency rescues the adverse effects of telomere loss and cooperates with telomere dysfunction to accelerate carcinogenesis. Cell 97, 527–538, doi:10.1016/s0092-8674(00)80762-x (1999).

42 Sperka, T. et al. Puma and p21 represent cooperating checkpoints limiting self-renewal and chromosomal instability of somatic stem cells in response to telomere dysfunction. Nat Cell Biol 14, 73–79, doi:10.1038/ncb2388 (2011).

43 Woo, D. H. et al. Enhancing a Wnt-Telomere Feedback Loop Restores Intestinal Stem Cell Function in a Human Organotypic Model of Dyskeratosis Congenita. Cell Stem Cell 19, 397–405, doi:10.1016/j.stem.2016.05.024 (2016).

44 Yang, T. B. et al. Mutual reinforcement between telomere capping and canonical Wnt signalling in the intestinal stem cell niche. Nat Commun 8, 14766, doi:10.1038/ncomms14766 (2017).

45 Chakravarti, D. et al. Telomere dysfunction activates YAP1 to drive tissue inflammation. Nat Commun 11, 4766, doi:10.1038/s41467-020-18420-w (2020).

46 Chen, J. et al. Hematopoietic lineage skewing and intestinal epithelia degeneration in aged mice with telomerase RNA component deletion. Exp Gerontol 72, 251–260, doi:10.1016/j.exger.2015.10.016 (2015).

47 Gao, N., White, P. & Kaestner, K. H. Establishment of intestinal identity and epithelial-mesenchymal signaling by Cdx2. Dev Cell 16, 588–599, doi:10.1016/j.devcel.2009.02.010 (2009).

48 Kawabata, K. et al. A highfructose diet induces epithelial barrier dysfunction and exacerbates the severity of dextran sulfate sodiuminduced colitis. Int J Mol Med 43, 1487–1496, doi:10.3892/ijmm.2018.4040 (2019).

49 Montrose, D. C. et al. Dietary Fructose Alters the Composition, Localization, and Metabolism of Gut Microbiota in Association With Worsening Colitis. Cell Mol Gastroenterol Hepatol 11, 525–550, doi:10.1016/j.jcmgh.2020.09.008 (2021).

50 Goncalves, M. D. et al. High-fructose corn syrup enhances intestinal tumor growth in mice. Science 363, 1345–1349, doi:10.1126/science.aat8515 (2019).

51 White, J. S. Straight talk about high-fructose corn syrup: what it is and what it ain’t. Am J Clin Nutr 88, 1716S–1721S, doi:10.3945/ajcn.2008.25825B (2008).

52 Barone, S. et al. Slc2a5 (Glut5) is essential for the absorption of fructose in the intestine and generation of fructose-induced hypertension. J Biol Chem 284, 5056–5066, doi:10.1074/jbc.M808128200 (2009).

53 Andres-Hernando, A. et al. Deletion of Fructokinase in the Liver or in the Intestine Reveals Differential Effects on Sugar-Induced Metabolic Dysfunction. Cell Metab 32, 117–127 e113, doi:10.1016/j.cmet.2020.05.012 (2020).

54 Ishimoto, T. et al. Opposing effects of fructokinase C and A isoforms on fructose-induced metabolic syndrome in mice. Proc Natl Acad Sci U S A 109, 4320–4325, doi:10.1073/pnas.1119908109 (2012).

55 Sellmann, C. et al. Diets rich in fructose, fat or fructose and fat alter intestinal barrier function and lead to the development of nonalcoholic fatty liver disease over time. J Nutr Biochem 26, 1183–1192, doi:10.1016/j.jnutbio.2015.05.011 (2015).

56 Volynets, V. et al. Intestinal Barrier Function and the Gut Microbiome Are Differentially Affected in Mice Fed a Western-Style Diet or Drinking Water Supplemented with Fructose. J Nutr 147, 770–780, doi:10.3945/jn.116.242859 (2017).

57 Zhao, S. et al. Dietary fructose feeds hepatic lipogenesis via microbiota-derived acetate. Nature 579, 586–591, doi:10.1038/s41586-020-2101-7 (2020).

58 Bergheim, I. et al. Antibiotics protect against fructose-induced hepatic lipid accumulation in mice: role of endotoxin. Journal of hepatology 48, 983–992, doi:10.1016/j.jhep.2008.01.035 (2008).

59 Di Luccia, B. et al. Rescue of Fructose-Induced Metabolic Syndrome by Antibiotics or Faecal Transplantation in a Rat Model of Obesity. PLoS One 10, e0134893, doi:10.1371/journal.pone.0134893 (2015).

60 Kennedy, E. A., King, K. Y. & Baldridge, M. T. Mouse Microbiota Models: Comparing Germ-Free Mice and Antibiotics Treatment as Tools for Modifying Gut Bacteria. Front Physiol 9, 1534, doi:10.3389/fphys.2018.01534 (2018).

61 Gazzaniga, F., Stebbins, R., Chang, S. Z., McPeek, M. A. & Brenner, C. Microbial NAD metabolism: lessons from comparative genomics. Microbiol Mol Biol Rev 73, 529–541, Table of Contents, doi:10.1128/MMBR.00042-08 (2009).

62 Jang, C., Chen, L. & Rabinowitz, J. D. Metabolomics and Isotope Tracing. Cell 173, 822–837, doi:10.1016/j.cell.2018.03.055 (2018).

63 Vries, R. G., Huch, M. & Clevers, H. Stem cells and cancer of the stomach and intestine. Mol Oncol 4, 373–384, doi:10.1016/j.molonc.2010.05.001 (2010).

64 Withers, H. R. & Elkind, M. M. Microcolony survival assay for cells of mouse intestinal mucosa exposed to radiation. Int J Radiat Biol Relat Stud Phys Chem Med 17, 261–267 (1970).

65 Canto, C. et al. The NAD(+) precursor nicotinamide riboside enhances oxidative metabolism and protects against high-fat diet-induced obesity. Cell Metab 15, 838–847, doi:10.1016/j.cmet.2012.04.022 (2012).

66 Tao, S. et al. Wnt activity and basal niche position sensitize intestinal stem and progenitor cells to DNA damage. EMBO J 34, 624–640, doi:10.15252/embj.201490700 (2015).

67 Merino, B., Fernandez-Diaz, C. M., Cozar-Castellano, I. & Perdomo, G. Intestinal Fructose and Glucose Metabolism in Health and Disease. Nutrients 12, doi:10.3390/nu12010094 (2019).

68 Hannou, S. A., Haslam, D. E., McKeown, N. M. & Herman, M. A. Fructose metabolism and metabolic disease. J Clin Invest 128, 545–555, doi:10.1172/JCI96702 (2018).

69 Missios, P. et al. Glucose substitution prolongs maintenance of energy homeostasis and lifespan of telomere dysfunctional mice. Nat Commun 5, 4924, doi:10.1038/ncomms5924 (2014).

70 Gerner, R. R. et al. NAD metabolism fuels human and mouse intestinal inflammation. Gut 67, 1813–1823, doi:10.1136/gutjnl-2017-314241 (2018).

71 Amano, H. et al. Telomere Dysfunction Induces Sirtuin Repression that Drives Telomere-Dependent Disease. Cell Metab 29, 1274–1290 e1279, doi:10.1016/j.cmet.2019.03.001 (2019).

72 Barker, N. et al. Crypt stem cells as the cells-of-origin of intestinal cancer. Nature 457, 608–611, doi:10.1038/nature07602 (2009).

73 Sahin, E. & Depinho, R. A. Linking functional decline of telomeres, mitochondria and stem cells during ageing. Nature 464, 520–528, doi:10.1038/nature08982 (2010).

74 Choudhury, A. R. et al. Cdkn1a deletion improves stem cell function and lifespan of mice with dysfunctional telomeres without accelerating cancer formation. Nat Genet 39, 99–105, doi:10.1038/ng1937 (2007).

75 Gu, B. W. et al. Impaired Telomere Maintenance and Decreased Canonical WNT Signaling but Normal Ribosome Biogenesis in Induced Pluripotent Stem Cells from X-Linked Dyskeratosis Congenita Patients. PLoS One 10, e0127414, doi:10.1371/journal.pone.0127414 (2015).

76 Munroe, M. et al. Telomere Dysfunction Activates p53 and Represses HNF4alpha Expression Leading to Impaired Human Hepatocyte Development and Function. Hepatology 72, 1412–1429, doi:10.1002/hep.31414 (2020).

77 Noah, T. K., Donahue, B. & Shroyer, N. F. Intestinal development and differentiation. Exp Cell Res 317, 2702–2710, doi:10.1016/j.yexcr.2011.09.006 (2011).

78 Srivillibhuthur, M. et al. TFAM is required for maturation of the fetal and adult intestinal epithelium. Dev Biol 439, 92–101, doi:10.1016/j.ydbio.2018.04.015 (2018).

79 Sahin, E. et al. Telomere dysfunction induces metabolic and mitochondrial compromise. Nature 470, 359–365, doi:nature09787 [pii] 10.1038/nature09787 (2011).

80 Cottliar, A. et al. Telomere length study in celiac disease. Am J Gastroenterol 98, 2727–2731, doi:10.1111/j.1572-0241.2003.08720.x (2003).

81 Hummel, S. et al. Telomere shortening in enterocytes of patients with uncontrolled acute intestinal graft-versus-host disease. Blood 126, 2518–2521, doi:10.1182/blood-2015-03-633289 (2015).

82 Hiyama, E. et al. Telomerase activity in human intestine. Int J Oncol 9, 453–458, doi:10.3892/ijo.9.3.453 (1996).

83 Dunne, M. R., Byrne, G., Chirdo, F. G. & Feighery, C. Coeliac Disease Pathogenesis: The Uncertainties of a Well-Known Immune Mediated Disorder. Front Immunol 11, 1374, doi:10.3389/fimmu.2020.01374 (2020).

84 Nalle, S. C. et al. Recipient NK cell inactivation and intestinal barrier loss are required for MHC-matched graft-versus-host disease. Sci Transl Med 6, 243ra287, doi:10.1126/scitranslmed.3008941 (2014).

85 Torrens-Mas, M. et al. Organoids: An Emerging Tool to Study Aging Signature across Human Tissues. Modeling Aging with Patient-Derived Organoids. Int J Mol Sci 22, doi:10.3390/ijms221910547 (2021).

86 Bilinski, C., Burleson, J. & Forouhar, F. Inflammation associated with neoplastic colonic polyps. Ann Clin Lab Sci 42, 266–270 (2012).

87 Claesson, M. J. et al. Gut microbiota composition correlates with diet and health in the elderly. Nature 488, 178–184, doi:10.1038/nature11319 (2012).

88 Sauvanet, C., Wayt, J., Pelaseyed, T. & Bretscher, A. Structure, regulation, and functional diversity of microvilli on the apical domain of epithelial cells. Annu Rev Cell Dev Biol 31, 593–621, doi:10.1146/annurev-cellbio-100814-125234 (2015).

89 Holohan, B., Wright, W. E. & Shay, J. W. Cell biology of disease: Telomeropathies: an emerging spectrum disorder. J Cell Biol 205, 289–299, doi:10.1083/jcb.201401012 (2014).

90 Savage, S. A. Human telomeres and telomere biology disorders. Prog Mol Biol Transl Sci 125, 41–66, doi:10.1016/B978-0-12-397898-1.00002-5 (2014).

