## Supplemental Figures for "Telomere dysfunction impairs intestinal differentiation and predisposes to diet-induced colitis"

**a**

| ACTIVE ION TRANSMEMBRANE | SYMPORTER ACTIVITY | ANION TRANSPORT | ACTIVE TRANSMEMBRANE TRANSPORTER ACTIVITY | SOLUTE CATION SYMPORTER | TRANS MEMBRANE TRANSPORT |
|---|---|---|---|---|---|
| Abcc3;Atp1a1;Atp1b1;Atp2a3;Atp2b1;Clcn5;Slc13a1;Slc13a2;Slc15a1;Slc20a1;Slc23a1;Slc26a11;Slc2a10;Slc2a8;Slc37a4;Slc4a11;Slc4a7;Slc5a1;Slc5a4a;Slc5a6;Slc6a19;Slc6a4;Slc6a6;Slc6a8 | Slc13a1;Slc13a2;Slc15a1;Slc16a5;Slc20a1;Slc23a1;Slc25a22;Slc2a10;Slc2a8;Slc4a11;Slc4a7;Slc5a1;Slc5a4a;Slc5a6;Slc6a19;Slc6a4;Slc6a6;Slc6a8 | Abat;Abcb1a;Abcc2;Abcc3;Clca4a;Clcn5;Cldn4;Clic1;Cln8;Cry2;Cyp4f18;Irs2;Ldlr;Lrrc8a;Per2;Plekha8;Plscr1;Pparg;Prkaa2;Prnp;Rxra;Slc13a1;Slc13a2;Slc16a5;Slc20a1;Slc23a1;Slc25a22;Slc26a11;Slc2a2;Slc2a8;Slc37a4;Slc3a1;Slc4a11;Slc4a7;Slc5a6;Slc6a19;Slc6a6;Slc6a8;Slc7a8;Slc9a3r1;Stxbp1;Ttyh3;Vamp2;Xkr9 | Abca8b;Abcb1a;Abcc2;Abcc3;Atp1a1;Atp1b1;Atp2a3;Atp2b1;Clcn5;Slc13a1;Slc13a2;Slc15a1;Slc16a5;Slc20a1;Slc23a1;Slc25a22;Slc26a11;Slc2a10;Slc2a8;Slc37a4;Slc4a11;Slc4a7;Slc5a1;Slc5a4a;Slc5a6;Slc6a19;Slc6a4;Slc6a6;Slc6a8 | Slc13a1;Slc13a2;Slc15a1;Slc20a1;Slc23a1;Slc2a10;Slc2a8;Slc4a7;Slc5a1;Slc5a4a;Slc5a6;Slc6a19;Slc6a4;Slc6a6;Slc6a8 | Abca8b;Abcb1a;Abcc2;Abcc3;Actb;Ank3;Anxa2;Atp1a1;Atp1b1;Atp2a3;Atp2b1;Azin1;Azin2;Bcr;Bin1;Chp1;Clca4a;Clcn5;Cldn4;Clic1;Cln8;Cnnm1;Gjb3;Hap1;Irs2;Itpr2;Klf15;Lrrc8a;Mcoln1;Nipal1;Otop2;Per2;Plcd1;Pm20d1;Prkaa2;Prnp;S100a6;Scnn1a;Sgk3;Slc13a1;Slc13a2;Slc15a1;Slc16a5;Slc20a1;Slc23a1;Slc25a22;Slc26a11;Slc2a10;Slc2a2;Slc2a5;Slc2a8;Slc30a2;Slc37a4;Slc3a1;Slc4a11;Slc4a7;Slc5a1;Slc5a4a;Slc5a6;Slc6a19;Slc6a4;Slc6a6;Slc6a8;Slc7a8;Slc9a3r1;Tmc7;Tmem144;Tmem63b;Trib3;Ttyh3;Ubc;Vamp2 |
| **CLUSTER OF ACTIN BASED CELL PROJECTIONS** | **BRUSH BORDER** | | | | |
| Abcc2;Clic1;Ezr;Mpp1;Myh10;Myo1a;Myo1c;Myo7b;Plec;Rhoc;Scin;Sis;Slc15a1;Slc2a2;Slc3a1;Slc4a7;Slc5a1;Slc5a6;Slc6a19;Slc9a3r | Abcc2;Clic1;Ezr;Myh10;Myo1a;Myo1c;Myo7b;Plec;Scin;Sis;Slc15a1;Slc2a2;Slc3a1;Slc5a1;Slc5a6;Slc6a19;Slc9a3r1 | | | | |

**sFig.1**

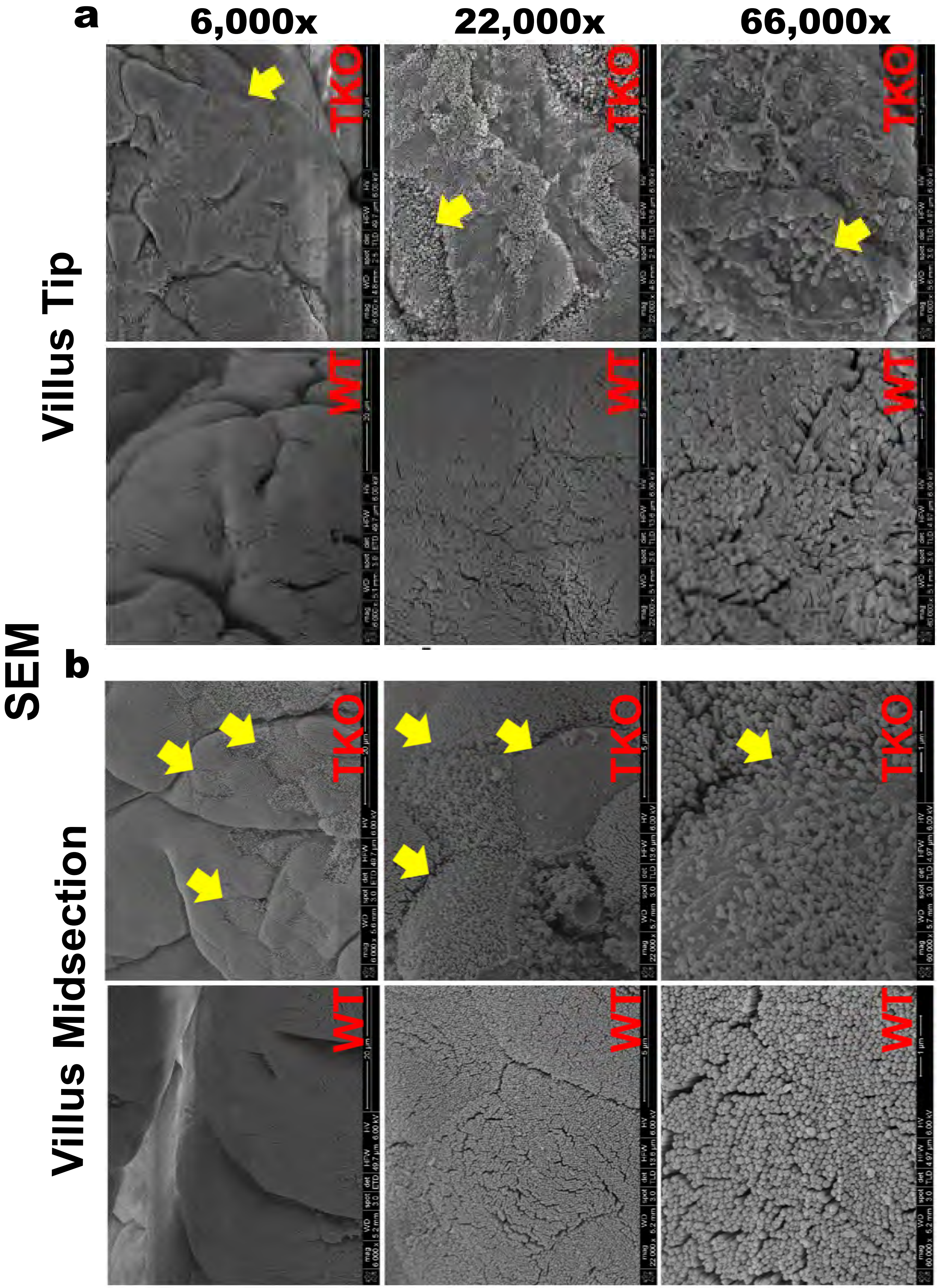
a
6,000x
22,000x
66,000x
Villus Tip
TKO
WT
SEM
b
Villus Midsection
TKO
WT

sFig.2

### COLON

c

SEM

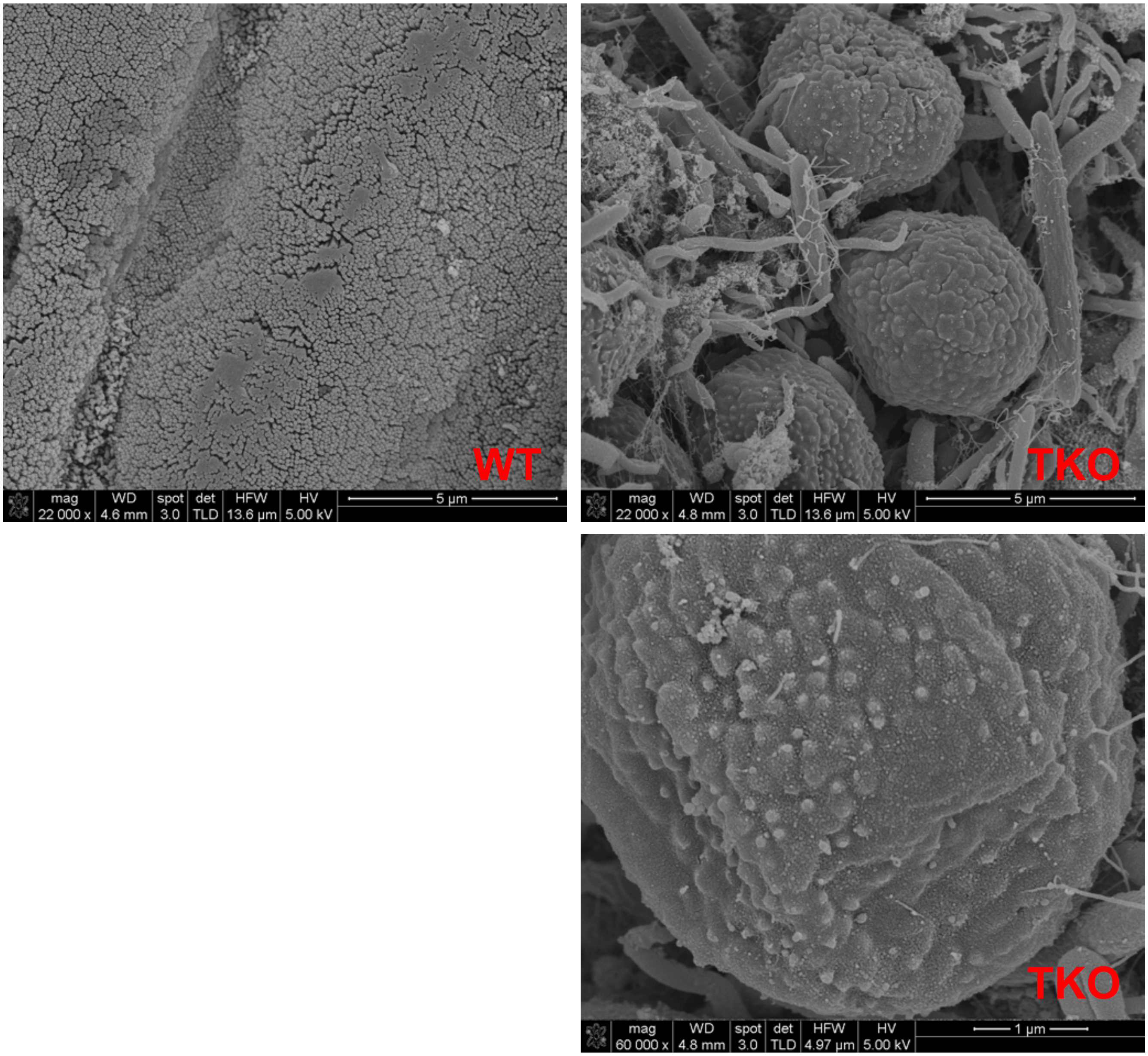


sFig.2

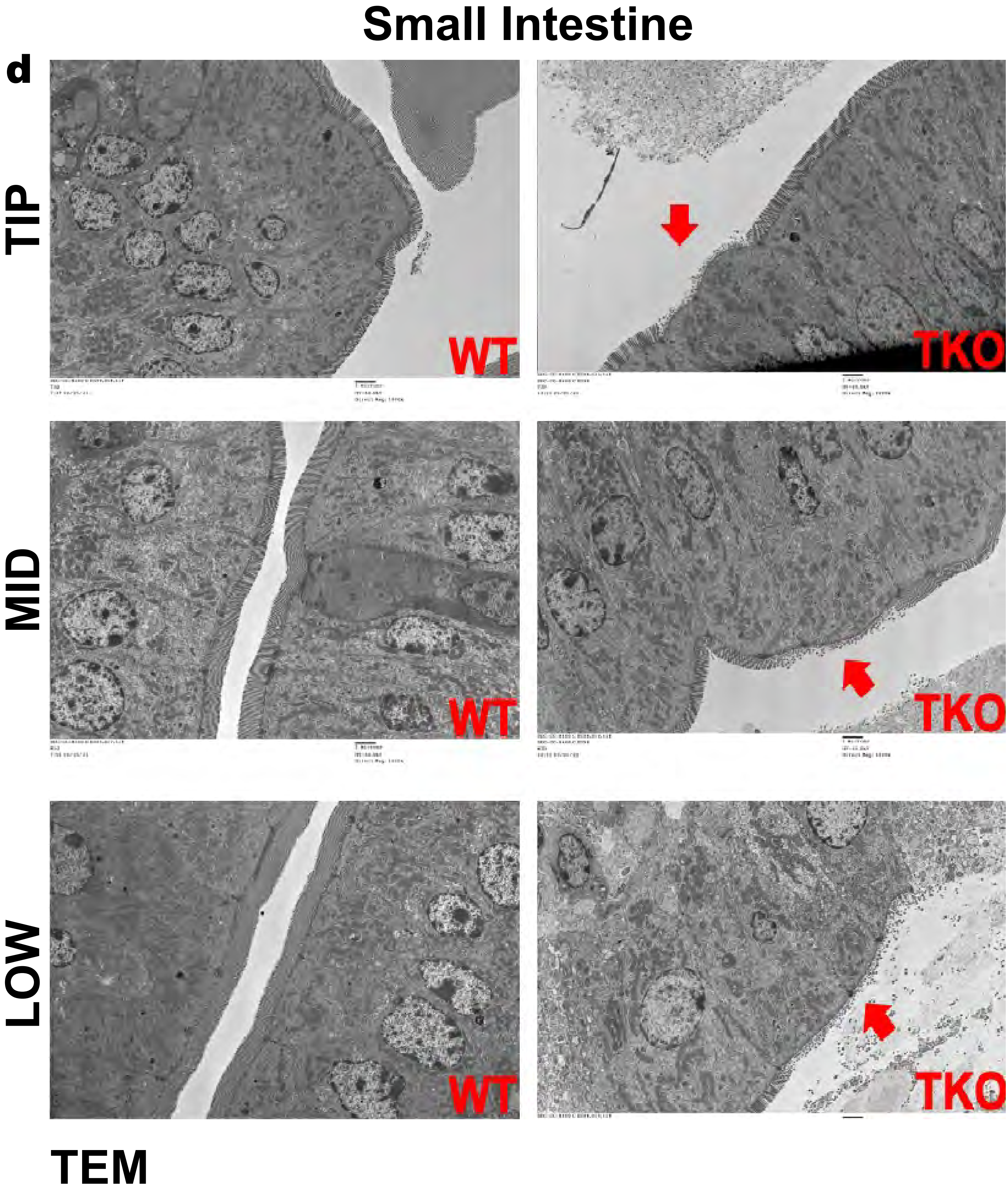


sFig.2

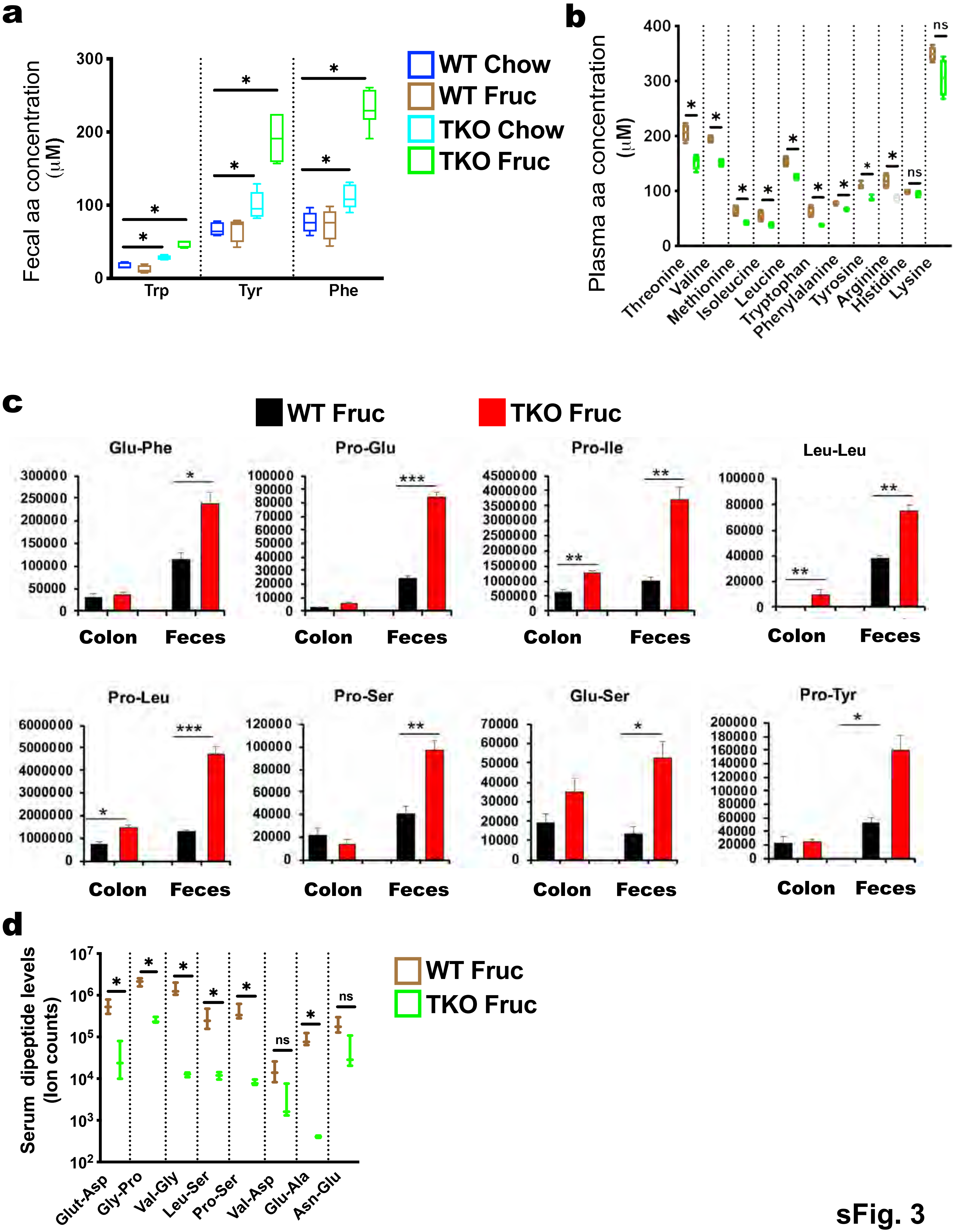
a
Fecal aa concentration (μM)
300
200
100
0
Trp
Tyr
Phe
WT Chow
WT Fruc
TKO Chow
TKO Fruc
b
Plasma aa concentration (μM)
400
300
200
100
0
ns
Threonine
Valine
Methionine
Isoleucine
Leucine
Tryptophan
Phenylalanine
Tyrosine
Arginine
Histidine
Lysine
c
WT Fruc
TKO Fruc
Glu-Phe
Pro-Glu
Pro-Ile
Leu-Leu
Pro-Leu
Pro-Ser
Glu-Ser
Pro-Tyr
Colon
Feces
d
Serum dipeptide levels (Ion counts)
Glut-Asp
Gly-Pro
Val-Gly
Leu-Ser
Pro-Ser
Val-Asp
Glu-Ala
Asn-Glu
WT Fruc
TKO Fruc

sFig. 3

**e**

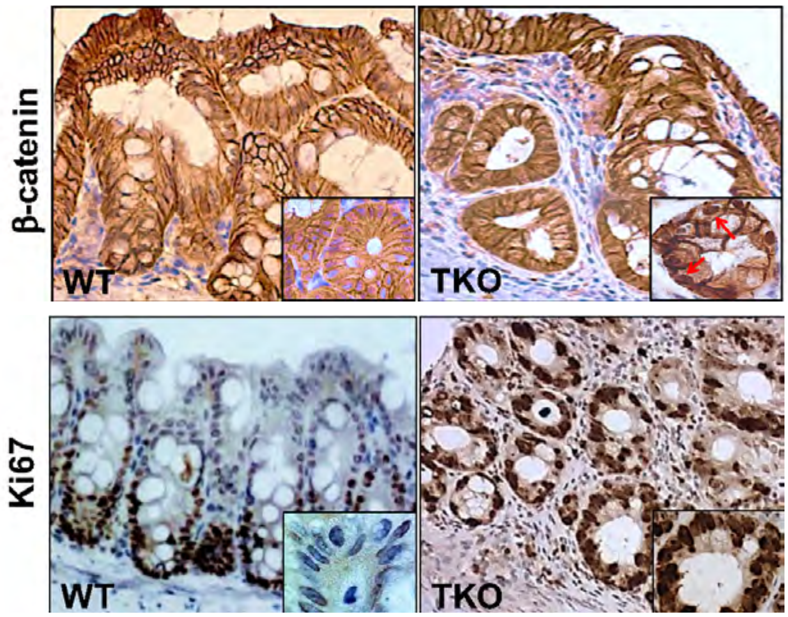


**f**

| Genotype | Diet Duration Chow +Fructose (months) | Number of mice analyzed | Age at which analyzed (months) | Mice with colitis | Percent of mice with colitis |
|---|---|---|---|---|---|
| WT | 0 | 20 | 2 | 0 | 0 |
| | 2 | 10 | 4 | 0 | 0 |
| | 3 | 10 | 5 | 0 | 0 |
| | 4 | 10 | 6 | 0 | 0 |
| | 6 | 8 | 8 | 0 | 0 |
| TKO | 0 | 20 | 2 | 0 | 0 |
| | 2 | 10 | 4 | 4 | 40 |
| | 3 | 12 | 5 | 8 | 66 |
| | 4 | 12 | 6 | 9 | 75 |
| | 6 | 10 | 8 | 10 | 100 |

**g**

| Genotype | Diet Duration Chow only (months) | Number of mice analyzed | Age at which analyzed (months) | Number of mice with colitis | Percent of mice with colitis |
|---|---|---|---|---|---|
| WT | 0 | 10 | 2 | 0 | 0 |
| | 2 | 8 | 4 | 0 | 0 |
| | 3 | 8 | 5 | 0 | 0 |
| | 4 | 8 | 6 | 0 | 0 |
| | 6 | 8 | 8 | 0 | 0 |
| | 10 | 8 | 10 | 0 | 0 |
| TKO | 0 | 15 | 0 | 0 | 0 |
| | 2 | 10 | 4 | 0 | 0 |
| | 3 | 10 | 5 | 1 | 10 |
| | 4 | 10 | 6 | 3 | 30 |
| | 6 | 10 | 8 | 5 | 50 |
| | 10 | 10 | 10 | 8 | 80 |

**h**

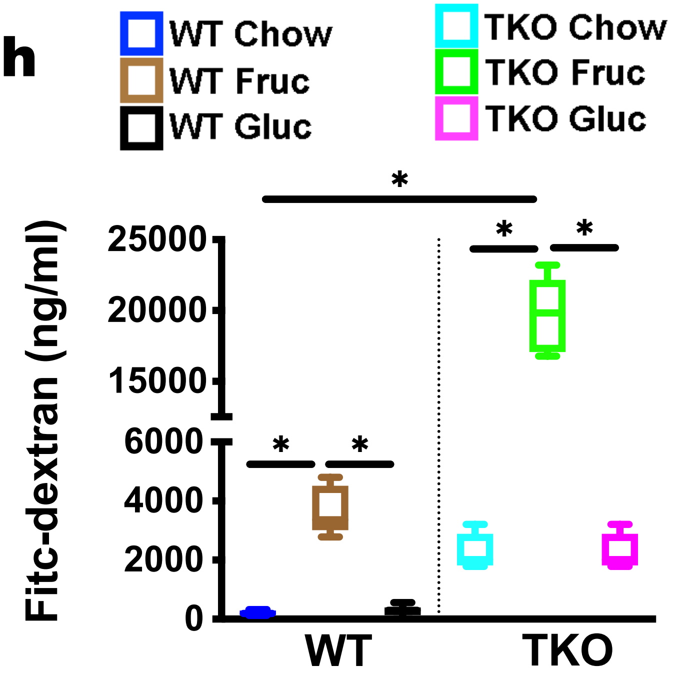


**i**

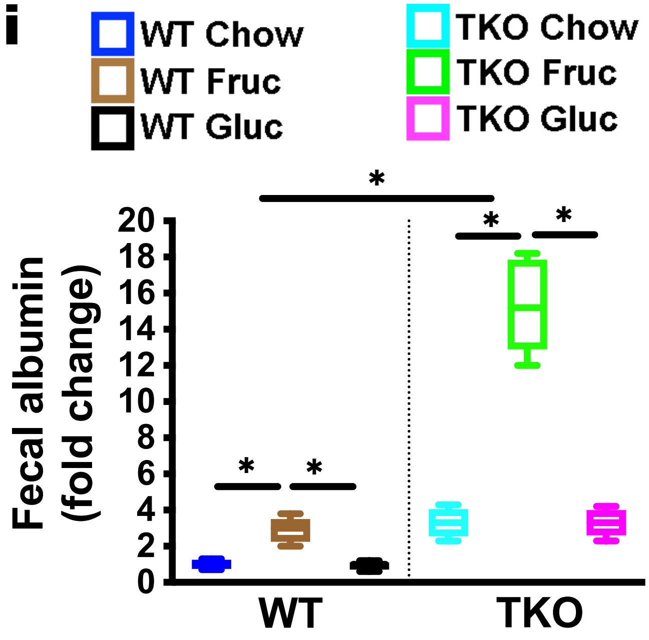


**j**

| Genotype | Diet Duration Chow +Glucose (months) | Number of mice analyzed | Age at which analyzed (months) | Mice with colitis | Percent of mice with colitis |
|---|---|---|---|---|---|
| WT | 0 | 20 | 2 | 0 | 0 |
| | 2 | 10 | 4 | 0 | 0 |
| | 3 | 10 | 5 | 0 | 0 |
| | 4 | 6 | 6 | 0 | 0 |
| | 6 | 6 | 8 | 0 | 0 |
| TKO | 0 | 20 | 2 | 0 | 0 |
| | 2 | 10 | 4 | 0 | 0 |
| | 3 | 10 | 5 | 0 | 0 |
| | 4 | 10 | 6 | 2 | 20 |
| | 6 | 10 | 8 | 5 | 50 |

sFig. 3

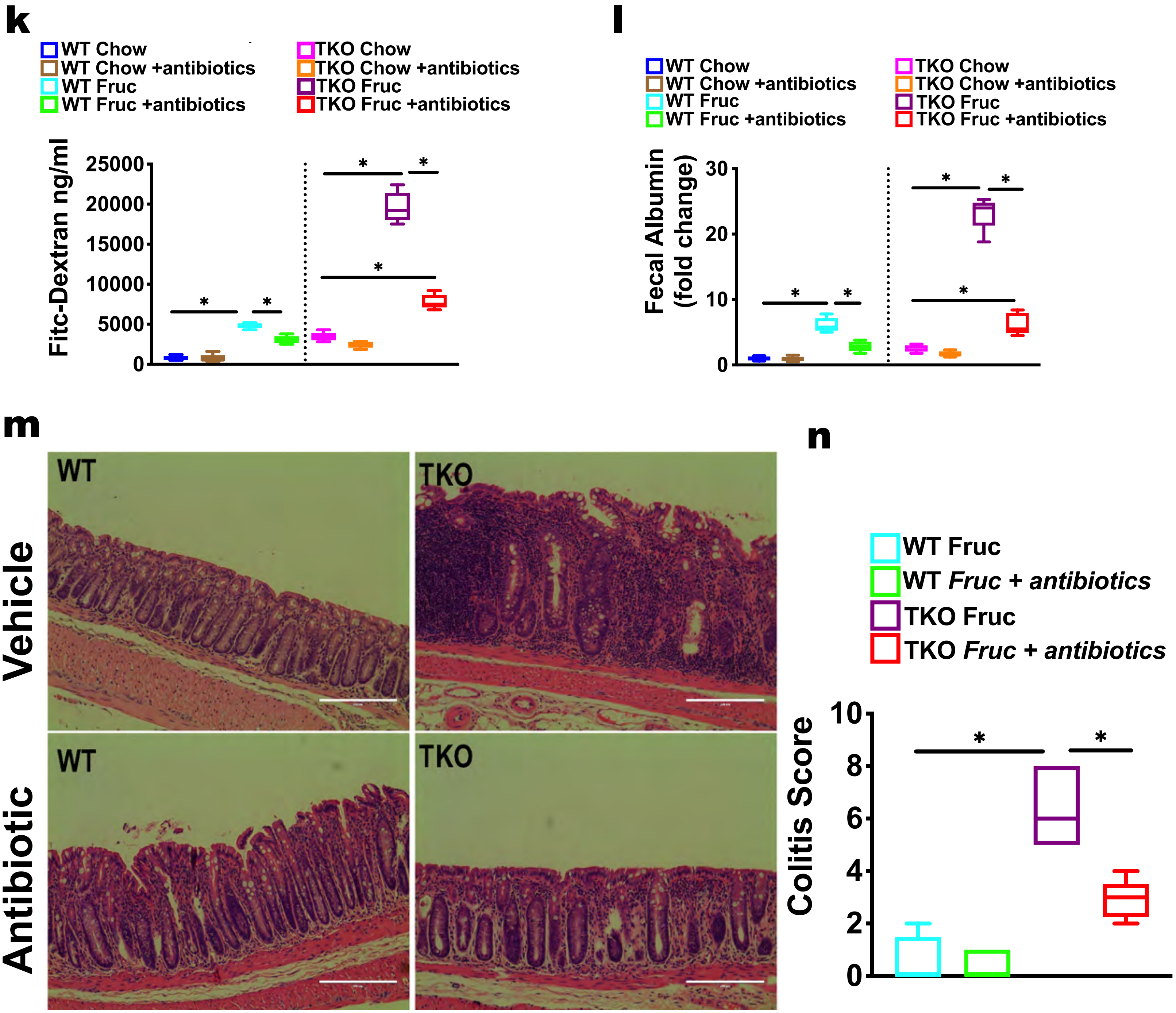
k
WT Chow
WT Chow +antibiotics
WT Fruc
WT Fruc +antibiotics
TKO Chow
TKO Chow +antibiotics
TKO Fruc
TKO Fruc +antibiotics
Fitc-Dextran ng/ml
25000
20000
15000
10000
5000
0
*
l
WT Chow
WT Chow +antibiotics
WT Fruc
WT Fruc +antibiotics
TKO Chow
TKO Chow +antibiotics
TKO Fruc
TKO Fruc +antibiotics
Fecal Albumin (fold change)
30
20
10
0
m
Vehicle
Antibiotic
WT
TKO
n
WT Fruc
WT Fruc + antibiotics
TKO Fruc
TKO Fruc + antibiotics
Colitis Score
10
8
6
4
2
0

sFig. 3

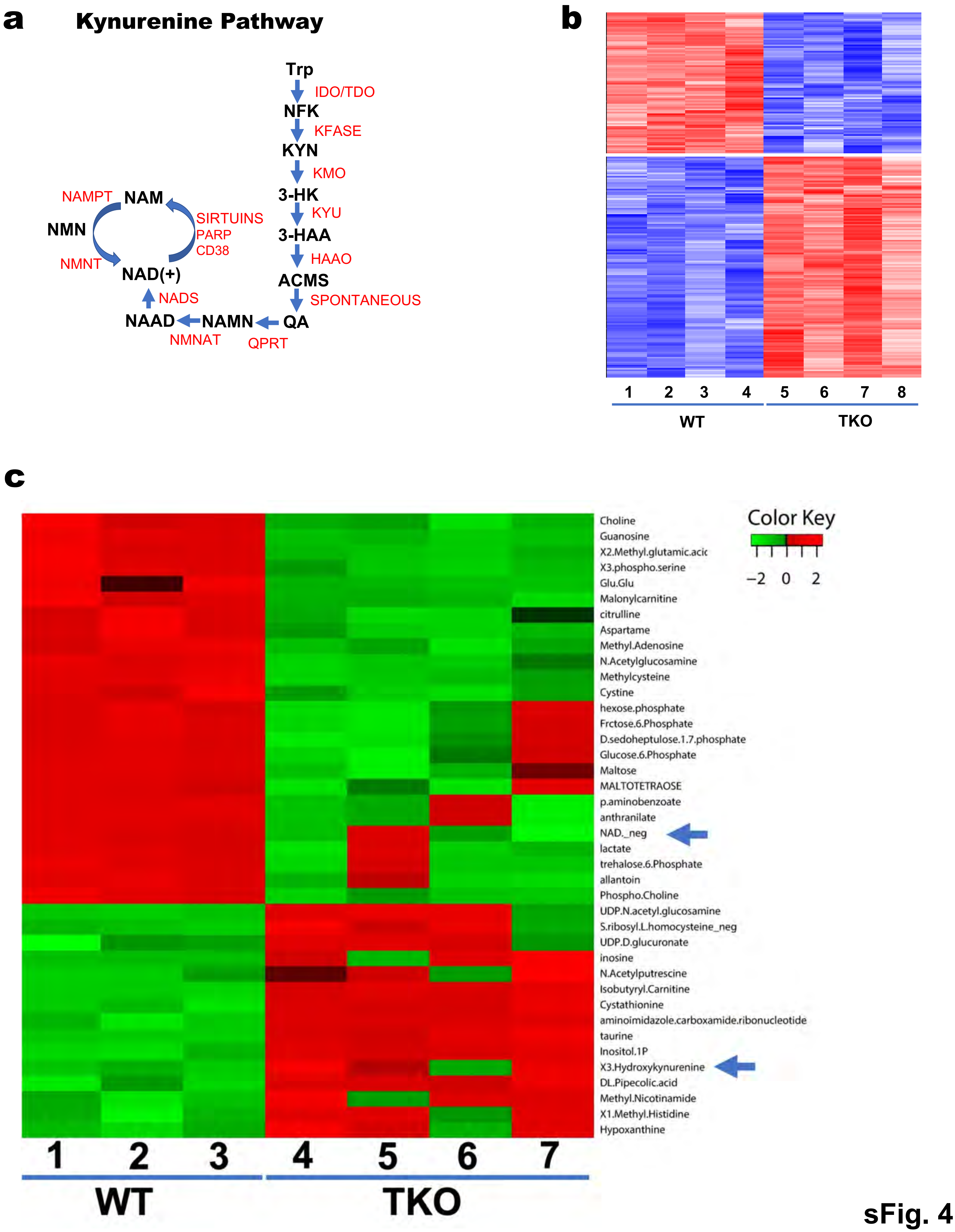
a
Kynurenine Pathway
Trp
IDO/TDO
NFK
KFASE
KYN
KMO
3-HK
KYU
3-HAA
HAAO
ACMS
SPONTANEOUS
QA
QPRT
NAMN
NMNAT
NAAD
NADS
NAD(+)
NAMPT
NAM
NMN
NMNT
SIRTUINS
PARP
CD38
b
1 2 3 4 5 6 7 8
WT
TKO
c
Choline
Guanosine
X2.Methyl.glutamic.acic
X3.phospho.serine
Glu.Glu
Malonylcarnitine
citrulline
Aspartame
Methyl.Adenosine
N.Acetylglucosamine
Methylcysteine
Cystine
hexose.phosphate
Frctose.6.Phosphate
D.sedoheptulose.1.7.phosphate
Glucose.6.Phosphate
Maltose
MALTOTETRAOSE
p.aminobenzoate
anthranilate
NAD._neg
lactate
trehalose.6.Phosphate
allantoin
Phospho.Choline
UDP.N.acetyl.glucosamine
S.ribosyl.L.homocysteine_neg
UDP.D.glucuronate
inosine
N.Acetylputrescine
Isobutyryl.Carnitine
Cystathionine
aminoimidazole.carboxamide.ribonucleotide
taurine
Inositol.1P
X3.Hydroxykynurenine
DL.Pipecolic.acid
Methyl.Nicotinamide
X1.Methyl.Histidine
Hypoxanthine
Color Key
−2 0 2
1 2 3 4 5 6 7
WT
TKO

sFig. 4

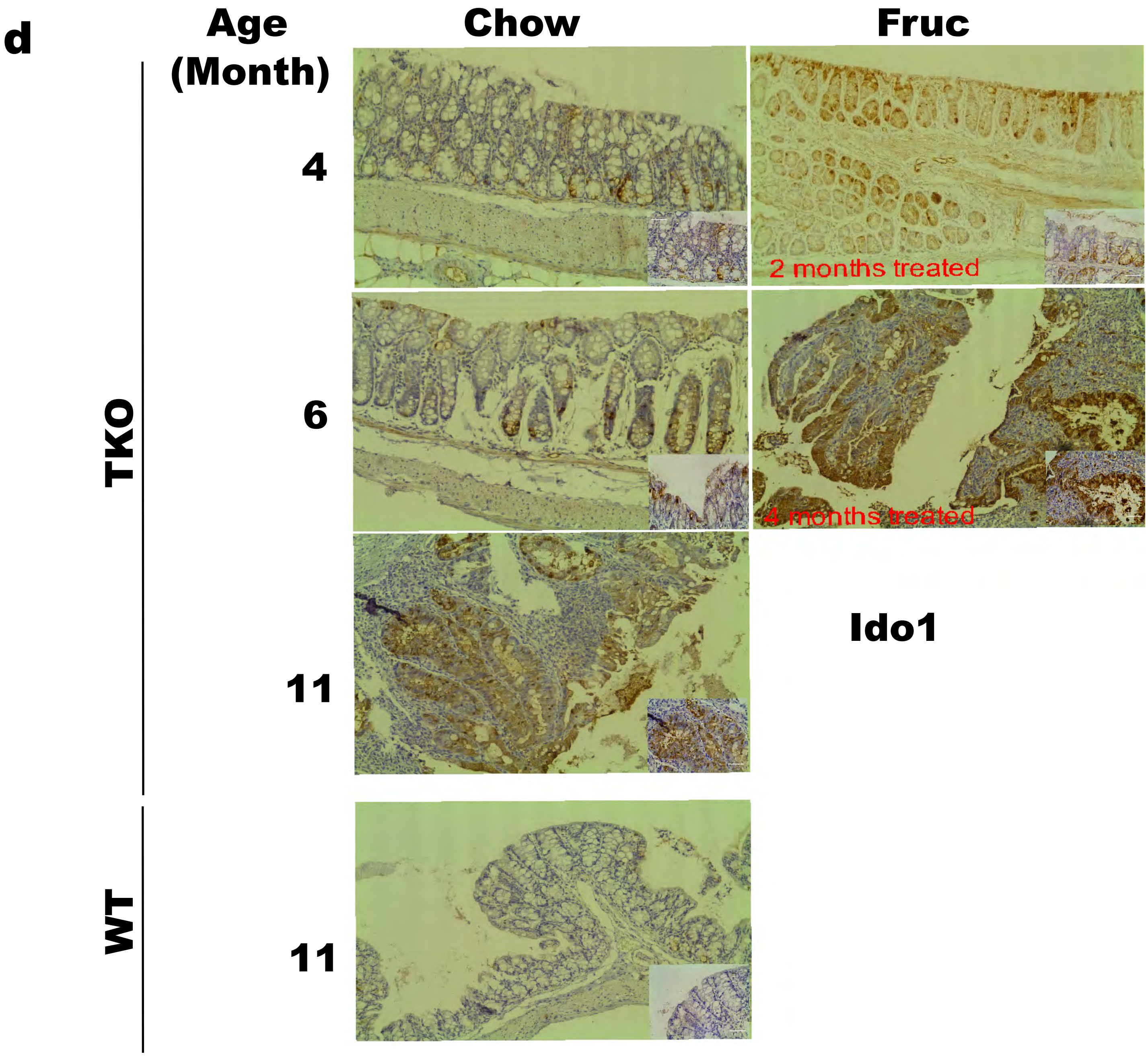
d
Age (Month)
Chow
Fruc
TKO
WT
4
6
11
11
2 months treated
4 months treated
Ido1

sFig. 4

**a**

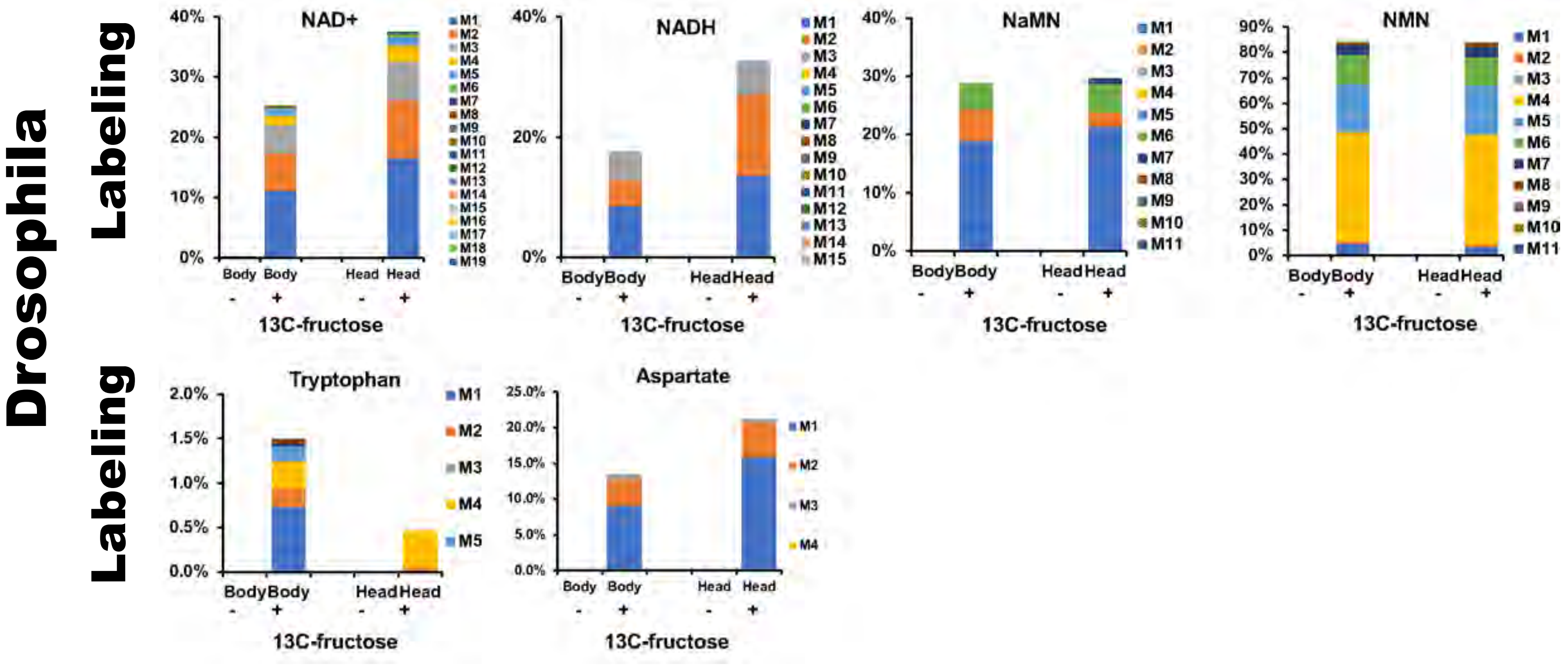
Drosophila
Labeling
NAD+
NADH
NaMN
NMN
Tryptophan
Aspartate
Body Body
Head Head
13C-fructose
M1
M2
M3
M4
M5
M6
M7
M8
M9
M10
M11
M12
M13
M14
M15
M16
M17
M18
M19

**b**

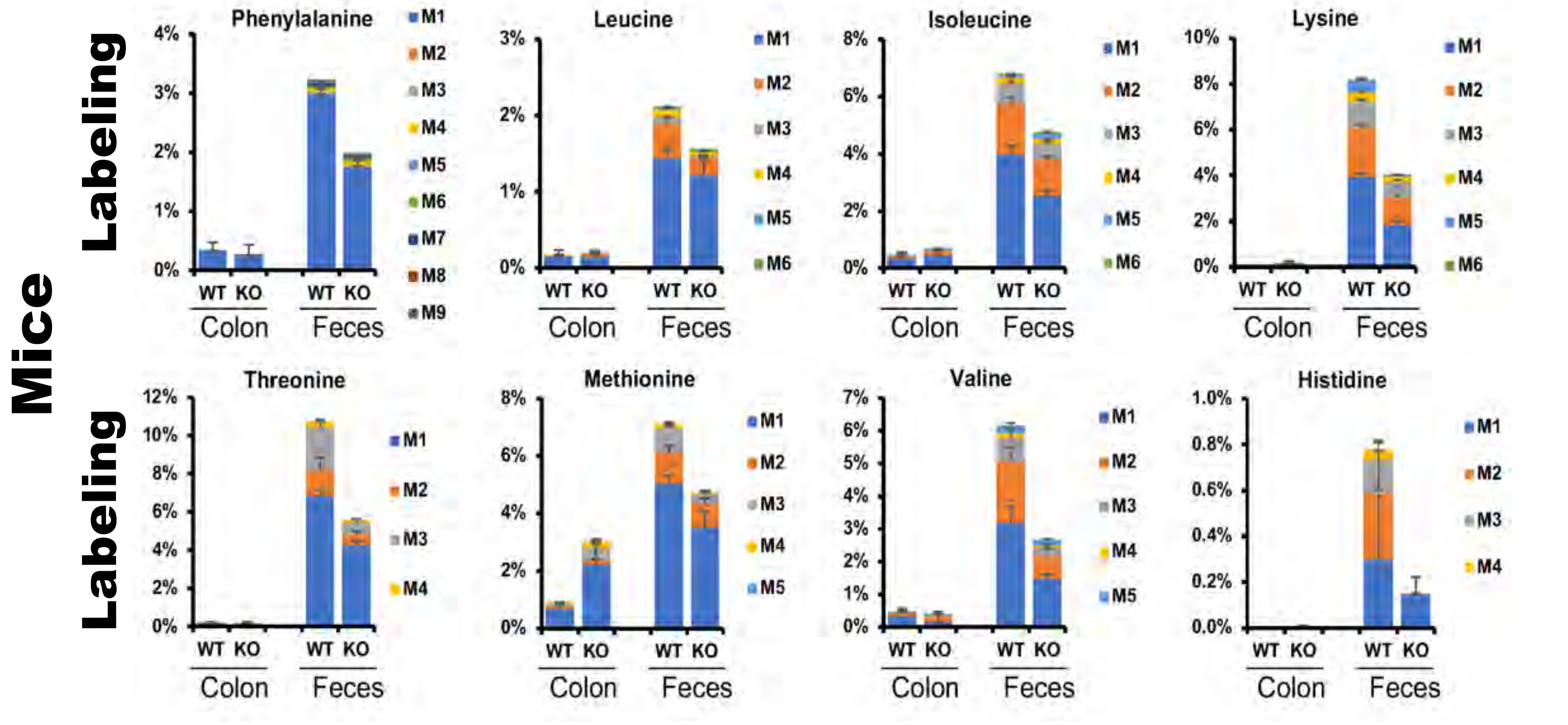
Mice
Labeling
Phenylalanine
Leucine
Isoleucine
Lysine
Threonine
Methionine
Valine
Histidine
WT KO
Colon
Feces
M1
M2
M3
M4
M5
M6
M7
M8
M9

**sFig. 5**

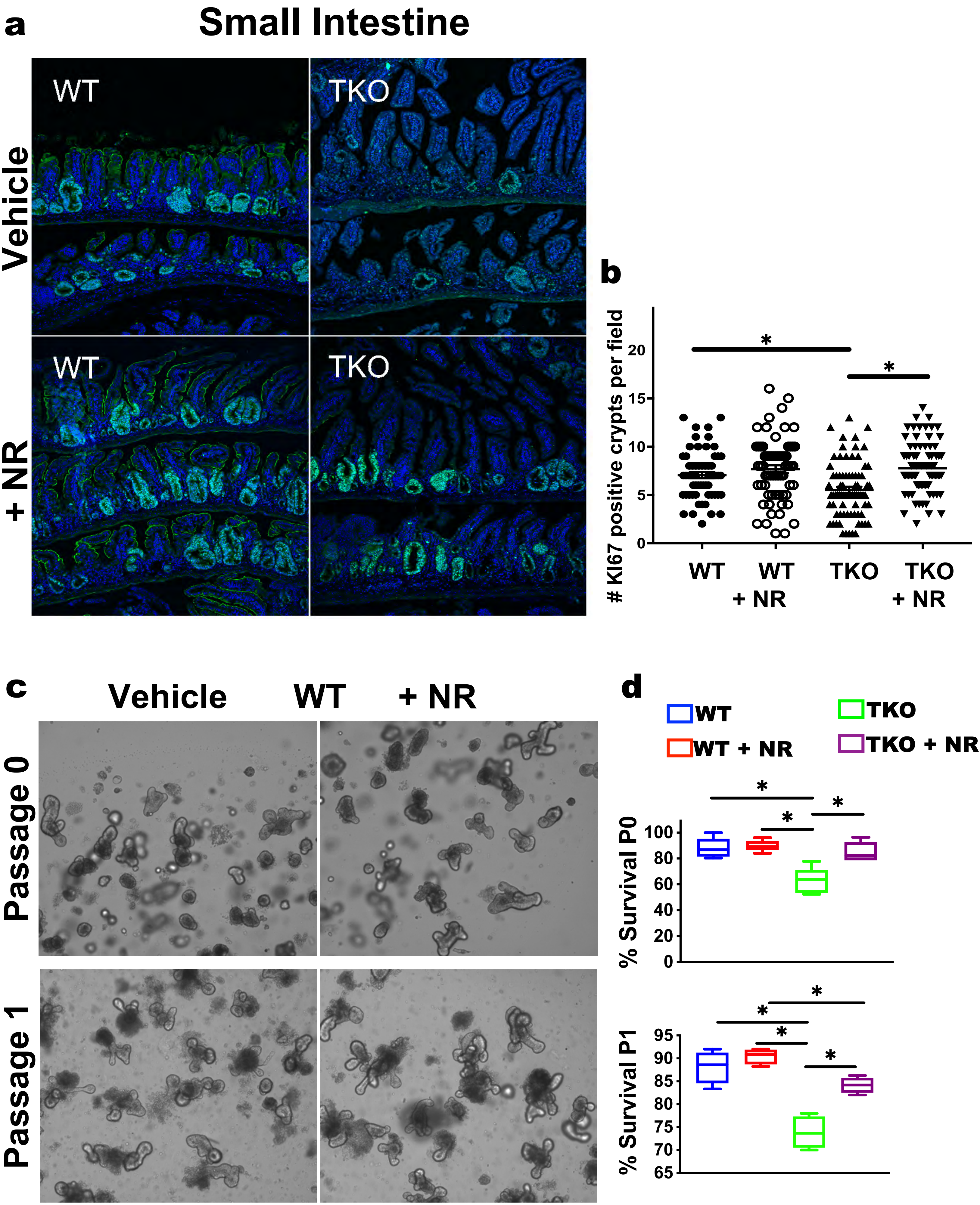
a
Small Intestine
WT
TKO
Vehicle
WT
TKO
+ NR
b
### KI67 positive crypts per field
20
15
10
5
0
*
*
WT
WT + NR
TKO
TKO + NR
c
Vehicle
WT
+ NR
Passage 0
Passage 1
d
WT
WT + NR
TKO
TKO + NR
% Survival P0
100
80
60
40
20
0
% Survival P1
95
90
85
80
75
70
65

sFig. 6

**sFig. 1 Telomere dysfunction causes decreased expression of transporters, microvilli structure and enzymes in the small intestine.** RNA-seq was carried out on intestinal epithelial scraps from four 14-week-old male wild type (WT) and telomerase deficient mice with short telomeres (TKO; n= 4 per group).

(a) All genes in transporter and microvilli-related pathway that are downregulated in TKO mice; threshold of differential expressed genes are FC > 2 and p < 0.05.

**sFig. 2 Telomere dysfunction impairs enterocyte maturation with loss of microvilli, malabsorption, and leaky barrier.** The small intestines of 14-16-week-old male WT and TKO mice were analyzed to assess morphology, absorption, and barrier function.

(a) Scanning electron microscopy (SEM) analysis of the villus tip of the proximal small intestine at low (6,000x), medium (22,000x) and high (66,000x) magnification shows patchy loss of microvilli in TKO mice (n= 3 per group; representative images).

(b) SEM of villus midsection at low, medium and high magnification shows patchy loss of microvilli in TKO mice (n= 3 per group; representative images).

(c) SEM analysis of WT and TKO colon shows absence of microvilli at low and high magnification in TKO mice (n= 3 per group; representative images)

(d) Transmission electron microscopy (TEM) images of villus tip (TIP), midsection (MID) and lower (LOW) part in the proximal intestine of TKO mice show abnormal microvilli and areas without microvilli (n= 3 per group; representative images).

**sFig. 3 Telomere dysfunction impairs fructose absorption and predisposes to fructose-induced colitis and dysplasia**.

(a, b) Increased fecal concentrations (a) and decreased blood concentrations (b) of indicated amino acids (aa) in TKO mice show their malabsorption worsened by fructose feeding; n= 5 mice per group.

(c, d) Increased fecal levels (c) and decreased blood levels (b) of the indicated dipeptides in TKO mice show their malabsorption worsened by fructose feeding; n= 3 mice per group.

(e) Representative β-catenin and Ki67 immunohistochemistry images show nuclear accumulation of β-catenin in the epithelium of TKO mice and widespread proliferation; n= 3 per group.

(f, g) Incidence of colitis at different ages and length of diet in WT and TKO mice on a regular chow with fructose drinking (f) or normal water (g). Note the significant acceleration of colitis in TKO mice on fructose compared to TKO mice on regular chow; n= 8 - 20 per time point and diet.

(h-j) Blood Fitc-Dextran levels (h), fecal albumin concentration (i), and colitis incidence (j) indicates impaired barrier defect in TKO mice with fructose but not with glucose drinking; n = 5 for barrier function measurement and n= 8-10 for colitis quantification per timepoint.

(k, l) Blood Fitc-Dextran levels (k) and fecal albumin concentration (l) indicates partial rescue of barrier defect by antibiotics treatment in fructose-fed TKO mice; n= 5 mice per group.

(m, n) Representative H&E images (m) and colitis score (n) indicate partial rescue of colitis by antibiotics treatment in fructose-fed TKO mice; n= 8 per group.

Statistical differences were calculated using Student's t-test with *p < 0.05 considered to be statistical different.

**sFig. 4 Fructose induces dysregulated Kynurenine/NAD metabolism in TKO colons.**

(a) Schematic of the kynurenine and NAD metabolism in mammals. Black indicates metabolites and red indicates enzymes.
(b) RNA-seq analysis shows very distinct transcriptional profiles between WT and TKO colons after fructose feeding; n = 4 per group.
(c) Metabolomics analysis shows decreased NAD and increased hydroxykynurenine in TKO colons after fructose feeding for 2 months; n = 3 and 4 for WT and TKO mice, respectively.
(d) Representative Ido1 immunohistochemistry images with insets shows upregulation in TKO mice after 2 or 4 months of fructose feeding. Old TKO mice on chow show increased Ido1 expression while WT do not; n= 3 per group.

**sFig. 5 Fructose is used to generate NAD precursors, NAD or amino acids by fly or mouse microbiota.**

(a) Flies were fed [U-$^{13}$C]-labeled fructose or unlabeled fructose for 3 days as a sole carbon source and labeled metabolites in heads and bodies were analyzed separately by LC-MS. Labeled isotopomer distribution of NAD, NADH and the indicated NAD precursors (nicotinic acid mononucleotide, NaMN; nicotinamide mononucleotide, NMN; aspartate and tryptophan) indicates microbial biosynthesis. Fifty flies per group were pooled for analysis.

(b) Labeled isotopomer distribution of the indicated essential amino acids in WT and TKO mouse feces after 2 days of [U-$^{13}$C]-fructose drinking indicates their microbial synthesis; n= 3 per group.

Results are expressed as average ± s.e.

**sFig. 6 Increasing NAD levels via NR supplementation improves intestinal stem cell regeneration in TKO mice and enteroids.**

(a, b) Representative KI67 immunofluorescence staining in small intestines (a) and quantification of regenerating crypts in 10 longitudinal images taken at 40x per mouse (b) shows increased small intestinal crypt numbers in TKO mice with NR supplementation; n = 8 per group.

(c, d) Representative images of WT-derived enteroids (c) and quantitation of WT and TKO (d) shows no effect of NR supplementation on survival of enteroids from WT mice at isolation (P0) and after first passage (P1); n= 3 mice per group.

Statistical differences were calculated using Student’s t-test with *$p < 0.05$ considered to be statistical different.

### Supplemental Table 1 GOs and downregulated genes in TKO

| System | MSigDB | Contrast | Overlap Size | Contrast Size | MSigDB. Size | Background Size | FE | Pvalue | P.adj | Overlap.Items |
|---|---|---|---|---|---|---|---|---|---|---|
| C5.BP | GO_ORGANIC_HYDROXY_COMPOUND_METABOLIC_PROCESS | WT-TKO_up | 45 | 574 | 261 | 8100 | 2.43301694 | 1.55E-08 | 0.00018908 | Abat;Acaa1a;Acox2;Acss1;Bco1;Bmp2;Ces1b;Cln8;Cyp4f18;Dgat2;Epas1;Fdft1;Fdps;Fdx1;Galk1;Gch1;Got1;Gpd1;Gpd2;Hagh;Hmgcs1;Idh1;Inpp5a;Insig1;Ldlr;Lss;Maoa;Maob;Msmo1;Mvk;Naaa;Nr1d1;Nsdhl;Osbpl3;Osbpl6;Per2;Plcd1;Pmel;Por;Prkaa2;Qdpr;Rxra;Srd5a2;Srebf2;Sult2b1 |
| C5.MF | GO_ACTIVE_ION_TRANSMEMBRANE_TRANSPORTER_ACTIVITY | WT-TKO_up | 24 | 574 | 107 | 8100 | 3.16519587 | 3.00E-07 | 0.00127404 | Abcc3;Atp1a1;Atp1b1;Atp2a3;Atp2b1;Clcn5;Slc13a1;Slc13a2;Slc15a1;Slc20a1;Slc23a1;Slc26a11;Slc2a10;Slc2a8;Slc37a4;Slc4a11;Slc4a7;Slc5a1;Slc5a4a;Slc5a6;Slc6a19;Slc6a4;Slc6a6;Slc6a8 |
| C5.BP | GO_LIPID_METABOLIC_PROCESS | WT-TKO_up | 81 | 574 | 660 | 8100 | 1.7318657 | 3.94E-07 | 0.00127404 | Abhd1;Abhd2;Abhd5;Acaa1a;Acox2;Acsm5;Acss1;Atp1a1;B4galt5;B4galt6;Bco1;Bmp2;Capn2;Ces1b;Ces2a;Ces2c;Cish;Cln8;Cwh43;Cyp2j6;Cyp2s1;Cyp2u1;Cyp4f18;Dgat2;Dgkd;Elovl1;Enpp7;Fdft1;Fdps;Fdx1;Gpd1;Hmgcs1;Hpgd;Hsd17b13;Hsd3b2;Hsd3b3;Idh1;Inpp5a;Insig1;Irs2;Klf4;Ldlr;Lpin3;Lss;Msmo1;Mvk;Naaa;Neu1;Nr1d1;Nr1d2;Nr1h3;Nsdhl;Osbpl3;Osbpl6;Per2;Phospho1;Pisd;Plbd2;Plcd1;Plekha8;Plscr1;Pm20d1;Por;Pparg;Prkaa2;Rb1;Rorc;Rxra;Sdc1;Sdc3;Srd5a2;Srebf2;St8sia6;Sult2b1;Tfcp2l1;Tnfrsf21;Trib3;Ugcg;Ugt2b5;Wdtc1;Zbtb20 |
| C5.BP | GO_RESPONSE_TO_ENDOGENOUS_STIMULUS | WT-TKO_up | 87 | 574 | 728 | 8100 | 1.68640158 | 4.48E-07 | 0.00127404 | Abcc2;Abhd2;Actb;Adcy2;Anxa2;Anxa3;Arnt2;Atp1a1;Atp2b1;Bace1;Bcar1;Bmp2;Capn2;Cited2;Cldn4;Cpn1;Cry2;Cx3cr1;Eef2k;Endog;Ezr;Fdx1;Fgfbp3;Fgfrl1;Flcn;Foxo3;Fzd1;Garem1;Gclc;Gcnt2;Got1;Gpd1;Hmgcs1;Hpgd;Hsd3b2;Hsd3b3;Idh1;Irs2;Itpr2;Jup;Klf15;Klf4;Klf9;Lats2;Ldlr;Lpin3;Maob;Myo1c;Nbl1;Nptn;Nr1d1;Nr1d2;Nr1h3;Nr3c2;P2ry2;Per1;Pmepa1;Por;Pparg;Prkaa2;Prnp;Qdpr;Rab31;Rapgef1;Rb1;Rorc;Runx1;Rxra;Sdc1;Sh3gl2;Slc2a8;Slc6a4;Slc9a3r1;Smad7;Spint1;Spred2;Spry2;Srebf2;Stxbp1;Trib3;Ubc;Ugcg;Vamp2;Wbp2;Wdtc1;Zfp703;Zyx |
| C2.CP | REACTOME_BIOLOGICAL_OXIDATIONS | WT-TKO_up | 24 | 574 | 111 | 8100 | 3.05113476 | 6.16E-07 | 0.00127404 | Acsm5;Acss1;Acy3;Arnt2;Ces1b;Ces2a;Ces2c;Cndp2;Cyp2j6;Cyp2s1;Cyp2u1;Cyp4b1;Cyp4f18;Dpep1;Fdx1;Gclc;Gstm2;Maoa;Maob;Oplah;Por;Rxra;Sult2b1;Ugt2b5 |
| C5.BP | GO_STEROID_METABOLIC_PROCESS | WT-TKO_up | 31 | 574 | 168 | 8100 | 2.60390742 | 6.26E-07 | 0.00127404 | Acaa1a;Acox2;Atp1a1;Bmp2;Ces1b;Cln8;Dgat2;Fdft1;Fdps;Fdx1;Hmgcs1;Hsd3b2;Hsd3b3;Insig1;Ldlr;Lss;Msmo1;Mvk;Nr1d1;Nsdhl;Osbpl3;Osbpl6;Por;Prkaa2;Rorc;Rxra;Srd5a2;Srebf2;Sult2b1;Tfcp2l1;Ugt2b5 |
| C2.CP | REACTOME_METABOLISM_OF_STEROIDS | WT-TKO_up | 18 | 574 | 70 | 8100 | 3.62867098 | 1.11E-06 | 0.00169923 | Abcc3;Acox2;Fdft1;Fdps;Fdx1;Hmgcs1;Hsd3b2;Hsd3b3;Insig1;Lss;Msmo1;Mvk;Nsdhl;Osbpl3;Osbpl6;Rxra;Srd5a2;Srebf2 |
| C5.MF | GO_SYMPORTER_ACTIVITY | WT-TKO_up | 18 | 574 | 70 | 8100 | 3.62867098 | 1.11E-06 | 0.00169923 | Slc13a1;Slc13a2;Slc15a1;Slc16a5;Slc20a1;Slc23a1;Slc25a22;Slc2a10;Slc2a8;Slc4a11;Slc4a7;Slc5a1;Slc5a4a;Slc5a6;Slc6a19;Slc6a4;Slc6a6;Slc6a8 |
| C5.BP | GO_LIPID_BIOSYNTHETIC_PROCESS | WT-TKO_up | 50 | 574 | 352 | 8100 | 2.00447418 | 1.33E-06 | 0.00175947 | Abhd5;Acox2;Acsm5;Acss1;Atp1a1;B4galt5;B4galt6;Bco1;Bmp2;Capn2;Ces1b;Cln8;Cwh43;Dgat2;Elovl1;Fdft1;Fdps;Fdx1;Gpd1;Hmgcs1;Hpgd;Hsd17b13;Hsd3b2;Hsd3b3;Idh1;Insig1;Ldlr;Lpin3;Lss;Msmo1;Mvk;Nr1d1;Nr1h3;Nsdhl;Osbpl3;Osbpl6;Phospho1;Pisd;Plekha8;Plscr1;Por;Prkaa2;Srd5a2;Srebf2;St8sia6;Tfcp2l1;Trib3;Ugcg;Wdtc1;Zbtb20 |
| C5.BP | GO_ORGANIC_ACID_METABOLIC_PROCESS | WT-TKO_up | 68 | 574 | 541 | 8100 | 1.77371882 | 1.61E-06 | 0.00175947 | Abat;Abcc2;Abhd5;Acaa1a;Acox2;Acsm5;Acss1;Amdhd2;Azin1;Azin2;B4galt5;B4galt6;Btd;Ces1b;Ces2a;Ces2c;Chp1;Cyp2j6;Cyp2s1;Cyp2u1;Cyp4f18;Dgat2;Dpep1;Elovl1;Fbp1;Galk1;Gch1;Gclc;Glud1;Gnpda1;Gnpda2;Got1;Gpd1;Hagh;Hkdc1;Hoga1;Hpgd;Hykk;Idh1;Il15;Insig1;Irs2;Lpin3;Me2;Me3;Msmo1;Naaa;Npl;Nr1d1;Nr1h3;Osbpl3;Osbpl6;Per2;Pgm2;Pm20d1;Por;Pparg;Prkaa2;Qdpr;Sardh;Slc23a1;Slc5a6;Slc6a8;Sult2b1;Trib3;Ugt2b5;Wdtc1;Zbtb20 |
| C5.BP | GO_SMALL_MOLECULE_METABOLIC_PROCESS | WT-TKO_up | 100 | 574 | 899 | 8100 | 1.56968835 | 1.69E-06 | 0.00175947 | Abat;Abcc2;Abhd5;Acaa1a;Acot9;Acox2;Acsm5;Acss1;Adcy2;Amdhd2;Azin1;Azin2;B4galt5;B4galt6;Bco1;Bmp2;Btd;Ces1b;Ces2a;Ces2c;Chp1;Cln8;Coq10b;Cyp2j6;Cyp2s1;Cyp2u1;Cyp4f18;Dgat2;Dpep1;Efl1;Elovl1;Entpd7;Fbp1;Fdft1;Fdps;Fdx1;Flcn;Galk1;Gch1;Gclc;Glud1;Gnpda1;Gnpda2;Got1;Gpd1;Gpd2;Guca2a;Hagh;Hkdc1;Hmgcs1;Hoga1;Hpgd;Hykk;Idh1;Il15;Inpp5a;Insig1;Irs2;Ldlr;Lpin3;Lss;Me2;Me3;Mpp1;Msmo1;Mvk;Naaa;Nmrk1;Npl;Nr1d1;Nr1h3;Nsdhl;Osbpl3;Osbpl6;Pank3;Pde10a;Per2;Pgm2;Plcd1;Pm20d1;Por;Pparg;Prkaa2;Qdpr;Rnaseh2b;Rorc;Rxra;Sardh;Slc23a1;Slc37a4;Slc5a6;Slc6a8;Smpdl3a;Srd5a2;Srebf2;Sult2b1;Trib3;Ugt2b5;Wdtc1;Zbtb20 |
| C5.BP | GO_ANION_TRANSPORT | WT-TKO_up | 44 | 574 | 296 | 8100 | 2.09765515 | 1.73E-06 | 0.00175947 | Abat;Abcb1a;Abcc2;Abcc3;Clca4a;Clcn5;Cldn4;Clic1;Cln8;Cry2;Cyp4f18;Irs2;Ldlr;Lrrc8a;Per2;Plekha8;Plscr1;Pparg;Prkaa2;Prnp;Rxra;Slc13a1;Slc13a2;Slc16a5;Slc20a1;Slc23a1;Slc25a22;Slc26a11;Slc2a2;Slc2a8;Slc37a4;Slc3a1;Slc4a11;Slc4a7;Slc5a6;Slc6a19;Slc6a6;Slc6a8;Slc7a8;Slc9a3r1;Stxbp1;Ttyh3;Vamp2;Xkr9 |
| C5.BP | GO_STEROID_BIOSYNTHETIC_PROCESS | WT-TKO_up | 23 | 574 | 110 | 8100 | 2.950586 | 1.96E-06 | 0.0018001 | Acox2;Atp1a1;Bmp2;Ces1b;Fdft1;Fdps;Fdx1;Hmgcs1;Hsd3b2;Hsd3b3;Insig1;Lss;Msmo1;Mvk;Nr1d1;Nsdhl;Osbpl3;Osbpl6;Por;Prkaa2;Srd5a2;Srebf2;Tfcp2l1 |
| C5.MF | GO_ACTIVE_TRANSMEMBRANE_TRANSPORTER_ACTIVITY | WT-TKO_up | 29 | 574 | 160 | 8100 | 2.55770906 | 2.11E-06 | 0.0018001 | Abca8b;Abcb1a;Abcc2;Abcc3;Atp1a1;Atp1b1;Atp2a3;Atp2b1;Clcn5;Slc13a1;Slc13a2;Slc15a1;Slc16a5;Slc20a1;Slc23a1;Slc25a22;Slc26a11;Slc2a10;Slc2a8;Slc37a4;Slc4a11;Slc4a7;Slc5a1;Slc5a4a;Slc5a6;Slc6a19;Slc6a4;Slc6a6;Slc6a8 |
| C5.MF | GO_SOLUTE_CATION_SYMPORTER_ACTIVITY | WT-TKO_up | 15 | 574 | 53 | 8100 | 3.99382026 | 2.38E-06 | 0.0018001 | Slc13a1;Slc13a2;Slc15a1;Slc20a1;Slc23a1;Slc2a10;Slc2a8;Slc4a7;Slc5a1;Slc5a4a;Slc5a6;Slc6a19;Slc6a4;Slc6a6;Slc6a8 |
| C5.BP | GO_MONOCARBOXYLIC_ACID_METABOLIC_PROCESS | WT-TKO_up | 47 | 574 | 330 | 8100 | 2.00981945 | 2.60E-06 | 0.0018001 | Abat;Abcc2;Abhd5;Acaa1a;Acox2;Acsm5;Acss1;Btd;Ces1b;Ces2a;Ces2c;Cyp2j6;Cyp2s1;Cyp2u1;Cyp4f18;Dgat2;Elovl1;Fbp1;Galk1;Gpd1;Hagh;Hkdc1;Hoga1;Hpgd;Idh1;Insig1;Irs2;Lpin3;Me2;Me3;Msmo1;Naaa;Nr1d1;Nr1h3;Osbpl3;Osbpl6;Per2;Pgm2;Por;Pparg;Prkaa2;Slc5a6;Slc6a8;Trib3;Ugt2b5;Wdtc1;Zbtb20 |
| C5.MF | GO_SOLUTE_SODIUM_SYMPORTER_ACTIVITY | WT-TKO_up | 12 | 574 | 35 | 8100 | 4.83822797 | 2.63E-06 | 0.0018001 | Slc13a1;Slc13a2;Slc20a1;Slc23a1;Slc4a7;Slc5a1;Slc5a4a;Slc5a6;Slc6a19;Slc6a4;Slc6a6;Slc6a8 |
| C5.BP | GO_TRANSMEMBRANE_TRANSPORT | WT-TKO_up | 72 | 574 | 592 | 8100 | 1.7162633 | 2.65E-06 | 0.0018001 | Abca8b;Abcb1a;Abcc2;Abcc3;Actb;Ank3;Anxa2;Atp1a1;Atp1b1;Atp2a3;Atp2b1;Azin1;Azin2;Bcr;Bin1;Chp1;Clca4a;Clcn5;Cldn4;Clic1;Cln8;Cnnm1;Gjb3;Hap1;Irs2;Itpr2;Klf15;Lrrc8a;Mcoln1;Nipal1;Otop2;Per2;Plcd1;Pm20d1;Prkaa2;Prnp;S100a6;Scnn1a;Sgk3;Slc13a1;Slc13a2;Slc15a1;Slc16a5;Slc20a1;Slc23a1;Slc25a22;Slc26a11;Slc2a10;Slc2a2;Slc2a5;Slc2a8;Slc30a2;Slc37a4;Slc3a1;Slc4a11;Slc4a7;Slc5a1;Slc5a4a;Slc5a6;Slc6a19;Slc6a4;Slc6a6;Slc6a8;Slc7a8;Slc9a3r1;Tmc7;Tmem144;Tmem63b;Trib3;Ttyh3;Ubc;Vamp2 |
| C5.BP | GO_ION_TRANSPORT | WT-TKO_up | 79 | 574 | 673 | 8100 | 1.65647602 | 3.32E-06 | 0.00206903 | Abat;Abcb1a;Abcc2;Abcc3;Ank3;Anxa2;Atp1a1;Atp1b1;Atp2a3;Atp2b1;Bin1;Chp1;Clca4a;Clcn5;Cldn4;Clic1;Cln8;Cnnm1;Cry2;Cyp4f18;Gja4;Hap1;Irs2;Itpr2;Ldlr;Lgals3;Lpar3;Lrrc8a;Maob;Mcoln1;Nipal1;Otop2;Per1;Per2;Plcd1;Plekha8;Plscr1;Pm20d1;Pparg;Prkaa2;Prnp;Pxk;Rxra;S100a6;Scnn1a;Sgk3;Slc13a1;Slc13a2;Slc15a1;Slc16a5;Slc20a1;Slc23a1;Slc25a22;Slc26a11;Slc2a10;Slc2a2;Slc2a8;Slc30a2;Slc37a4;Slc3a1;Slc4a11;Slc4a7;Slc5a1;Slc5a4a;Slc5a6;Slc6a19;Slc6a4;Slc6a6;Slc6a8;Slc7a8;Slc9a3r1;Stxbp1;Syt12;Syt7;Tmc7;Tmem63b;Ttyh3;Vamp2;Xkr9 |
| C5.BP | GO_ANION_TRANSMEMBRANE_TRANSPORT | WT-TKO_up | 26 | 574 | 138 | 8100 | 2.65868808 | 3.39E-06 | 0.00206903 | Abcb1a;Clca4a;Clcn5;Cldn4;Clic1;Cln8;Irs2;Lrrc8a;Per2;Prkaa2;Prnp;Slc13a1;Slc20a1;Slc23a1;Slc25a22;Slc26a11;Slc37a4;Slc3a1;Slc4a11;Slc4a7;Slc5a6;Slc6a19;Slc6a6;Slc6a8;Slc7a8;Ttyh3 |
| C5.BP | GO_REGULATION_OF_CIRCADIAN_RHYTHM | WT-TKO_up | 13 | 574 | 42 | 8100 | 4.3678447 | 3.73E-06 | 0.00217022 | Bhlhe40;Cipc;Cry2;Nr1d1;Nr1d2;Nr1h3;Per1;Per2;Per3;Pparg;Prkaa2;Rorc;Usp2 |

| | | | | | | | | | | |
|---|---|---|---|---|---|---|---|---|---|---|
| C5.MF | GO_SODIUM_ION_TRANSMEMBRANE_TRANSPORTER_ACTIVITY | WT-TKO_up | 16 | 574 | 62 | 8100 | 3.64167697 | 4.18E-06 | 0.00232142 | Atp1a1;Atp1b1;Scnn1a;Slc13a1;Slc13a2;Slc20a1;Slc23a1;Slc4a11;Slc4a7;Slc5a1;Slc5a4a;Slc5a6;Slc6a19;Slc6a4;Slc6a6;Slc6a8 |
| C5.BP | GO_CELLULAR_RESPONSE_TO_ENDOGENOUS_STIMULUS | WT-TKO_up | 72 | 574 | 603 | 8100 | 1.68495502 | 5.19E-06 | 0.00275848 | Abcc2;Abhd2;Actb;Adcy2;Atp1a1;Atp2b1;Bace1;Bcar1;Bmp2;Capn2;Cited2;Cry2;Cx3cr1;Eef2k;Ezr;Fdx1;Fgfbp3;Fgfrl1;Flcn;Foxo3;Fzd1;Garem1;Gclc;Gcnt2;Got1;Gpd1;Hmgcs1;Hpgd;Irs2;Itpr2;Jup;Klf15;Klf4;Klf9;Lats2;Lpin3;Myo1c;Nbl1;Nptn;Nr1d1;Nr1d2;Nr1h3;Nr3c2;P2ry2;Per1;Pmepa1;Por;Pparg;Prkaa2;Prnp;Rab31;Rapgef1;Rb1;Rorc;Runx1;Rxra;Sh3gl2;Slc2a8;Slc6a4;Slc9a3r1;Smad7;Spint1;Spred2;Spry2;Trib3;Ubc;Ugcg;Vamp2;Wbp2;Wdtc1;Zfp703;Zyx |
| C5.CC | GO_CLUSTER_OF_ACTIN_BASED_CELL_PROJECTIONS | WT-TKO_up | 20 | 574 | 93 | 8100 | 3.03473081 | 5.82E-06 | 0.00296081 | Abcc2;Clic1;Ezr;Mpp1;Myh10;Myo1a;Myo1c;Myo7b;Plec;Rhoc;Scin;Sis;Slc15a1;Slc2a2;Slc3a1;Slc4a7;Slc5a1;Slc5a6;Slc6a19;Slc9a3r1 |
| C2.CP | REACTOME_METABOLISM_OF_LIPIDS | WT-TKO_up | 47 | 574 | 343 | 8100 | 1.93364553 | 7.60E-06 | 0.00371572 | Abcc3;Abhd5;Acaa1a;Acot9;Acox2;Arnt2;Cidec;Cyp2j6;Cyp2u1;Cyp4b1;Cyp4f18;Dgat2;Dpep1;Elovl1;Enpp7;Fdft1;Fdps;Fdx1;Gpd1;Gpd2;Hmgcs1;Hpgd;Hsd17b13;Hsd3b2;Hsd3b3;Insig1;Lpin3;Lss;Msmo1;Mvk;Neu1;Nr1d1;Nr1h3;Nsdhl;Osbpl3;Osbpl6;Phospho1;Pisd;Plekha8;Pparg;Prkaa2;Rxra;Srd5a2;Srebf2;Tnfrsf21;Trib3;Ugcg |
| C5.CC | GO_BRUSH_BORDER | WT-TKO_up | 17 | 574 | 73 | 8100 | 3.28623932 | 9.49E-06 | 0.00444005 | Abcc2;Clic1;Ezr;Myh10;Myo1a;Myo1c;Myo7b;Plec;Scin;Sis;Slc15a1;Slc2a2;Slc3a1;Slc5a1;Slc5a6;Slc6a19;Slc9a3r1 |
| C5.BP | GO_ORGANIC_HYDROXY_COMPOUND_BIOSYNTHETIC_PROCESS | WT-TKO_up | 24 | 574 | 129 | 8100 | 2.62539503 | 1.01E-05 | 0.00444005 | Acox2;Bmp2;Ces1b;Fdft1;Fdps;Gch1;Got1;Hmgcs1;Insig1;Lss;Msmo1;Mvk;Nr1d1;Nsdhl;Osbpl3;Osbpl6;Per2;Plcd1;Pmel;Por;Prkaa2;Qdpr;Srd5a2;Srebf2 |
| C2.CP | REACTOME_PHASE_I_FUNCTIONALIZATION_OF_COMPOUNDS | WT-TKO_up | 15 | 574 | 59 | 8100 | 3.58766905 | 1.02E-05 | 0.00444005 | Acss1;Arnt2;Ces1b;Ces2a;Ces2c;Cyp2j6;Cyp2s1;Cyp2u1;Cyp4b1;Cyp4f18;Fdx1;Maoa;Maob;Por;Rxra |
| C5.MF | GO_ION_TRANSMEMBRANE_TRANSPORTER_ACTIVITY | WT-TKO_up | 45 | 574 | 327 | 8100 | 1.9419493 | 1.07E-05 | 0.00450106 | Abcc2;Abcc3;Anxa2;Atp1a1;Atp1b1;Atp2a3;Atp2b1;Clca4a;Clcn5;Cldn4;Clic1;Itpr2;Lrrc8a;Mcoln1;Nipal1;Otop2;S100a6;Scnn1a;Slc13a1;Slc13a2;Slc15a1;Slc16a5;Slc20a1;Slc23a1;Slc25a22;Slc26a11;Slc2a10;Slc2a2;Slc2a8;Slc30a2;Slc37a4;Slc3a1;Slc4a11;Slc4a7;Slc5a1;Slc5a4a;Slc5a6;Slc6a19;Slc6a4;Slc6a6;Slc6a8;Slc7a8;Tmc7;Tmem63b;Ttyh3 |
| C5.BP | GO_ALCOHOL_METABOLIC_PROCESS | WT-TKO_up | 31 | 574 | 195 | 8100 | 2.24336639 | 1.58E-05 | 0.00642184 | Acss1;Bco1;Bmp2;Ces1b;Cln8;Dgat2;Fdft1;Fdps;Fdx1;Galk1;Gch1;Got1;Gpd1;Gpd2;Hmgcs1;Idh1;Inpp5a;Insig1;Ldlr;Lss;Msmo1;Mvk;Naaa;Nsdhl;Plcd1;Por;Prkaa2;Qdpr;Rxra;Srebf2;Sult2b1 |
| C5.BP | GO_REGULATION_OF_LIPID_BIOSYNTHETIC_PROCESS | WT-TKO_up | 21 | 574 | 108 | 8100 | 2.74390244 | 1.79E-05 | 0.0070518 | Atp1a1;Bmp2;Capn2;Dgat2;Fdft1;Fdps;Hmgcs1;Hsd17b13;Idh1;Insig1;Ldlr;Lss;Mvk;Nr1d1;Nr1h3;Por;Prkaa2;Srebf2;Trib3;Wdtc1;Zbtb20 |
| C5.BP | GO_PHOTOPERIODISM | WT-TKO_up | 6 | 574 | 10 | 8100 | 8.46689896 | 2.03E-05 | 0.00736294 | Bhlhe40;Cry2;Per1;Per2;Per3;Usp2 |
| C5.BP | GO_ENTRAINMENT_OF_CIRCADIAN_CLOCK_BY_PHO | WT-TKO_up | 6 | 574 | 10 | 8100 | 8.46689896 | 2.03E-05 | 0.00736294 | Bhlhe40;Cry2;Per1;Per2;Per3;Usp2 |
| C5.BP | GO_CHEMICAL_HOMEOSTASIS | WT-TKO_up | 62 | 574 | 516 | 8100 | 1.69556762 | 2.05E-05 | 0.00736294 | Abcc2;Abhd5;Ackr4;Acox2;Adcy2;Ank3;Anxa2;Anxa7;Atp1a1;Atp1b1;Atp2a3;Atp2b1;Bnip3;Chp1;Cldn4;Cry2;Cx3cr1;Cxcr3;Cyp4f18;Dgat2;Elovl1;Endog;Epas1;F2rl1;Foxo3;Gclc;Got1;Hap1;Herpud1;Hkdc1;Hmox1;Insig1;Irs2;Itpr2;Klf15;Ldlr;Lpar3;Mcoln1;Mt1;Mt2;Nptn;Nr1d1;Nr1d2;Nr1h3;P2ry2;Plcd1;Pparg;Prkaa2;Prnp;S100a14;Scnn1a;Slc30a2;Slc4a11;Slc4a7;Slc7a8;Slc9a3r1;Smad7;Srebf2;Synpo;Ugcg;Wdtc1;Zbtb20 |
| C5.MF | GO_TRANSCRIPTION_COFACTOR_BINDING | WT-TKO_up | 8 | 574 | 19 | 8100 | 5.94168348 | 2.27E-05 | 0.00793315 | Klf4;Mafk;Nek6;Nr1d1;Per1;Per2;Per3;Rorc |
| C5.BP | GO_CELLULAR_RESPONSE_TO_ORGANIC_CYCLIC_COMPOUND | WT-TKO_up | 38 | 574 | 269 | 8100 | 1.99344585 | 2.92E-05 | 0.00990871 | Abcc2;Abhd2;Actb;Adcy2;Atp1a1;Atp2b1;Bmp2;Cry2;Eef2k;Ezr;Fdx1;Gclc;Gpd1;Hmgcs1;Il15;Insig1;Itpr2;Jup;Klf4;Klf9;Nr1d1;Nr1d2;Nr1h3;Nr3c2;P2ry2;Per1;Pmepa1;Pparg;Ralb;Rapgef1;Rb1;Rorc;Runx1;Rxra;Slc6a4;Slc9a3r1;Wbp2;Zfp703 |
| C5.MF | GO_SECONDARY_ACTIVE_TRANSMEMBRANE_TRANSPORTER_ACTIVITY | WT-TKO_up | 21 | 574 | 112 | 8100 | 2.64590592 | 3.18E-05 | 0.01034658 | Clcn5;Slc13a1;Slc13a2;Slc15a1;Slc16a5;Slc20a1;Slc23a1;Slc25a22;Slc26a11;Slc2a10;Slc2a8;Slc37a4;Slc4a11;Slc4a7;Slc5a1;Slc5a4a;Slc5a6;Slc6a19;Slc6a4;Slc6a6;Slc6a8 |
| C5.MF | GO_TRANSCRIPTION_COREPRESSOR_BINDING | WT-TKO_up | 5 | 574 | 7 | 8100 | 10.0796416 | 3.27E-05 | 0.01034658 | Nek6;Nr1d1;Per1;Per2;Per3 |
| C2.CP | REACTOME_CHOLESTEROL_BIOSYNTHESIS | WT-TKO_up | 7 | 574 | 15 | 8100 | 6.58536585 | 3.37E-05 | 0.01034658 | Fdft1;Fdps;Hmgcs1;Lss;Msmo1;Mvk;Nsdhl |
| C5.BP | GO_CIRCADIAN_RHYTHM | WT-TKO_up | 18 | 574 | 88 | 8100 | 2.88644283 | 3.48E-05 | 0.01034658 | Bhlhe40;Ciart;Cipc;Cldn4;Cry2;Dbp;Klf9;Nr1d1;Nr1d2;Nr1h3;Per1;Per2;Per3;Pparg;Prkaa2;Rorc;Slc6a4;Usp2 |
| C5.BP | GO_STEROL_METABOLIC_PROCESS | WT-TKO_up | 18 | 574 | 88 | 8100 | 2.88644283 | 3.48E-05 | 0.01034658 | Ces1b;Cln8;Dgat2;Fdft1;Fdps;Fdx1;Hmgcs1;Insig1;Ldlr;Lss;Msmo1;Mvk;Nsdhl;Por;Prkaa2;Rxra;Srebf2;Sult2b1 |
| C5.BP | GO_RESPONSE_TO_HORMONE | WT-TKO_up | 54 | 574 | 438 | 8100 | 1.73977376 | 3.56E-05 | 0.01034658 | Abcc2;Abhd2;Adcy2;Anxa2;Anxa3;Arnt2;Atp1a1;Atp2b1;Bcar1;Cldn4;Cpn1;Cry2;Eef2k;Endog;Foxo3;Gclc;Got1;Hmgcs1;Hpgd;Hsd3b2;Hsd3b3;Idh1;Irs2;Klf15;Klf9;Lats2;Lpin3;Maob;Myo1c;Nr1d1;Nr1d2;Nr1h3;Nr3c2;Per1;Pmepa1;Por;Pparg;Prkaa2;Qdpr;Rab31;Rb1;Rorc;Runx1;Rxra;Sdc1;Slc2a8;Slc6a4;Stxbp1;Trib3;Ugcg;Vamp2;Wbp2;Wdtc1;Zfp703 |
| C5.MF | GO_ANION_TRANSMEMBRANE_TRANSPORTER_ACTIVITY | WT-TKO_up | 25 | 574 | 148 | 8100 | 2.38369903 | 3.72E-05 | 0.01056519 | Abcc2;Abcc3;Clca4a;Clcn5;Cldn4;Clic1;Lrrc8a;Slc13a1;Slc13a2;Slc16a5;Slc23a1;Slc25a22;Slc26a11;Slc2a2;Slc2a8;Slc37a4;Slc3a1;Slc4a11;Slc4a7;Slc6a19;Slc6a4;Slc6a6;Slc6a8;Slc7a8;Ttyh3 |
| C5.BP | GO_ENTRAINMENT_OF_CIRCADIAN_CLOCK | WT-TKO_up | 6 | 574 | 11 | 8100 | 7.69718087 | 4.19E-05 | 0.01163418 | Bhlhe40;Cry2;Per1;Per2;Per3;Usp2 |
| C2.CP | REACTOME_TRANSPORT_OF_SMALL_MOLECULES | WT-TKO_up | 43 | 574 | 326 | 8100 | 1.86133259 | 4.76E-05 | 0.01271896 | Abca8b;Abcb1a;Abcc2;Abcc3;Adcy2;Atp1a1;Atp1b1;Atp2a3;Atp2b1;Bmp1;Cidec;Clca4a;Clcn5;Creb3l3;Hmox1;Ldlr;Lrrc8a;Mcoln1;Nipal1;Nr1h3;Runx1;Scnn1a;Sgk3;Slc13a1;Slc13a2;Slc15a1;Slc20a1;Slc25a22;Slc26a11;Slc2a10;Slc2a2;Slc2a8;Slc30a2;Slc3a1;Slc4a7;Slc5a1;Slc5a4a;Slc5a6;Slc6a19;Slc6a6;Slc7a8;Ttyh3;Ubc |
| C5.BP | GO_RHYTHMIC_PROCESS | WT-TKO_up | 21 | 574 | 115 | 8100 | 2.57688229 | 4.79E-05 | 0.01271896 | Bhlhe40;Ciart;Cipc;Cldn4;Cry2;Dbp;Enox1;Hlf;Klf9;Nr1d1;Nr1d2;Nr1h3;Per1;Per2;Per3;Pparg;Prkaa2;Rorc;Slc6a4;Tef;Usp2 |
| C5.BP | GO_REGULATION_OF_CELLULAR_COMPONENT_MOVEMENT | WT-TKO_up | 60 | 574 | 509 | 8100 | 1.66343791 | 4.97E-05 | 0.01290973 | Abhd2;Acvr1b;Amotl2;Anxa3;Atp1a1;Bcar1;Bcas3;Bcr;Bin1;Bmp2;Cadm4;Cited2;Cldn4;Cx3cr1;Cxcl16;Cxcr3;Dock1;Dpep1;Dusp10;Epha1;F2rl1;F3;Fam83h;Flcn;Foxo3;Gcnt2;Hap1;Hmox1;Irs2;Jup;Klf4;Lamc2;Lgals3;Map2k3;Mapre2;Mpp1;Myo1c;Nbl1;Nedd9;Pard6b;Plxna2;Pparg;Prag1;Prkx;Ptprr;Rhob;Rhoc;Rhof;S100a14;Sema3b;Sema7a;Sgk3;Slc9a3r1;Smad7;Spata13;Spry2;Stk10;Syde1;Zfp703;Zswim4 |
| C5.BP | GO_ORGANIC_ANION_TRANSPORT | WT-TKO_up | 35 | 574 | 246 | 8100 | 2.00773349 | 5.19E-05 | 0.01320803 | Abat;Abcb1a;Abcc2;Abcc3;Cln8;Cyp4f18;Irs2;Ldlr;Lrrc8a;Per2;Plekha8;Plscr1;Pparg;Prkaa2;Rxra;Slc13a1;Slc13a2;Slc16a5;Slc23a1;Slc25a22;Slc2a2;Slc2a8;Slc37a4;Slc3a1;Slc4a11;Slc4a7; |

| | | | | | | | | | | |
|---|---|---|---|---|---|---|---|---|---|---|
| C5.MF | GO_SODIUM_ION_TRANSMEMBRANE_TRANSPORTER_ACTIVITY | WT-TKO_up | 16 | 574 | 62 | 8100 | 3.64167697 | 4.18E-06 | 0.00232142 | Atp1a1;Atp1b1;Scnn1a;Slc13a1;Slc13a2;Slc20a1;Slc23a1;Slc4a11;Slc4a7;Slc5a1;Slc5a4a;Slc5a6;Slc6a19;Slc6a4;Slc6a6;Slc6a8 |
| C5.BP | GO_CELLULAR_RESPONSE_TO_ENDOGENOUS_STIMULUS | WT-TKO_up | 72 | 574 | 603 | 8100 | 1.68495502 | 5.19E-06 | 0.00275848 | Abcc2;Abhd2;Actb;Adcy2;Atp1a1;Atp2b1;Bace1;Bcar1;Bmp2;Capn2;Cited2;Cry2;Cx3cr1;Eef2k;Ezr;Fdx1;Fgfbp3;Fgfrl1;Flcn;Foxo3;Fzd1;Garem1;Gclc;Gcnt2;Got1;Gpd1;Hmgcs1;Hpgd;Irs2;Itpr2;Jup;Klf15;Klf4;Klf9;Lats2;Lpin3;Myo1c;Nbl1;Nptn;Nr1d1;Nr1d2;Nr1h3;Nr3c2;P2ry2;Per1;Pmepa1;Por;Pparg;Prkaa2;Prnp;Rab31;Rapgef1;Rb1;Rorc;Runx1;Rxra;Sh3gl2;Slc2a8;Slc6a4;Slc9a3r1;Smad7;Spint1;Spred2;Spry2;Trib3;Ubc;Ugcg;Vamp2;Wbp2;Wdtc1;Zfp703;Zyx |
| C5.CC | GO_CLUSTER_OF_ACTIN_BASED_CELL_PROJECTIONS | WT-TKO_up | 20 | 574 | 93 | 8100 | 3.03473081 | 5.82E-06 | 0.00296081 | Abcc2;Clic1;Ezr;Mpp1;Myh10;Myo1a;Myo1c;Myo7b;Plec;Rhoc;Scin;Sis;Slc15a1;Slc2a2;Slc3a1;Slc4a7;Slc5a1;Slc5a6;Slc6a19;Slc9a3r1 |
| C2.CP | REACTOME_METABOLISM_OF_LIPIDS | WT-TKO_up | 47 | 574 | 343 | 8100 | 1.93364553 | 7.60E-06 | 0.00371572 | Abcc3;Abhd5;Acaa1a;Acot9;Acox2;Arnt2;Cidec;Cyp2j6;Cyp2u1;Cyp4b1;Cyp4f18;Dgat2;Dpep1;Elovl1;Enpp7;Fdft1;Fdps;Fdx1;Gpd1;Gpd2;Hmgcs1;Hpgd;Hsd17b13;Hsd3b2;Hsd3b3;Insig1;Lpin3;Lss;Msmo1;Mvk;Neu1;Nr1d1;Nr1h3;Nsdhl;Osbpl3;Osbpl6;Phospho1;Pisd;Plekha8;Pparg;Prkaa2;Rxra;Srd5a2;Srebf2;Tnfrsf21;Trib3;Ugcg |
| C5.CC | GO_BRUSH_BORDER | WT-TKO_up | 17 | 574 | 73 | 8100 | 3.28623932 | 9.49E-06 | 0.00444005 | Abcc2;Clic1;Ezr;Myh10;Myo1a;Myo1c;Myo7b;Plec;Scin;Sis;Slc15a1;Slc2a2;Slc3a1;Slc5a1;Slc5a6;Slc6a19;Slc9a3r1 |
| C5.BP | GO_ORGANIC_HYDROXY_COMPOUND_BIOSYNTHETIC_PROCESS | WT-TKO_up | 24 | 574 | 129 | 8100 | 2.62539503 | 1.01E-05 | 0.00444005 | Acox2;Bmp2;Ces1b;Fdft1;Fdps;Gch1;Got1;Hmgcs1;Insig1;Lss;Msmo1;Mvk;Nr1d1;Nsdhl;Osbpl3;Osbpl6;Per2;Plcd1;Pmel;Por;Prkaa2;Qdpr;Srd5a2;Srebf2 |
| C2.CP | REACTOME_PHASE_I_FUNCTIONALIZATION_OF_COMPOUNDS | WT-TKO_up | 15 | 574 | 59 | 8100 | 3.58766905 | 1.02E-05 | 0.00444005 | Acss1;Arnt2;Ces1b;Ces2a;Ces2c;Cyp2j6;Cyp2s1;Cyp2u1;Cyp4b1;Cyp4f18;Fdx1;Maoa;Maob;Por;Rxra |
| C5.MF | GO_ION_TRANSMEMBRANE_TRANSPORTER_ACTIVITY | WT-TKO_up | 45 | 574 | 327 | 8100 | 1.9419493 | 1.07E-05 | 0.00450106 | Abcc2;Abcc3;Anxa2;Atp1a1;Atp1b1;Atp2a3;Atp2b1;Clca4a;Clcn5;Cldn4;Clic1;Itpr2;Lrrc8a;Mcoln1;Nipal1;Otop2;S100a6;Scnn1a;Slc13a1;Slc13a2;Slc15a1;Slc16a5;Slc20a1;Slc23a1;Slc25a22;Slc26a11;Slc2a10;Slc2a2;Slc2a8;Slc30a2;Slc37a4;Slc3a1;Slc4a11;Slc4a7;Slc5a1;Slc5a4a;Slc5a6;Slc6a19;Slc6a4;Slc6a6;Slc6a8;Slc7a8;Tmc7;Tmem63b;Ttyh3 |
| C5.BP | GO_ALCOHOL_METABOLIC_PROCESS | WT-TKO_up | 31 | 574 | 195 | 8100 | 2.24336639 | 1.58E-05 | 0.00642184 | Acss1;Bco1;Bmp2;Ces1b;Cln8;Dgat2;Fdft1;Fdps;Fdx1;Galk1;Gch1;Got1;Gpd1;Gpd2;Hmgcs1;Idh1;Inpp5a;Insig1;Ldlr;Lss;Msmo1;Mvk;Naaa;Nsdhl;Plcd1;Por;Prkaa2;Qdpr;Rxra;Srebf2;Sult2b1 |
| C5.BP | GO_REGULATION_OF_LIPID_BIOSYNTHETIC_PROCESS | WT-TKO_up | 21 | 574 | 108 | 8100 | 2.74390244 | 1.79E-05 | 0.0070518 | Atp1a1;Bmp2;Capn2;Dgat2;Fdft1;Fdps;Hmgcs1;Hsd17b13;Idh1;Insig1;Ldlr;Lss;Mvk;Nr1d1;Nr1h3;Por;Prkaa2;Srebf2;Trib3;Wdtc1;Zbtb20 |
| C5.BP | GO_PHOTOPERIODISM | WT-TKO_up | 6 | 574 | 10 | 8100 | 8.46689896 | 2.03E-05 | 0.00736294 | Bhlhe40;Cry2;Per1;Per2;Per3;Usp2 |
| C5.BP | GO_ENTRAINMENT_OF_CIRCADIAN_CLOCK_BY_PHO | WT-TKO_up | 6 | 574 | 10 | 8100 | 8.46689896 | 2.03E-05 | 0.00736294 | Bhlhe40;Cry2;Per1;Per2;Per3;Usp2 |
| C5.BP | GO_CHEMICAL_HOMEOSTASIS | WT-TKO_up | 62 | 574 | 516 | 8100 | 1.69556762 | 2.05E-05 | 0.00736294 | Abcc2;Abhd5;Ackr4;Acox2;Adcy2;Ank3;Anxa2;Anxa7;Atp1a1;Atp1b1;Atp2a3;Atp2b1;Bnip3;Chp1;Cldn4;Cry2;Cx3cr1;Cxcr3;Cyp4f18;Dgat2;Elovl1;Endog;Epas1;F2rl1;Foxo3;Gclc;Got1;Hap1;Herpud1;Hkdc1;Hmox1;Insig1;Irs2;Itpr2;Klf15;Ldlr;Lpar3;Mcoln1;Mt1;Mt2;Nptn;Nr1d1;Nr1d2;Nr1h3;P2ry2;Plcd1;Pparg;Prkaa2;Prnp;S100a14;Scnn1a;Slc30a2;Slc4a11;Slc4a7;Slc7a8;Slc9a3r1;Smad7;Srebf2;Synpo;Ugcg;Wdtc1;Zbtb20 |
| C5.MF | GO_TRANSCRIPTION_COFACTOR_BINDING | WT-TKO_up | 8 | 574 | 19 | 8100 | 5.94168348 | 2.27E-05 | 0.00793315 | Klf4;Mafk;Nek6;Nr1d1;Per1;Per2;Per3;Rorc |
| C5.BP | GO_CELLULAR_RESPONSE_TO_ORGANIC_CYCLIC_COMPOUND | WT-TKO_up | 38 | 574 | 269 | 8100 | 1.99344585 | 2.92E-05 | 0.00990871 | Abcc2;Abhd2;Actb;Adcy2;Atp1a1;Atp2b1;Bmp2;Cry2;Eef2k;Ezr;Fdx1;Gclc;Gpd1;Hmgcs1;Il15;Insig1;Itpr2;Jup;Klf4;Klf9;Nr1d1;Nr1d2;Nr1h3;Nr3c2;P2ry2;Per1;Pmepa1;Pparg;Ralb;Rapgef1;Rb1;Rorc;Runx1;Rxra;Slc6a4;Slc9a3r1;Wbp2;Zfp703 |
| C5.MF | GO_SECONDARY_ACTIVE_TRANSMEMBRANE_TRANSPORTER_ACTIVITY | WT-TKO_up | 21 | 574 | 112 | 8100 | 2.64590592 | 3.18E-05 | 0.01034658 | Clcn5;Slc13a1;Slc13a2;Slc15a1;Slc16a5;Slc20a1;Slc23a1;Slc25a22;Slc26a11;Slc2a10;Slc2a8;Slc37a4;Slc4a11;Slc4a7;Slc5a1;Slc5a4a;Slc5a6;Slc6a19;Slc6a4;Slc6a6;Slc6a8 |
| C5.MF | GO_TRANSCRIPTION_COREPRESSOR_BINDING | WT-TKO_up | 5 | 574 | 7 | 8100 | 10.0796416 | 3.27E-05 | 0.01034658 | Nek6;Nr1d1;Per1;Per2;Per3 |
| C2.CP | REACTOME_CHOLESTEROL_BIOSYNTHESIS | WT-TKO_up | 7 | 574 | 15 | 8100 | 6.58536585 | 3.37E-05 | 0.01034658 | Fdft1;Fdps;Hmgcs1;Lss;Msmo1;Mvk;Nsdhl |
| C5.BP | GO_CIRCADIAN_RHYTHM | WT-TKO_up | 18 | 574 | 88 | 8100 | 2.88644283 | 3.48E-05 | 0.01034658 | Bhlhe40;Ciart;Cipc;Cldn4;Cry2;Dbp;Klf9;Nr1d1;Nr1d2;Nr1h3;Per1;Per2;Per3;Pparg;Prkaa2;Rorc;Slc6a4;Usp2 |
| C5.BP | GO_STEROL_METABOLIC_PROCESS | WT-TKO_up | 18 | 574 | 88 | 8100 | 2.88644283 | 3.48E-05 | 0.01034658 | Ces1b;Cln8;Dgat2;Fdft1;Fdps;Fdx1;Hmgcs1;Insig1;Ldlr;Lss;Msmo1;Mvk;Nsdhl;Por;Prkaa2;Rxra;Srebf2;Sult2b1 |
| C5.BP | GO_RESPONSE_TO_HORMONE | WT-TKO_up | 54 | 574 | 438 | 8100 | 1.73977376 | 3.56E-05 | 0.01034658 | Abcc2;Abhd2;Adcy2;Anxa2;Anxa3;Arnt2;Atp1a1;Atp2b1;Bcar1;Cldn4;Cpn1;Cry2;Eef2k;Endog;Foxo3;Gclc;Got1;Hmgcs1;Hpgd;Hsd3b2;Hsd3b3;Idh1;Irs2;Klf15;Klf9;Lats2;Lpin3;Maob;Myo1c;Nr1d1;Nr1d2;Nr1h3;Nr3c2;Per1;Pmepa1;Por;Pparg;Prkaa2;Qdpr;Rab31;Rb1;Rorc;Runx1;Rxra;Sdc1;Slc2a8;Slc6a4;Stxbp1;Trib3;Ugcg;Vamp2;Wbp2;Wdtc1;Zfp703 |
| C5.MF | GO_ANION_TRANSMEMBRANE_TRANSPORTER_ACTIVITY | WT-TKO_up | 25 | 574 | 148 | 8100 | 2.38369903 | 3.72E-05 | 0.01056519 | Abcc2;Abcc3;Clca4a;Clcn5;Cldn4;Clic1;Lrrc8a;Slc13a1;Slc13a2;Slc16a5;Slc23a1;Slc25a22;Slc26a11;Slc2a2;Slc2a8;Slc37a4;Slc3a1;Slc4a11;Slc4a7;Slc6a19;Slc6a4;Slc6a6;Slc6a8;Slc7a8;Ttyh3 |
| C5.BP | GO_ENTRAINMENT_OF_CIRCADIAN_CLOCK | WT-TKO_up | 6 | 574 | 11 | 8100 | 7.69718087 | 4.19E-05 | 0.01163418 | Bhlhe40;Cry2;Per1;Per2;Per3;Usp2 |
| C2.CP | REACTOME_TRANSPORT_OF_SMALL_MOLECULES | WT-TKO_up | 43 | 574 | 326 | 8100 | 1.86133259 | 4.76E-05 | 0.01271896 | Abca8b;Abcb1a;Abcc2;Abcc3;Adcy2;Atp1a1;Atp1b1;Atp2a3;Atp2b1;Bmp1;Cidec;Clca4a;Clcn5;Creb3l3;Hmox1;Ldlr;Lrrc8a;Mcoln1;Nipal1;Nr1h3;Runx1;Scnn1a;Sgk3;Slc13a1;Slc13a2;Slc15a1;Slc20a1;Slc25a22;Slc26a11;Slc2a10;Slc2a2;Slc2a8;Slc30a2;Slc3a1;Slc4a7;Slc5a1;Slc5a4a;Slc5a6;Slc6a19;Slc6a6;Slc7a8;Ttyh3;Ubc |
| C5.BP | GO_RHYTHMIC_PROCESS | WT-TKO_up | 21 | 574 | 115 | 8100 | 2.57688229 | 4.79E-05 | 0.01271896 | Bhlhe40;Ciart;Cipc;Cldn4;Cry2;Dbp;Enox1;Hlf;Klf9;Nr1d1;Nr1d2;Nr1h3;Per1;Per2;Per3;Pparg;Prkaa2;Rorc;Slc6a4;Tef;Usp2 |
| C5.BP | GO_REGULATION_OF_CELLULAR_COMPONENT_MOVEMENT | WT-TKO_up | 60 | 574 | 509 | 8100 | 1.66343791 | 4.97E-05 | 0.01290973 | Abhd2;Acvr1b;Amotl2;Anxa3;Atp1a1;Bcar1;Bcas3;Bcr;Bin1;Bmp2;Cadm4;Cited2;Cldn4;Cx3cr1;Cxcl16;Cxcr3;Dock1;Dpep1;Dusp10;Epha1;F2rl1;F3;Fam83h;Flcn;Foxo3;Gcnt2;Hap1;Hmox1;Irs2;Jup;Klf4;Lamc2;Lgals3;Map2k3;Mapre2;Mpp1;Myo1c;Nbl1;Nedd9;Pard6b;Plxna2;Pparg;Prag1;Prkx;Ptprr;Rhob;Rhoc;Rhof;S100a14;Sema3b;Sema7a;Sgk3;Slc9a3r1;Smad7;Spata13;Spry2;Stk10;Syde1;Zfp703;Zswim4 |
| C5.BP | GO_ORGANIC_ANION_TRANSPORT | WT-TKO_up | 35 | 574 | 246 | 8100 | 2.00773349 | 5.19E-05 | 0.01320803 | Abat;Abcb1a;Abcc2;Abcc3;Cln8;Cyp4f18;Irs2;Ldlr;Lrrc8a;Per2;Plekha8;Plscr1;Pparg;Prkaa2;Rxra;Slc13a1;Slc13a2;Slc16a5;Slc23a1;Slc25a22;Slc2a2;Slc2a8;Slc37a4;Slc3a1;Slc4a11;Slc4a7; |

| System | MSigDB | Contrast | Overlap Size | Contrast Size | MSigDB. Size | Background Size | FE | Pvalue | P.adj | Overlap.Items |
|---|---|---|---|---|---|---|---|---|---|---|
| C5.BP | GO_ORGANIC_HYDROXY_COMPOUND_METABOLIC_PROCESS | WT-TKO_up | 45 | 574 | 261 | 8100 | 2.43301694 | 1.55E-08 | 0.00018908 | Abat;Acaa1a;Acox2;Acss1;Bco1;Bmp2;Ces1b;Cln8;Cyp4f18;Dgat2;Epas1;Fdft1;Fdps;Fdx1;Galk1;Gch1;Got1;Gpd1;Gpd2;Hagh;Hmgcs1;Idh1;Inpp5a;Insig1;Ldlr;Lss;Maoa;Maob;Msmo1;Mvk;Naaa;Nr1d1;Nsdhl;Osbpl3;Osbpl6;Per2;Plcd1;Pmel;Por;Prkaa2;Qdpr;Rxra;Srd5a2;Srebf2;Sult2b1 |
| C5.MF | GO_ACTIVE_ION_TRANSMEMBRANE_TRANSPORTER_ACTIVITY | WT-TKO_up | 24 | 574 | 107 | 8100 | 3.16519587 | 3.00E-07 | 0.00127404 | Abcc3;Atp1a1;Atp1b1;Atp2a3;Atp2b1;Clcn5;Slc13a1;Slc13a2;Slc15a1;Slc20a1;Slc23a1;Slc26a11;Slc2a10;Slc2a8;Slc37a4;Slc4a11;Slc4a7;Slc5a1;Slc5a4a;Slc5a6;Slc6a19;Slc6a4;Slc6a6;Slc6a8 |
| C5.BP | GO_LIPID_METABOLIC_PROCESS | WT-TKO_up | 81 | 574 | 660 | 8100 | 1.7318657 | 3.94E-07 | 0.00127404 | Abhd1;Abhd2;Abhd5;Acaa1a;Acox2;Acsm5;Acss1;Atp1a1;B4galt5;B4galt6;Bco1;Bmp2;Capn2;Ces1b;Ces2a;Ces2c;Cish;Cln8;Cwh43;Cyp2j6;Cyp2s1;Cyp2u1;Cyp4f18;Dgat2;Dgkd;Elovl1;Enpp7;Fdft1;Fdps;Fdx1;Gpd1;Hmgcs1;Hpgd;Hsd17b13;Hsd3b2;Hsd3b3;Idh1;Inpp5a;Insig1;Irs2;Klf4;Ldlr;Lpin3;Lss;Msmo1;Mvk;Naaa;Neu1;Nr1d1;Nr1d2;Nr1h3;Nsdhl;Osbpl3;Osbpl6;Per2;Phospho1;Pisd;Plbd2;Plcd1;Plekha8;Plscr1;Pm20d1;Por;Pparg;Prkaa2;Rb1;Rorc;Rxra;Sdc1;Sdc3;Srd5a2;Srebf2;St8sia6;Sult2b1;Tfcp2l1;Tnfrsf21;Trib3;Ugcg;Ugt2b5;Wdtc1;Zbtb20 |
| C5.BP | GO_RESPONSE_TO_ENDOGENOUS_STIMULUS | WT-TKO_up | 87 | 574 | 728 | 8100 | 1.68640158 | 4.48E-07 | 0.00127404 | Abcc2;Abhd2;Actb;Adcy2;Anxa2;Anxa3;Arnt2;Atp1a1;Atp2b1;Bace1;Bcar1;Bmp2;Capn2;Cited2;Cldn4;Cpn1;Cry2;Cx3cr1;Eef2k;Endog;Ezr;Fdx1;Fgfbp3;Fgfrl1;Flcn;Foxo3;Fzd1;Garem1;Gclc;Gcnt2;Got1;Gpd1;Hmgcs1;Hpgd;Hsd3b2;Hsd3b3;Idh1;Irs2;Itpr2;Jup;Klf15;Klf4;Klf9;Lats2;Ldlr;Lpin3;Maob;Myo1c;Nbl1;Nptn;Nr1d1;Nr1d2;Nr1h3;Nr3c2;P2ry2;Per1;Pmepa1;Por;Pparg;Prkaa2;Prnp;Qdpr;Rab31;Rapgef1;Rb1;Rorc;Runx1;Rxra;Sdc1;Sh3gl2;Slc2a8;Slc6a4;Slc9a3r1;Smad7;Spint1;Spred2;Spry2;Srebf2;Stxbp1;Trib3;Ubc;Ugcg;Vamp2;Wbp2;Wdtc1;Zfp703;Zyx |
| C2.CP | REACTOME_BIOLOGICAL_OXIDATIONS | WT-TKO_up | 24 | 574 | 111 | 8100 | 3.05113476 | 6.16E-07 | 0.00127404 | Acsm5;Acss1;Acy3;Arnt2;Ces1b;Ces2a;Ces2c;Cndp2;Cyp2j6;Cyp2s1;Cyp2u1;Cyp4b1;Cyp4f18;Dpep1;Fdx1;Gclc;Gstm2;Maoa;Maob;Oplah;Por;Rxra;Sult2b1;Ugt2b5 |
| C5.BP | GO_STEROID_METABOLIC_PROCESS | WT-TKO_up | 31 | 574 | 168 | 8100 | 2.60390742 | 6.26E-07 | 0.00127404 | Acaa1a;Acox2;Atp1a1;Bmp2;Ces1b;Cln8;Dgat2;Fdft1;Fdps;Fdx1;Hmgcs1;Hsd3b2;Hsd3b3;Insig1;Ldlr;Lss;Msmo1;Mvk;Nr1d1;Nsdhl;Osbpl3;Osbpl6;Por;Prkaa2;Rorc;Rxra;Srd5a2;Srebf2;Sult2b1;Tfcp2l1;Ugt2b5 |
| C2.CP | REACTOME_METABOLISM_OF_STEROIDS | WT-TKO_up | 18 | 574 | 70 | 8100 | 3.62867098 | 1.11E-06 | 0.00169923 | Abcc3;Acox2;Fdft1;Fdps;Fdx1;Hmgcs1;Hsd3b2;Hsd3b3;Insig1;Lss;Msmo1;Mvk;Nsdhl;Osbpl3;Osbpl6;Rxra;Srd5a2;Srebf2 |
| C5.MF | GO_SYMPORTER_ACTIVITY | WT-TKO_up | 18 | 574 | 70 | 8100 | 3.62867098 | 1.11E-06 | 0.00169923 | Slc13a1;Slc13a2;Slc15a1;Slc16a5;Slc20a1;Slc23a1;Slc25a22;Slc2a10;Slc2a8;Slc4a11;Slc4a7;Slc5a1;Slc5a4a;Slc5a6;Slc6a19;Slc6a4;Slc6a6;Slc6a8 |
| C5.BP | GO_LIPID_BIOSYNTHETIC_PROCESS | WT-TKO_up | 50 | 574 | 352 | 8100 | 2.00447418 | 1.33E-06 | 0.00175947 | Abhd5;Acox2;Acsm5;Acss1;Atp1a1;B4galt5;B4galt6;Bco1;Bmp2;Capn2;Ces1b;Cln8;Cwh43;Dgat2;Elovl1;Fdft1;Fdps;Fdx1;Gpd1;Hmgcs1;Hpgd;Hsd17b13;Hsd3b2;Hsd3b3;Idh1;Insig1;Ldlr;Lpin3;Lss;Msmo1;Mvk;Nr1d1;Nr1h3;Nsdhl;Osbpl3;Osbpl6;Phospho1;Pisd;Plekha8;Plscr1;Por;Prkaa2;Srd5a2;Srebf2;St8sia6;Tfcp2l1;Trib3;Ugcg;Wdtc1;Zbtb20 |
| C5.BP | GO_ORGANIC_ACID_METABOLIC_PROCESS | WT-TKO_up | 68 | 574 | 541 | 8100 | 1.77371882 | 1.61E-06 | 0.00175947 | Abat;Abcc2;Abhd5;Acaa1a;Acox2;Acsm5;Acss1;Amdhd2;Azin1;Azin2;B4galt5;B4galt6;Btd;Ces1b;Ces2a;Ces2c;Chp1;Cyp2j6;Cyp2s1;Cyp2u1;Cyp4f18;Dgat2;Dpep1;Elovl1;Fbp1;Galk1;Gch1;Gclc;Glud1;Gnpda1;Gnpda2;Got1;Gpd1;Hagh;Hkdc1;Hoga1;Hpgd;Hykk;Idh1;Il15;Insig1;Irs2;Lpin3;Me2;Me3;Msmo1;Naaa;Npl;Nr1d1;Nr1h3;Osbpl3;Osbpl6;Per2;Pgm2;Pm20d1;Por;Pparg;Prkaa2;Qdpr;Sardh;Slc23a1;Slc5a6;Slc6a8;Sult2b1;Trib3;Ugt2b5;Wdtc1;Zbtb20 |
| C5.BP | GO_SMALL_MOLECULE_METABOLIC_PROCESS | WT-TKO_up | 100 | 574 | 899 | 8100 | 1.56968835 | 1.69E-06 | 0.00175947 | Abat;Abcc2;Abhd5;Acaa1a;Acot9;Acox2;Acsm5;Acss1;Adcy2;Amdhd2;Azin1;Azin2;B4galt5;B4galt6;Bco1;Bmp2;Btd;Ces1b;Ces2a;Ces2c;Chp1;Cln8;Coq10b;Cyp2j6;Cyp2s1;Cyp2u1;Cyp4f18;Dgat2;Dpep1;Efl1;Elovl1;Entpd7;Fbp1;Fdft1;Fdps;Fdx1;Flcn;Galk1;Gch1;Gclc;Glud1;Gnpda1;Gnpda2;Got1;Gpd1;Gpd2;Guca2a;Hagh;Hkdc1;Hmgcs1;Hoga1;Hpgd;Hykk;Idh1;Il15;Inpp5a;Insig1;Irs2;Ldlr;Lpin3;Lss;Me2;Me3;Mpp1;Msmo1;Mvk;Naaa;Nmrk1;Npl;Nr1d1;Nr1h3;Nsdhl;Osbpl3;Osbpl6;Pank3;Pde10a;Per2;Pgm2;Plcd1;Pm20d1;Por;Pparg;Prkaa2;Qdpr;Rnaseh2b;Rorc;Rxra;Sardh;Slc23a1;Slc37a4;Slc5a6;Slc6a8;Smpdl3a;Srd5a2;Srebf2;Sult2b1;Trib3;Ugt2b5;Wdtc1;Zbtb20 |
| C5.BP | GO_ANION_TRANSPORT | WT-TKO_up | 44 | 574 | 296 | 8100 | 2.09765515 | 1.73E-06 | 0.00175947 | Abat;Abcb1a;Abcc2;Abcc3;Clca4a;Clcn5;Cldn4;Clic1;Cln8;Cry2;Cyp4f18;Irs2;Ldlr;Lrrc8a;Per2;Plekha8;Plscr1;Pparg;Prkaa2;Prnp;Rxra;Slc13a1;Slc13a2;Slc16a5;Slc20a1;Slc23a1;Slc25a22;Slc26a11;Slc2a2;Slc2a8;Slc37a4;Slc3a1;Slc4a11;Slc4a7;Slc5a6;Slc6a19;Slc6a6;Slc6a8;Slc7a8;Slc9a3r1;Stxbp1;Ttyh3;Vamp2;Xkr9 |
| C5.BP | GO_STEROID_BIOSYNTHETIC_PROCESS | WT-TKO_up | 23 | 574 | 110 | 8100 | 2.950586 | 1.96E-06 | 0.0018001 | Acox2;Atp1a1;Bmp2;Ces1b;Fdft1;Fdps;Fdx1;Hmgcs1;Hsd3b2;Hsd3b3;Insig1;Lss;Msmo1;Mvk;Nr1d1;Nsdhl;Osbpl3;Osbpl6;Por;Prkaa2;Srd5a2;Srebf2;Tfcp2l1 |
| C5.MF | GO_ACTIVE_TRANSMEMBRANE_TRANSPORTER_ACTIVITY | WT-TKO_up | 29 | 574 | 160 | 8100 | 2.55770906 | 2.11E-06 | 0.0018001 | Abca8b;Abcb1a;Abcc2;Abcc3;Atp1a1;Atp1b1;Atp2a3;Atp2b1;Clcn5;Slc13a1;Slc13a2;Slc15a1;Slc16a5;Slc20a1;Slc23a1;Slc25a22;Slc26a11;Slc2a10;Slc2a8;Slc37a4;Slc4a11;Slc4a7;Slc5a1;Slc5a4a;Slc5a6;Slc6a19;Slc6a4;Slc6a6;Slc6a8 |
| C5.MF | GO_SOLUTE_CATION_SYMPORTER_ACTIVITY | WT-TKO_up | 15 | 574 | 53 | 8100 | 3.99382026 | 2.38E-06 | 0.0018001 | Slc13a1;Slc13a2;Slc15a1;Slc20a1;Slc23a1;Slc2a10;Slc2a8;Slc4a7;Slc5a1;Slc5a4a;Slc5a6;Slc6a19;Slc6a4;Slc6a6;Slc6a8 |
| C5.BP | GO_MONOCARBOXYLIC_ACID_METABOLIC_PROCESS | WT-TKO_up | 47 | 574 | 330 | 8100 | 2.00981945 | 2.60E-06 | 0.0018001 | Abat;Abcc2;Abhd5;Acaa1a;Acox2;Acsm5;Acss1;Btd;Ces1b;Ces2a;Ces2c;Cyp2j6;Cyp2s1;Cyp2u1;Cyp4f18;Dgat2;Elovl1;Fbp1;Galk1;Gpd1;Hagh;Hkdc1;Hoga1;Hpgd;Idh1;Insig1;Irs2;Lpin3;Me2;Me3;Msmo1;Naaa;Nr1d1;Nr1h3;Osbpl3;Osbpl6;Per2;Pgm2;Por;Pparg;Prkaa2;Slc5a6;Slc6a8;Trib3;Ugt2b5;Wdtc1;Zbtb20 |
| C5.MF | GO_SOLUTE_SODIUM_SYMPORTER_ACTIVITY | WT-TKO_up | 12 | 574 | 35 | 8100 | 4.83822797 | 2.63E-06 | 0.0018001 | Slc13a1;Slc13a2;Slc20a1;Slc23a1;Slc4a7;Slc5a1;Slc5a4a;Slc5a6;Slc6a19;Slc6a4;Slc6a6;Slc6a8 |
| C5.BP | GO_TRANSMEMBRANE_TRANSPORT | WT-TKO_up | 72 | 574 | 592 | 8100 | 1.7162633 | 2.65E-06 | 0.0018001 | Abca8b;Abcb1a;Abcc2;Abcc3;Actb;Ank3;Anxa2;Atp1a1;Atp1b1;Atp2a3;Atp2b1;Azin1;Azin2;Bcr;Bin1;Chp1;Clca4a;Clcn5;Cldn4;Clic1;Cln8;Cnnm1;Gjb3;Hap1;Irs2;Itpr2;Klf15;Lrrc8a;Mcoln1;Nipal1;Otop2;Per2;Plcd1;Pm20d1;Prkaa2;Prnp;S100a6;Scnn1a;Sgk3;Slc13a1;Slc13a2;Slc15a1;Slc16a5;Slc20a1;Slc23a1;Slc25a22;Slc26a11;Slc2a10;Slc2a2;Slc2a5;Slc2a8;Slc30a2;Slc37a4;Slc3a1;Slc4a11;Slc4a7;Slc5a1;Slc5a4a;Slc5a6;Slc6a19;Slc6a4;Slc6a6;Slc6a8;Slc7a8;Slc9a3r1;Tmc7;Tmem144;Tmem63b;Trib3;Ttyh3;Ubc;Vamp2 |
| C5.BP | GO_ION_TRANSPORT | WT-TKO_up | 79 | 574 | 673 | 8100 | 1.65647602 | 3.32E-06 | 0.00206903 | Abat;Abcb1a;Abcc2;Abcc3;Ank3;Anxa2;Atp1a1;Atp1b1;Atp2a3;Atp2b1;Bin1;Chp1;Clca4a;Clcn5;Cldn4;Clic1;Cln8;Cnnm1;Cry2;Cyp4f18;Gja4;Hap1;Irs2;Itpr2;Ldlr;Lgals3;Lpar3;Lrrc8a;Maob;Mcoln1;Nipal1;Otop2;Per1;Per2;Plcd1;Plekha8;Plscr1;Pm20d1;Pparg;Prkaa2;Prnp;Pxk;Rxra;S100a6;Scnn1a;Sgk3;Slc13a1;Slc13a2;Slc15a1;Slc16a5;Slc20a1;Slc23a1;Slc25a22;Slc26a11;Slc2a10;Slc2a2;Slc2a8;Slc30a2;Slc37a4;Slc3a1;Slc4a11;Slc4a7;Slc5a1;Slc5a4a;Slc5a6;Slc6a19;Slc6a4;Slc6a6;Slc6a8;Slc7a8;Slc9a3r1;Stxbp1;Syt12;Syt7;Tmc7;Tmem63b;Ttyh3;Vamp2;Xkr9 |
| C5.BP | GO_ANION_TRANSMEMBRANE_TRANSPORT | WT-TKO_up | 26 | 574 | 138 | 8100 | 2.65868808 | 3.39E-06 | 0.00206903 | Abcb1a;Clca4a;Clcn5;Cldn4;Clic1;Cln8;Irs2;Lrrc8a;Per2;Prkaa2;Prnp;Slc13a1;Slc20a1;Slc23a1;Slc25a22;Slc26a11;Slc37a4;Slc3a1;Slc4a11;Slc4a7;Slc5a6;Slc6a19;Slc6a6;Slc6a8;Slc7a8;Ttyh3 |
| C5.BP | GO_REGULATION_OF_CIRCADIAN_RHYTHM | WT-TKO_up | 13 | 574 | 42 | 8100 | 4.3678447 | 3.73E-06 | 0.00217022 | Bhlhe40;Cipc;Cry2;Nr1d1;Nr1d2;Nr1h3;Per1;Per2;Per3;Pparg;Prkaa2;Rorc;Usp2 |

| System | MSigDB | Contrast | Overlap Size | Contrast Size | MSigDB. Size | Background Size | FE | Pvalue | P.adj | Overlap.Items |
|---|---|---|---|---|---|---|---|---|---|---|
| C5.BP | GO_ORGANIC_HYDROXY_COMPOUND_METABOLIC_PROCESS | WT-TKO_up | 45 | 574 | 261 | 8100 | 2.43301694 | 1.55E-08 | 0.00018908 | Abat;Acaa1a;Acox2;Acss1;Bco1;Bmp2;Ces1b;Cln8;Cyp4f18;Dgat2;Epas1;Fdft1;Fdps;Fdx1;Galk1;Gch1;Got1;Gpd1;Gpd2;Hagh;Hmgcs1;Idh1;Inpp5a;Insig1;Ldlr;Lss;Maoa;Maob;Msmo1;Mvk;Naaa;Nr1d1;Nsdhl;Osbpl3;Osbpl6;Per2;Plcd1;Pmel;Por;Prkaa2;Qdpr;Rxra;Srd5a2;Srebf2;Sult2b1 |
| C5.MF | GO_ACTIVE_ION_TRANSMEMBRANE_TRANSPORTER_ACTIVITY | WT-TKO_up | 24 | 574 | 107 | 8100 | 3.16519587 | 3.00E-07 | 0.00127404 | Abcc3;Atp1a1;Atp1b1;Atp2a3;Atp2b1;Clcn5;Slc13a1;Slc13a2;Slc15a1;Slc20a1;Slc23a1;Slc26a11;Slc2a10;Slc2a8;Slc37a4;Slc4a11;Slc4a7;Slc5a1;Slc5a4a;Slc5a6;Slc6a19;Slc6a4;Slc6a6;Slc6a8 |
| C5.BP | GO_LIPID_METABOLIC_PROCESS | WT-TKO_up | 81 | 574 | 660 | 8100 | 1.7318657 | 3.94E-07 | 0.00127404 | Abhd1;Abhd2;Abhd5;Acaa1a;Acox2;Acsm5;Acss1;Atp1a1;B4galt5;B4galt6;Bco1;Bmp2;Capn2;Ces1b;Ces2a;Ces2c;Cish;Cln8;Cwh43;Cyp2j6;Cyp2s1;Cyp2u1;Cyp4f18;Dgat2;Dgkd;Elovl1;Enpp7;Fdft1;Fdps;Fdx1;Gpd1;Hmgcs1;Hpgd;Hsd17b13;Hsd3b2;Hsd3b3;Idh1;Inpp5a;Insig1;Irs2;Klf4;Ldlr;Lpin3;Lss;Msmo1;Mvk;Naaa;Neu1;Nr1d1;Nr1d2;Nr1h3;Nsdhl;Osbpl3;Osbpl6;Per2;Phospho1;Pisd;Plbd2;Plcd1;Plekha8;Plscr1;Pm20d1;Por;Pparg;Prkaa2;Rb1;Rorc;Rxra;Sdc1;Sdc3;Srd5a2;Srebf2;St8sia6;Sult2b1;Tfcp2l1;Tnfrsf21;Trib3;Ugcg;Ugt2b5;Wdtc1;Zbtb20 |
| C5.BP | GO_RESPONSE_TO_ENDOGENOUS_STIMULUS | WT-TKO_up | 87 | 574 | 728 | 8100 | 1.68640158 | 4.48E-07 | 0.00127404 | Abcc2;Abhd2;Actb;Adcy2;Anxa2;Anxa3;Arnt2;Atp1a1;Atp2b1;Bace1;Bcar1;Bmp2;Capn2;Cited2;Cldn4;Cpn1;Cry2;Cx3cr1;Eef2k;Endog;Ezr;Fdx1;Fgfbp3;Fgfrl1;Flcn;Foxo3;Fzd1;Garem1;Gclc;Gcnt2;Got1;Gpd1;Hmgcs1;Hpgd;Hsd3b2;Hsd3b3;Idh1;Irs2;Itpr2;Jup;Klf15;Klf4;Klf9;Lats2;Ldlr;Lpin3;Maob;Myo1c;Nbl1;Nptn;Nr1d1;Nr1d2;Nr1h3;Nr3c2;P2ry2;Per1;Pmepa1;Por;Pparg;Prkaa2;Prnp;Qdpr;Rab31;Rapgef1;Rb1;Rorc;Runx1;Rxra;Sdc1;Sh3gl2;Slc2a8;Slc6a4;Slc9a3r1;Smad7;Spint1;Spred2;Spry2;Srebf2;Stxbp1;Trib3;Ubc;Ugcg;Vamp2;Wbp2;Wdtc1;Zfp703;Zyx |
| C2.CP | REACTOME_BIOLOGICAL_OXIDATIONS | WT-TKO_up | 24 | 574 | 111 | 8100 | 3.05113476 | 6.16E-07 | 0.00127404 | Acsm5;Acss1;Acy3;Arnt2;Ces1b;Ces2a;Ces2c;Cndp2;Cyp2j6;Cyp2s1;Cyp2u1;Cyp4b1;Cyp4f18;Dpep1;Fdx1;Gclc;Gstm2;Maoa;Maob;Oplah;Por;Rxra;Sult2b1;Ugt2b5 |
| C5.BP | GO_STEROID_METABOLIC_PROCESS | WT-TKO_up | 31 | 574 | 168 | 8100 | 2.60390742 | 6.26E-07 | 0.00127404 | Acaa1a;Acox2;Atp1a1;Bmp2;Ces1b;Cln8;Dgat2;Fdft1;Fdps;Fdx1;Hmgcs1;Hsd3b2;Hsd3b3;Insig1;Ldlr;Lss;Msmo1;Mvk;Nr1d1;Nsdhl;Osbpl3;Osbpl6;Por;Prkaa2;Rorc;Rxra;Srd5a2;Srebf2;Sult2b1;Tfcp2l1;Ugt2b5 |
| C2.CP | REACTOME_METABOLISM_OF_STEROIDS | WT-TKO_up | 18 | 574 | 70 | 8100 | 3.62867098 | 1.11E-06 | 0.00169923 | Abcc3;Acox2;Fdft1;Fdps;Fdx1;Hmgcs1;Hsd3b2;Hsd3b3;Insig1;Lss;Msmo1;Mvk;Nsdhl;Osbpl3;Osbpl6;Rxra;Srd5a2;Srebf2 |
| C5.MF | GO_SYMPORTER_ACTIVITY | WT-TKO_up | 18 | 574 | 70 | 8100 | 3.62867098 | 1.11E-06 | 0.00169923 | Slc13a1;Slc13a2;Slc15a1;Slc16a5;Slc20a1;Slc23a1;Slc25a22;Slc2a10;Slc2a8;Slc4a11;Slc4a7;Slc5a1;Slc5a4a;Slc5a6;Slc6a19;Slc6a4;Slc6a6;Slc6a8 |
| C5.BP | GO_LIPID_BIOSYNTHETIC_PROCESS | WT-TKO_up | 50 | 574 | 352 | 8100 | 2.00447418 | 1.33E-06 | 0.00175947 | Abhd5;Acox2;Acsm5;Acss1;Atp1a1;B4galt5;B4galt6;Bco1;Bmp2;Capn2;Ces1b;Cln8;Cwh43;Dgat2;Elovl1;Fdft1;Fdps;Fdx1;Gpd1;Hmgcs1;Hpgd;Hsd17b13;Hsd3b2;Hsd3b3;Idh1;Insig1;Ldlr;Lpin3;Lss;Msmo1;Mvk;Nr1d1;Nr1h3;Nsdhl;Osbpl3;Osbpl6;Phospho1;Pisd;Plekha8;Plscr1;Por;Prkaa2;Srd5a2;Srebf2;St8sia6;Tfcp2l1;Trib3;Ugcg;Wdtc1;Zbtb20 |
| C5.BP | GO_ORGANIC_ACID_METABOLIC_PROCESS | WT-TKO_up | 68 | 574 | 541 | 8100 | 1.77371882 | 1.61E-06 | 0.00175947 | Abat;Abcc2;Abhd5;Acaa1a;Acox2;Acsm5;Acss1;Amdhd2;Azin1;Azin2;B4galt5;B4galt6;Btd;Ces1b;Ces2a;Ces2c;Chp1;Cyp2j6;Cyp2s1;Cyp2u1;Cyp4f18;Dgat2;Dpep1;Elovl1;Fbp1;Galk1;Gch1;Gclc;Glud1;Gnpda1;Gnpda2;Got1;Gpd1;Hagh;Hkdc1;Hoga1;Hpgd;Hykk;Idh1;Il15;Insig1;Irs2;Lpin3;Me2;Me3;Msmo1;Naaa;Npl;Nr1d1;Nr1h3;Osbpl3;Osbpl6;Per2;Pgm2;Pm20d1;Por;Pparg;Prkaa2;Qdpr;Sardh;Slc23a1;Slc5a6;Slc6a8;Sult2b1;Trib3;Ugt2b5;Wdtc1;Zbtb20 |
| C5.BP | GO_SMALL_MOLECULE_METABOLIC_PROCESS | WT-TKO_up | 100 | 574 | 899 | 8100 | 1.56968835 | 1.69E-06 | 0.00175947 | Abat;Abcc2;Abhd5;Acaa1a;Acot9;Acox2;Acsm5;Acss1;Adcy2;Amdhd2;Azin1;Azin2;B4galt5;B4galt6;Bco1;Bmp2;Btd;Ces1b;Ces2a;Ces2c;Chp1;Cln8;Coq10b;Cyp2j6;Cyp2s1;Cyp2u1;Cyp4f18;Dgat2;Dpep1;Efl1;Elovl1;Entpd7;Fbp1;Fdft1;Fdps;Fdx1;Flcn;Galk1;Gch1;Gclc;Glud1;Gnpda1;Gnpda2;Got1;Gpd1;Gpd2;Guca2a;Hagh;Hkdc1;Hmgcs1;Hoga1;Hpgd;Hykk;Idh1;Il15;Inpp5a;Insig1;Irs2;Ldlr;Lpin3;Lss;Me2;Me3;Mpp1;Msmo1;Mvk;Naaa;Nmrk1;Npl;Nr1d1;Nr1h3;Nsdhl;Osbpl3;Osbpl6;Pank3;Pde10a;Per2;Pgm2;Plcd1;Pm20d1;Por;Pparg;Prkaa2;Qdpr;Rnaseh2b;Rorc;Rxra;Sardh;Slc23a1;Slc37a4;Slc5a6;Slc6a8;Smpdl3a;Srd5a2;Srebf2;Sult2b1;Trib3;Ugt2b5;Wdtc1;Zbtb20 |
| C5.BP | GO_ANION_TRANSPORT | WT-TKO_up | 44 | 574 | 296 | 8100 | 2.09765515 | 1.73E-06 | 0.00175947 | Abat;Abcb1a;Abcc2;Abcc3;Clca4a;Clcn5;Cldn4;Clic1;Cln8;Cry2;Cyp4f18;Irs2;Ldlr;Lrrc8a;Per2;Plekha8;Plscr1;Pparg;Prkaa2;Prnp;Rxra;Slc13a1;Slc13a2;Slc16a5;Slc20a1;Slc23a1;Slc25a22;Slc26a11;Slc2a2;Slc2a8;Slc37a4;Slc3a1;Slc4a11;Slc4a7;Slc5a6;Slc6a19;Slc6a6;Slc6a8;Slc7a8;Slc9a3r1;Stxbp1;Ttyh3;Vamp2;Xkr9 |
| C5.BP | GO_STEROID_BIOSYNTHETIC_PROCESS | WT-TKO_up | 23 | 574 | 110 | 8100 | 2.950586 | 1.96E-06 | 0.0018001 | Acox2;Atp1a1;Bmp2;Ces1b;Fdft1;Fdps;Fdx1;Hmgcs1;Hsd3b2;Hsd3b3;Insig1;Lss;Msmo1;Mvk;Nr1d1;Nsdhl;Osbpl3;Osbpl6;Por;Prkaa2;Srd5a2;Srebf2;Tfcp2l1 |
| C5.MF | GO_ACTIVE_TRANSMEMBRANE_TRANSPORTER_ACTIVITY | WT-TKO_up | 29 | 574 | 160 | 8100 | 2.55770906 | 2.11E-06 | 0.0018001 | Abca8b;Abcb1a;Abcc2;Abcc3;Atp1a1;Atp1b1;Atp2a3;Atp2b1;Clcn5;Slc13a1;Slc13a2;Slc15a1;Slc16a5;Slc20a1;Slc23a1;Slc25a22;Slc26a11;Slc2a10;Slc2a8;Slc37a4;Slc4a11;Slc4a7;Slc5a1;Slc5a4a;Slc5a6;Slc6a19;Slc6a4;Slc6a6;Slc6a8 |
| C5.MF | GO_SOLUTE_CATION_SYMPORTER_ACTIVITY | WT-TKO_up | 15 | 574 | 53 | 8100 | 3.99382026 | 2.38E-06 | 0.0018001 | Slc13a1;Slc13a2;Slc15a1;Slc20a1;Slc23a1;Slc2a10;Slc2a8;Slc4a7;Slc5a1;Slc5a4a;Slc5a6;Slc6a19;Slc6a4;Slc6a6;Slc6a8 |
| C5.BP | GO_MONOCARBOXYLIC_ACID_METABOLIC_PROCESS | WT-TKO_up | 47 | 574 | 330 | 8100 | 2.00981945 | 2.60E-06 | 0.0018001 | Abat;Abcc2;Abhd5;Acaa1a;Acox2;Acsm5;Acss1;Btd;Ces1b;Ces2a;Ces2c;Cyp2j6;Cyp2s1;Cyp2u1;Cyp4f18;Dgat2;Elovl1;Fbp1;Galk1;Gpd1;Hagh;Hkdc1;Hoga1;Hpgd;Idh1;Insig1;Irs2;Lpin3;Me2;Me3;Msmo1;Naaa;Nr1d1;Nr1h3;Osbpl3;Osbpl6;Per2;Pgm2;Por;Pparg;Prkaa2;Slc5a6;Slc6a8;Trib3;Ugt2b5;Wdtc1;Zbtb20 |
| C5.MF | GO_SOLUTE_SODIUM_SYMPORTER_ACTIVITY | WT-TKO_up | 12 | 574 | 35 | 8100 | 4.83822797 | 2.63E-06 | 0.0018001 | Slc13a1;Slc13a2;Slc20a1;Slc23a1;Slc4a7;Slc5a1;Slc5a4a;Slc5a6;Slc6a19;Slc6a4;Slc6a6;Slc6a8 |
| C5.BP | GO_TRANSMEMBRANE_TRANSPORT | WT-TKO_up | 72 | 574 | 592 | 8100 | 1.7162633 | 2.65E-06 | 0.0018001 | Abca8b;Abcb1a;Abcc2;Abcc3;Actb;Ank3;Anxa2;Atp1a1;Atp1b1;Atp2a3;Atp2b1;Azin1;Azin2;Bcr;Bin1;Chp1;Clca4a;Clcn5;Cldn4;Clic1;Cln8;Cnnm1;Gjb3;Hap1;Irs2;Itpr2;Klf15;Lrrc8a;Mcoln1;Nipal1;Otop2;Per2;Plcd1;Pm20d1;Prkaa2;Prnp;S100a6;Scnn1a;Sgk3;Slc13a1;Slc13a2;Slc15a1;Slc16a5;Slc20a1;Slc23a1;Slc25a22;Slc26a11;Slc2a10;Slc2a2;Slc2a5;Slc2a8;Slc30a2;Slc37a4;Slc3a1;Slc4a11;Slc4a7;Slc5a1;Slc5a4a;Slc5a6;Slc6a19;Slc6a4;Slc6a6;Slc6a8;Slc7a8;Slc9a3r1;Tmc7;Tmem144;Tmem63b;Trib3;Ttyh3;Ubc;Vamp2 |
| C5.BP | GO_ION_TRANSPORT | WT-TKO_up | 79 | 574 | 673 | 8100 | 1.65647602 | 3.32E-06 | 0.00206903 | Abat;Abcb1a;Abcc2;Abcc3;Ank3;Anxa2;Atp1a1;Atp1b1;Atp2a3;Atp2b1;Bin1;Chp1;Clca4a;Clcn5;Cldn4;Clic1;Cln8;Cnnm1;Cry2;Cyp4f18;Gja4;Hap1;Irs2;Itpr2;Ldlr;Lgals3;Lpar3;Lrrc8a;Maob;Mcoln1;Nipal1;Otop2;Per1;Per2;Plcd1;Plekha8;Plscr1;Pm20d1;Pparg;Prkaa2;Prnp;Pxk;Rxra;S100a6;Scnn1a;Sgk3;Slc13a1;Slc13a2;Slc15a1;Slc16a5;Slc20a1;Slc23a1;Slc25a22;Slc26a11;Slc2a10;Slc2a2;Slc2a8;Slc30a2;Slc37a4;Slc3a1;Slc4a11;Slc4a7;Slc5a1;Slc5a4a;Slc5a6;Slc6a19;Slc6a4;Slc6a6;Slc6a8;Slc7a8;Slc9a3r1;Stxbp1;Syt12;Syt7;Tmc7;Tmem63b;Ttyh3;Vamp2;Xkr9 |
| C5.BP | GO_ANION_TRANSMEMBRANE_TRANSPORT | WT-TKO_up | 26 | 574 | 138 | 8100 | 2.65868808 | 3.39E-06 | 0.00206903 | Abcb1a;Clca4a;Clcn5;Cldn4;Clic1;Cln8;Irs2;Lrrc8a;Per2;Prkaa2;Prnp;Slc13a1;Slc20a1;Slc23a1;Slc25a22;Slc26a11;Slc37a4;Slc3a1;Slc4a11;Slc4a7;Slc5a6;Slc6a19;Slc6a6;Slc6a8;Slc7a8;Ttyh3 |
| C5.BP | GO_REGULATION_OF_CIRCADIAN_RHYTHM | WT-TKO_up | 13 | 574 | 42 | 8100 | 4.3678447 | 3.73E-06 | 0.00217022 | Bhlhe40;Cipc;Cry2;Nr1d1;Nr1d2;Nr1h3;Per1;Per2;Per3;Pparg;Prkaa2;Rorc;Usp2 |

| | | | | | | | | | | |
|---|---|---|---|---|---|---|---|---|---|---|
| C2.CP | REACTOME_CYTOCHROME_P450_ARRANGED_BY_SUBSTRATE_TYPE | WT-TKO_up | 9 | 574 | 27 | 8100 | 4.70383275 | 6.23E-05 | 0.01522397 | Arnt2;Cyp2j6;Cyp2s1;Cyp2u1;Cyp4b1;Cyp4f18;Fdx1;Por;Rxra |
| C5.BP | GO_INORGANIC_ANION_TRANSPORT | WT-TKO_up | 15 | 574 | 68 | 8100 | 3.1128305 | 6.23E-05 | 0.01522397 | Abcb1a;Clca4a;Clcn5;Cldn4;Clic1;Cry2;Lrrc8a;Prnp;Slc13a1;Slc20a1;Slc26a11;Slc37a4;Slc4a11;Slc4a7;Ttyh3 |
| C5.BP | GO_RESPONSE_TO_ORGANIC_CYCLIC_COMPOUND | WT-TKO_up | 53 | 574 | 439 | 8100 | 1.70366608 | 7.43E-05 | 0.01771354 | Abat;Abcc2;Abhd2;Actb;Adcy2;Anxa3;Anxa7;Arnt2;Atp1a1;Atp2b1;Bmp2;Cldn4;Cpn1;Cry2;Eef2k;Endog;Ezr;Fdx1;Gclc;Got1;Gpd1;Hmgcs1;Hpgd;Hsd3b2;Hsd3b3;Idh1;Il15;Insig1;Itpr2;Jup;Klf4;Klf9;Maob;Nr1d1;Nr1d2;Nr1h3;Nr3c2;P2ry2;Per1;Pmepa1;Pparg;Ralb;Rapgef1;Rb1;Rorc;Runx1;Rxra;Sdc1;Slc6a4;Slc9a3r1;Stxbp1;Wbp2;Zfp703 |
| C5.BP | GO_REGULATION_OF_LIPID_METABOLIC_PROCESS | WT-TKO_up | 31 | 574 | 211 | 8100 | 2.0732533 | 7.54E-05 | 0.01771354 | Abhd5;Atp1a1;Bmp2;Capn2;Cish;Dgat2;Fdft1;Fdps;Hmgcs1;Hsd17b13;Idh1;Insig1;Irs2;Klf4;Ldlr;Lss;Mvk;Nr1d1;Nr1d2;Nr1h3;Por;Pparg;Prkaa2;Rb1;Rorc;Rxra;Srebf2;Tnfrsf21;Trib3;Wdtc1;Zbtb20 |
| C2.CP | KEGG_CIRCADIAN_RHYTHM_MAMMAL | WT-TKO_up | 6 | 574 | 12 | 8100 | 7.05574913 | 7.88E-05 | 0.0179765 | Bhlhe40;Cry2;Nr1d1;Per1;Per2;Per3 |
| C5.MF | GO_CARBOHYDRATE_TRANSMEMBRANE_TRANSPOR | WT-TKO_up | 8 | 574 | 22 | 8100 | 5.13145391 | 7.95E-05 | 0.0179765 | Slc23a1;Slc2a10;Slc2a2;Slc2a5;Slc2a8;Slc5a1;Slc5a4a;Tmem144 |
| C2.CP | BIOCARTA_ARENRF2_PATHWAY | WT-TKO_up | 5 | 574 | 8 | 8100 | 8.81968641 | 8.22E-05 | 0.01825291 | Cryz;Hmox1;Maff;Mafk;Por |
| C5.BP | GO_RESPONSE_TO_STEROID_HORMONE | WT-TKO_up | 27 | 574 | 174 | 8100 | 2.18971525 | 8.51E-05 | 0.01845686 | Abcc2;Abhd2;Anxa3;Atp1a1;Atp2b1;Cldn4;Cpn1;Cry2;Got1;Hsd3b2;Hsd3b3;Idh1;Klf9;Maob;Nr1d1;Nr1d2;Nr1h3;Nr3c2;Per1;Pmepa1;Pparg;Rb1;Rorc;Runx1;Rxra;Sdc1;Wbp2 |
| C5.MF | GO_CALCIUM_DEPENDENT_PROTEIN_BINDING | WT-TKO_up | 9 | 574 | 28 | 8100 | 4.53583873 | 8.61E-05 | 0.01845686 | Anxa2;Anxa3;Anxa7;Chp1;Masp1;Prnp;S100a6;Vamp2;Vps37b |
| C5.BP | GO_SODIUM_ION_TRANSPORT | WT-TKO_up | 17 | 574 | 86 | 8100 | 2.78948221 | 8.99E-05 | 0.0188772 | Ank3;Atp1a1;Atp1b1;Chp1;Per1;Scnn1a;Slc13a1;Slc13a2;Slc20a1;Slc23a1;Slc4a11;Slc4a7;Slc5a1;Slc5a4a;Slc5a6;Slc6a8;Slc9a3r1 |
| C5.BP | GO_CARBOHYDRATE_METABOLIC_PROCESS | WT-TKO_up | 37 | 574 | 273 | 8100 | 1.91254738 | 9.12E-05 | 0.0188772 | Amdhd2;B4galt5;B4galt6;Dgat2;Fbp1;Galk1;Gclc;Gcnt3;Gnpda1;Gnpda2;Got1;Gpd1;Gpd2;Hkdc1;Idh1;Inpp5a;Irs2;Mgam;Neu1;Npl;Nr1d1;Per2;Pgm2;Prkaa2;Rorc;Sis;Slc23a1;Slc2a2;Slc2a5;Slc2a8;Slc37a4;Slc3a1;St8sia6;Stk40;Ugt2b5;Wdtc1;Zbtb20 |
| C5.MF | GO_TRANSPORTER_ACTIVITY | WT-TKO_up | 54 | 574 | 455 | 8100 | 1.67477122 | 0.0001 | 0.02046143 | Abca8b;Abcb1a;Abcc2;Abcc3;Anxa2;Atp1a1;Atp1b1;Atp2a3;Atp2b1;Clca4a;Clcn5;Cldn4;Clic1;Cnnm1;Gjb3;Itpr2;Lrrc8a;Mcoln1;Nipal1;Otop2;Pacsin2;Plekha8;Plscr1;S100a6;Scnn1a;Slc13a1;Slc13a2;Slc15a1;Slc16a5;Slc20a1;Slc23a1;Slc25a22;Slc26a11;Slc2a10;Slc2a2;Slc2a5;Slc2a8;Slc30a2;Slc37a4;Slc3a1;Slc4a11;Slc4a7;Slc5a1;Slc5a4a;Slc5a6;Slc6a19;Slc6a4;Slc6a6;Slc6a8;Slc7a8;Tmc7;Tmem144;Tmem63b;Ttyh3 |
| C2.CP | REACTOME_CIRCADIAN_CLOCK | WT-TKO_up | 9 | 574 | 29 | 8100 | 4.37943049 | 0.000117 | 0.0224917 | Bhlhe40;Cry2;Dbp;Klf15;Nr1d1;Per1;Per2;Rxra;Ubc |
| C2.CP | REACTOME_MISCELLANEOUS_SUBSTRATES | WT-TKO_up | 4 | 574 | 5 | 8100 | 11.2891986 | 0.000118 | 0.0224917 | Cyp2s1;Cyp2u1;Cyp4b1;Cyp4f18 |
| C2.CP | REACTOME_RHO_GTPASES_ACTIVATE_RHOTEKIN_AND_RHOPHILINS | WT-TKO_up | 4 | 574 | 5 | 8100 | 11.2891986 | 0.000118 | 0.0224917 | Rhob;Rhoc;Rhpn2;Rtkn |
| C5.BP | GO_N_ACETYLNEURAMINATE_CATABOLIC_PROCESS | WT-TKO_up | 4 | 574 | 5 | 8100 | 11.2891986 | 0.000118 | 0.0224917 | Amdhd2;Gnpda1;Gnpda2;Npl |
| C5.BP | GO_ION_TRANSMEMBRANE_TRANSPORT | WT-TKO_up | 52 | 574 | 437 | 8100 | 1.67917142 | 0.000127 | 0.02382966 | Abcb1a;Ank3;Anxa2;Atp1a1;Atp1b1;Atp2a3;Atp2b1;Bin1;Chp1;Clca4a;Clcn5;Cldn4;Clic1;Cln8;Hap1;Irs2;Itpr2;Lrrc8a;Mcoln1;Nipal1;Otop2;Per2;Plcd1;Pm20d1;Prkaa2;Prnp;S100a6;Scnn1a;Sgk3;Slc13a1;Slc15a1;Slc20a1;Slc23a1;Slc25a22;Slc26a11;Slc2a10;Slc2a8;Slc30a2;Slc37a4;Slc3a1;Slc4a11;Slc4a7;Slc5a6;Slc6a19;Slc6a6;Slc6a8;Slc7a8;Slc9a3r1;Tmc7;Tmem63b;Ttyh3;Vamp2 |
| C5.BP | GO_RESPONSE_TO_LIPID | WT-TKO_up | 51 | 574 | 428 | 8100 | 1.68151031 | 0.000142 | 0.02572014 | Abcc2;Abhd2;Adcy2;Anxa3;Arnt2;Atp1a1;Atp2b1;Bcr;Capn2;Cldn4;Cpn1;Cry2;Cx3cr1;Dgat2;Dusp10;Endog;Fdx1;Gch1;Gjb3;Got1;Hmgcs1;Hpgd;Hsd3b2;Hsd3b3;Idh1;Il15;Insig1;Klf4;Klf9;Ldlr;Maob;Mrc1;Nr1d1;Nr1d2;Nr1h3;Nr3c2;Per1;Pmepa1;Pparg;Prkaa2;Rb1;Rorc;Runx1;Rxra;S100a14;Sdc1;Slc6a4;Srebf2;Stxbp1;Wbp2;Zfp703 |
| C5.BP | GO_STEROL_BIOSYNTHETIC_PROCESS | WT-TKO_up | 12 | 574 | 50 | 8100 | 3.38675958 | 0.000143 | 0.02572014 | Ces1b;Fdft1;Fdps;Hmgcs1;Insig1;Lss;Msmo1;Mvk;Nsdhl;Por;Prkaa2;Srebf2 |
| C5.BP | GO_REGULATION_OF_STEROID_METABOLIC_PROCESS | WT-TKO_up | 14 | 574 | 65 | 8100 | 3.03939963 | 0.000143 | 0.02572014 | Atp1a1;Bmp2;Dgat2;Fdft1;Fdps;Hmgcs1;Insig1;Ldlr;Lss;Mvk;Nr1d1;Por;Rorc;Srebf2 |
| C5.BP | GO_INORGANIC_ANION_TRANSMEMBRANE_TRANSPORT | WT-TKO_up | 11 | 574 | 43 | 8100 | 3.60991816 | 0.000146 | 0.02585775 | Clca4a;Clcn5;Cldn4;Clic1;Prnp;Slc13a1;Slc20a1;Slc26a11;Slc37a4;Slc4a11;Ttyh3 |
| C5.MF | GO_MONOVALENT_INORGANIC_CATION_TRANSMEMBRANE_TRANSPORTER_ACTIVITY | WT-TKO_up | 22 | 574 | 133 | 8100 | 2.3342328 | 0.000148 | 0.02585775 | Atp1a1;Atp1b1;Atp2a3;Clcn5;Otop2;Scnn1a;Slc13a1;Slc13a2;Slc15a1;Slc20a1;Slc23a1;Slc2a10;Slc2a8;Slc4a11;Slc4a7;Slc5a1;Slc5a4a;Slc5a6;Slc6a19;Slc6a4;Slc6a6;Slc6a8 |
| C5.BP | GO_CELLULAR_LIPID_METABOLIC_PROCESS | WT-TKO_up | 59 | 574 | 518 | 8100 | 1.6072942 | 0.000152 | 0.02593349 | Abhd1;Abhd2;Abhd5;Acaa1a;Acox2;Acsm5;Acss1;B4galt5;B4galt6;Bco1;Capn2;Ces1b;Ces2a;Ces2c;Cish;Cln8;Cwh43;Cyp2j6;Cyp2s1;Cyp2u1;Cyp4f18;Dgat2;Dgkd;Elovl1;Enpp7;Fdft1;Fdps;Gpd1;Hmgcs1;Hpgd;Idh1;Inpp5a;Insig1;Irs2;Klf4;Ldlr;Lpin3;Lss;Msmo1;Mvk;Naaa;Neu1;Nr1h3;Per2;Phospho1;Pisd;Plcd1;Plekha8;Plscr1;Pm20d1;Por;Pparg;Prkaa2;Sdc1;Sdc3;St8sia6;Trib3;Ugcg;Wdtc1 |
| C5.BP | GO_MONOVALENT_INORGANIC_CATION_TRANSPORT | WT-TKO_up | 27 | 574 | 180 | 8100 | 2.11672474 | 0.000153 | 0.02593349 | Ank3;Atp1a1;Atp1b1;Atp2a3;Bin1;Chp1;Clcn5;Otop2;Per1;Pm20d1;Prnp;Scnn1a;Slc13a1;Slc13a2;Slc15a1;Slc20a1;Slc23a1;Slc2a10;Slc2a8;Slc4a11;Slc4a7;Slc5a1;Slc5a4a;Slc5a6;Slc6a8;Slc9a3r1;Vamp2 |
| C5.BP | GO_REGULATION_OF_GLIAL_CELL_DIFFERENTIATION | WT-TKO_up | 9 | 574 | 30 | 8100 | 4.23344948 | 0.000157 | 0.02626167 | Bin1;Bmp2;Dusp10;Il34;Ldlr;Nr1d1;Pparg;Spint1;Tnfrsf21 |
| C5.BP | GO_EPIDERMIS_DEVELOPMENT | WT-TKO_up | 22 | 574 | 134 | 8100 | 2.31681315 | 0.000166 | 0.0273673 | Acvr1b;Apcdd1;Bcr;Emp1;Fosl2;Foxq1;Jup;Klf4;Krt20;Krt23;Krt8;Lamb3;Lamc2;Maff;Nsdhl;Palld;Pkp3;Runx1;Slc4a7;Slc9a3r1;Sult2b1;Ugcg |
| C5.BP | GO_ORGANIC_ACID_TRANSPORT | WT-TKO_up | 25 | 574 | 162 | 8100 | 2.17770035 | 0.000169 | 0.02752023 | Abat;Abcc2;Abcc3;Cln8;Cyp4f18;Irs2;Lrrc8a;Per2;Pparg;Prkaa2;Rxra;Slc13a1;Slc13a2;Slc16a5;Slc23a1;Slc25a22;Slc3a1;Slc5a6;Slc6a19;Slc6a6;Slc6a8;Slc7a8;Slc9a3r1;Stxbp1;Vamp2 |
| C5.BP | GO_NEUROTRANSMITTER_CATABOLIC_PROCESS | WT-TKO_up | 5 | 574 | 9 | 8100 | 7.83972125 | 0.000174 | 0.02798896 | Abat;Maoa;Maob;Por;Sardh |

| | | | | | | | | | | |
|---|---|---|---|---|---|---|---|---|---|---|
| C5.BP | GO_RESPONSE_TO_METAL_ION | WT-TKO_up | 26 | 574 | 173 | 8100 | 2.12080321 | 0.000195 | 0.03099443 | Abat;Abcc2;Ank3;Anxa7;Bace1;Bnip3;Cpne2;Dpep1;Eef2k;Endog;Fbp1;Gclc;Got1;Hmox1;Maob;Mcoln1;Mt1;Mt2;Ndrg1;Prkaa2;Prnp;Qdpr;Sdc1;Slc30a2;Syt12;Syt7 |
| C5.MF | GO_STEROID_HORMONE_RECEPTOR_ACTIVITY | WT-TKO_up | 8 | 574 | 25 | 8100 | 4.51567944 | 0.000222 | 0.03480232 | Abhd2;Nr1d1;Nr1d2;Nr1h3;Nr3c2;Pparg;Rorc;Rxra |
| C5.BP | GO_RESPONSE_TO_DRUG | WT-TKO_up | 56 | 574 | 492 | 8100 | 1.60618679 | 0.000229 | 0.03535051 | Abat;Abcb1a;Abcc2;Abhd2;Actb;Adcy2;Atp1a1;Bcar3;Bnip3;Capn2;Cldn4;Cln8;Cyp2j6;Cyp2s1;Cyp2u1;Cyp4b1;Dpep1;Endog;Fbp1;Fdx1;Fzd1;Gclc;Hmgcs1;Hmox1;Hpgd;Itpr2;Jup;Kdm6b;Klf4;Klf9;Maob;Mt2;Nat8;Nlrp1b;Nr1h3;P2ry2;Per2;Por;Pparg;Prkaa2;Prnp;Qdpr;Rgs17;Rhob;Sdc1;Slc13a1;Slc25a22;Slc4a11;Slc5a6;Slc6a4;Slc6a8;Slc9a3r1;Syt12;Syt7;Trdmt1;Wbp2 |
| C5.BP | GO_NEUROGENESIS | WT-TKO_up | 69 | 574 | 641 | 8100 | 1.51902243 | 0.000232 | 0.03544732 | Actb;Als2;Ank3;Apcdd1;Asap1;B4galt5;B4galt6;Bhlhe40;Bin1;Bmp2;Bnip3;Cln8;Cx3cr1;Dclk2;Dpysl2;Dusp10;Dvl2;Eef2k;Efhd1;Eml1;Epha1;Ezr;Fzd1;Fzd8;Hap1;Ifrd1;Il34;Irs2;Kalrn;Klf15;Klf4;Lamc2;Ldlr;Lpar3;Mapk6;Mmp24;Myh10;Nbl1;Ndrg1;Nptn;Nr1d1;Obsl1;Palld;Pard6b;Per2;Plk5;Plxna2;Pmp22;Pparg;Ppp2r5b;Prag1;Rapgef1;Rb1;Runx1;S100a6;Sema3b;Sema7a;Sh3gl2;Slc4a7;Slc6a4;Slc9a3r1;Spint1;Stxbp1;Tnfrsf12a;Tnfrsf21;Tubb2a;Ugcg;Wee1;Zswim4 |
| C5.BP | GO_DRUG_CATABOLIC_PROCESS | WT-TKO_up | 13 | 574 | 61 | 8100 | 3.00736848 | 0.000275 | 0.04147779 | Amdhd2;Cpn1;Cyp2j6;Cyp2s1;Cyp2u1;Cyp4b1;Gnpda1;Gnpda2;Maoa;Maob;Npl;Por;Qdpr |
| C5.BP | GO_XENOBIOTIC_METABOLIC_PROCESS | WT-TKO_up | 15 | 574 | 78 | 8100 | 2.71374967 | 0.000317 | 0.04715966 | Acaa1a;Acy3;Arnt2;Ces1b;Ces2a;Ces2c;Cryz;Cyp2j6;Cyp2s1;Cyp2u1;Dpep1;Gstm2;Por;Rorc;Ugt2b5 |
| C5.BP | GO_OXIDATION_REDUCTION_PROCESS | WT-TKO_up | 48 | 574 | 409 | 8100 | 1.65611716 | 0.00032 | 0.04715966 | Acaa1a;Acox2;Acss1;Akr1e1;Bco1;Bmp2;Bnip3;Coq10b;Cryz;Cyp2j6;Cyp2s1;Cyp2u1;Cyp4b1;Cyp4f18;Dgat2;Enox1;Fdx1;Flcn;Gfod1;Glud1;Gpd1;Gpd2;Hmox1;Hpgd;Hsd17b13;Hsd3b2;Hsd3b3;Idh1;Irs2;Kdm6b;Maoa;Maob;Me2;Me3;Mical2;Msmo1;Nr1d1;Nsdhl;Per2;Pgm2;Por;Pparg;Qdpr;Sardh;Slc25a22;Srd5a2;Stk40;Vat1 |
| C5.MF | GO_ORGANIC_ACID_SODIUM_SYMPORTER_ACTIVITY | WT-TKO_up | 5 | 574 | 10 | 8100 | 7.05574913 | 0.000328 | 0.04770031 | Slc13a1;Slc13a2;Slc23a1;Slc6a6;Slc6a8 |
| C5.BP | GO_INTESTINAL_HEXOSE_ABSORPTION | WT-TKO_up | 4 | 574 | 6 | 8100 | 9.40766551 | 0.000334 | 0.04795007 | Ezr;Slc2a2;Slc2a5;Slc5a1 |
| C5.BP | GO_CELLULAR_RESPONSE_TO_HORMONE_STIMULUS | WT-TKO_up | 38 | 574 | 302 | 8100 | 1.77561899 | 0.000348 | 0.04817943 | Abcc2;Abhd2;Adcy2;Atp1a1;Atp2b1;Bcar1;Cry2;Eef2k;Foxo3;Gclc;Got1;Hmgcs1;Irs2;Klf9;Lats2;Lpin3;Myo1c;Nr1d1;Nr1d2;Nr1h3;Nr3c2;Per1;Pmepa1;Por;Pparg;Prkaa2;Rab31;Rb1;Rorc;Runx1;Rxra;Slc2a8;Trib3;Ugcg;Vamp2;Wbp2;Wdtc1;Zfp703 |
| C2.CP | BIOCARTA_FXR_PATHWAY | WT-TKO_up | 3 | 574 | 3 | 8100 | 14.1114983 | 0.000354 | 0.04817943 | Ldlr;Nr1h3;Rxra |
| C5.BP | GO_DETOXIFICATION_OF_INORGANIC_COMPOUND | WT-TKO_up | 3 | 574 | 3 | 8100 | 14.1114983 | 0.000354 | 0.04817943 | Abcc2;Mt1;Mt2 |
| C5.MF | GO_3_BETA_HYDROXY_DELTA5_STEROID_DEHYDROGENASE_ACTIVITY | WT-TKO_up | 3 | 574 | 3 | 8100 | 14.1114983 | 0.000354 | 0.04817943 | Hsd3b2;Hsd3b3;Nsdhl |
| C5.MF | GO_SUGAR_TRANSMEMBRANE_TRANSPORTER_AC | WT-TKO_up | 6 | 574 | 15 | 8100 | 5.6445993 | 0.000355 | 0.04817943 | Slc23a1;Slc2a2;Slc2a5;Slc2a8;Slc5a1;Slc5a4a |
| C5.BP | GO_ALCOHOL_BIOSYNTHETIC_PROCESS | WT-TKO_up | 17 | 574 | 96 | 8100 | 2.49891115 | 0.000362 | 0.0486327 | Bmp2;Ces1b;Fdft1;Fdps;Gch1;Got1;Hmgcs1;Insig1;Lss;Msmo1;Mvk;Nsdhl;Plcd1;Por;Prkaa2;Qdpr;Srebf2 |
| C5.MF | GO_INORGANIC_ANION_TRANSMEMBRANE_TRANSP | WT-TKO_up | 12 | 574 | 55 | 8100 | 3.07887235 | 0.000373 | 0.0495648 | Clca4a;Clcn5;Cldn4;Clic1;Lrrc8a;Slc13a1;Slc26a11;Slc37a4;Slc4a11;Slc4a7;Slc6a4;Ttyh3 |

### Supplemental Table 2 GOs and upregulated genes in TKO

| MSigDB | Contrast | Overlap Size | Contrast Size | MSigDB Size | Background Size | FE | Pvalue | P.adj | Overlap.Items | neglogP |
|---|---|---|---|---|---|---|---|---|---|---|
| GO_RIBONUCLEOPROTEIN_COMPLEX_BIOGENESIS | WT-TKO_down | 67 | 657 | 192 | 8100 | 4.3 | 1.32E-26 | 1.62E-22 | Abce1;Atm;C1qbp;Cebpz;Celf5;Ddx18;Ddx21;Ddx27;Dimt1;Dis3;Dkc1;Eif3a;Emg1;Esf1;Exosc8;Fastkd2;Ftsj3;Gar1;Gemin5;Gtpbp4;Heatr1;Heatr3;Hsp90aa1;Hsp90ab1;Luc7l;Luc7l3;Lyar;Mdn1;Mphosph10;Nat10;Nhp2;Nip7;Nob1;Noc2l;Nol8;Nol9;Nolc1;Nom1;Nop14;Nop56;Nop58;Npm3;Ppan;Prmt5;Psip1;Riok1;Rpl13a;Rpp38;Rpusd2;Rrp15;Rrp1b;Rrp8;Rrs1;Rsl1d1;Ruvbl1;Snrpg;Surf6;Tex10;Tsr1;Txnl4a;Utp14a;Wdr12;Wdr18;Wdr3;Wdr43;Znhit6;Zrsr2 | 21.7913 |
| GO_CHROMOSOME_ORGANIZATION | WT-TKO_down | 98 | 657 | 429 | 8100 | 2.8 | 2.04E-22 | 1.25E-18 | Anp32b;Arrb1;Atad2;Atm;Aurka;Bub1;Bub1b;Ccna2;Ccnb1;Cdc20;Cdc45;Cdca5;Cdca8;Cdk1;Cenpa;Cenpe;Cenpf;Cenpi;Cenpw;Dcp2;Ddx11;Dkc1;Dlgap5;Dnajc2;Dpf3;Dtx3l;Ercc6l;Fancd2;Gar1;Gnl3;Gtf2h4;Gtf3c4;Hells;Hirip3;Hmgb2;Hmgn1;Hsp90aa1;Hsp90ab1;Incenp;Kat2a;Kif18b;Kif22;Kif23;Kifc1;Knstrn;L3mbtl2;Leo1;Mcm2;Mcm4;Mcm5;Mcm7;Mettl4;Mki67;Myb;Myc;Nap1l1;Nat10;Ncapd2;Ncaph;Ncor1;Ndc1;Ndc80;Nek2;Nhp2;Noc2l;Npm3;Nucks1;Nuf2;Nup155;Pif1;Pinx1;Plk1;Prc1;Prmt5;Psrc1;Rad54l;Rfc1;Rfc4;Rnf168;Rpa2;Rrp8;Rrs1;Ruvbl1;Setd6;Smc2;Spag5;Spdl1;Supt16;Syce2;Tacc3;Tert;Top2a;Trim37;Ttk;Twnk;Uhrf1;Usp16;Zfp451 | 17.9039 |
| GO_RIBOSOME_BIOGENESIS | WT-TKO_down | 52 | 657 | 144 | 8100 | 4.5 | 1.29E-21 | 5.26E-18 | Abce1;C1qbp;Cebpz;Ddx18;Ddx21;Ddx27;Dimt1;Dis3;Dkc1;Emg1;Esf1;Exosc8;Fastkd2;Ftsj3;Gar1;Gtpbp4;Heatr1;Heatr3;Lyar;Mdn1;Mphosph10;Nat10;Nhp2;Nip7;Nob1;Noc2l;Nol8;Nol9;Nolc1;Nom1;Nop14;Nop56;Nop58;Npm3;Ppan;Riok1;Rpp38;Rpusd2;Rrp15;Rrp1b;Rrp8;Rrs1;Rsl1d1;Surf6;Tex10;Tsr1;Utp14a;Wdr12;Wdr18;Wdr3;Wdr43;Znhit6 | 17.2786 |
| GO_RNA_PROCESSING | WT-TKO_down | 80 | 657 | 321 | 8100 | 3.1 | 7.63E-21 | 2.33E-17 | C1qbp;Ccnb1;Celf5;Cpsf2;Cwf19l2;Dcps;Ddx18;Ddx21;Ddx27;Dimt1;Dis3;Dkc1;Emg1;Ern1;Esf1;Exosc8;Ftsj1;Ftsj3;Gar1;Gemin5;Gtf2h4;Gtpbp4;Heatr1;Kti12;Leo1;Luc7l;Luc7l3;Lyar;Mdn1;Mphosph10;Nat10;Ncbp2;Ncl;Ncor1;Nhp2;Nob1;Nol8;Nol9;Nolc1;Nop14;Nop56;Nop58;Npm3;Polr2l;Ppie;Ppih;Ppwd1;Prmt5;Prpf4b;Psip1;Pus7;Rbm25;Rbm28;Riok1;Rpp38;Rpusd2;Rrp15;Rrp1b;Rrp8;Rrs1;Rsl1d1;Snrpa1;Snrpg;Tert;Tex10;Thg1l;Trit1;Trmt13;Tsr1;Txnl4a;Upf3b;Utp14a;Wdr12;Wdr18;Wdr3;Wdr4;Wdr43;Zfc3h1;Znhit6;Zrsr2 | 16.633 |
| GO_RRNA_METABOLIC_PROCESS | WT-TKO_down | 44 | 657 | 114 | 8100 | 4.8 | 9.04E-20 | 1.89E-16 | Ddx18;Ddx21;Ddx27;Dimt1;Dis3;Dkc1;Emg1;Esf1;Exosc8;Ftsj3;Gar1;Gtpbp4;Heatr1;Lyar;Mdn1;Mphosph10;Nat10;Nhp2;Nifk;Nob1;Nol8;Nol9;Nolc1;Nop14;Nop56;Nop58;Npm3;Riok1;Rpp38;Rpusd2;Rrp15;Rrp1b;Rrp8;Rrs1;Rsl1d1;Slfn9;Tex10;Tsr1;Utp14a;Wdr12;Wdr18;Wdr3;Wdr43;Znhit6 | 15.723 |
| GO_RNA_BINDING | WT-TKO_down | 118 | 657 | 622 | 8100 | 2.3 | 9.30E-20 | 1.89E-16 | 2810004N23Rik;3110082I17Rik;Aldh18a1;C1qbp;Cebpz;Celf5;Cpsf2;Dcp2;Dcps;Ddx11;Ddx18;Ddx21;Ddx27;Dhx29;Dhx33;Dimt1;Dis3;Dkc1;Dnajc2;Dzip3;Eif3a;Eif5b;Emg1;Esf1;Exosc8;Fastkd2;Ftsj3;Gar1;Gemin5;Gnl3;Gpatch4;Gtpbp4;Heatr1;Hmgb2;Hsp90aa1;Hsp90ab1;Hspd1;Hspe1;Ifit1;Ifit1bl1;Ifit2;Impdh2;Knop1;Luc7l;Luc7l3;Lyar;Mak16;Map3k20;Mex3a;Mki67;Mphosph10;Mrps27;Mrps5;Msi1;Myef2;Naa15;Nap1l1;Nat10;Ncbp2;Ncl;Nhp2;Nifk;Nip7;Noc2l;Noc3l;Nol12;Nol8;Nol9;Nolc1;Nom1;Nop14;Nop56;Nop58;Npm3;Nucks1;Pinx1;Ppan;Ppargc1b;Ppie;Pprc1;Prpf4b;Prrc2c;Psip1;Pus7;Rad51ap1;Rangap1;Rbm25;Rbm28;Rnaset2b;Rpl13a;Rpp38;Rpusd2;Rrbp1;Rrp1b;Rrp8;Rrs1;Rsl1d1;Sidt1;Slfn9;Snrpa1;Snrpg;Supt16;Surf6;Synj2;Tcof1;Tert;Thg1l;Top2a;Trim21;Tsr1;Tuba1b;Upf3b;Utp14a;Wdr3;Wdr43;Zcchc17;Zfc3h1;Zrsr2 | 15.723 |
| GO_NUCLEOLUS | WT-TKO_down | 92 | 657 | 427 | 8100 | 2.7 | 3.34E-19 | 5.83E-16 | Aen;Anp32b;Arntl2;Atm;C1qbp;Cbx5;Cdca8;Cenpw;Cep85;Ddx11;Ddx18;Ddx21;Ddx27;Dhx33;Dimt1;Dkc1;Dph6;Eif3a;Emg1;Esf1;Exosc8;Fancd2;Ftsj3;Gar1;Gnl3;Gtpbp4;Heatr1;Hirip3;Hmgb2;Ier5;Knop1;Leo1;Lyar;Mak16;Mdn1;Mki67;Mphosph10;Mrps27;Myc;Myo10;Nat10;Ncl;Nek2;Nepro;Nhp2;Nifk;Nip7;Noc2l;Noc3l;Nol12;Nol8;Nol9;Nolc1;Nom1;Nop14;Nop56;Nop58;Npm3;Nucks1;Pinx1;Polr1a;Polr2l;Ppan;Pycard;Rbm28;Rfc1;Rnf213;Rpl13a;Rpp38;Rrp1b;Rrp8;Rrs1;Rsl1d1;Smc2;Surf6;Tcof1;Tert;Tex10;Top2a;Tsr1;Ttc3;Tulp3;Upf3b;Utp14a;Vmp1;Wdr12;Wdr18;Wdr3;Wdr43;Ypel2;Zcchc17;Zfp692 | 15.234 |
| GO_MITOTIC_NUCLEAR_DIVISION | WT-TKO_down | 47 | 657 | 142 | 8100 | 4.1 | 7.32E-18 | 1.12E-14 | Aaas;Atm;Aurka;Birc5;Bub1;Bub1b;Ccnb1;Cdc20;Cdc25c;Cdca5;Cdca8;Cenpe;Cenpf;Cep192;Cep85;Cep97;Dlgap5;Incenp;Kif18b;Kif22;Kif23;Kifc1;Klhdc8b;Knstrn;Mki67;Mybl2;Ncapd2;Ncaph;Ndc80;Nde1;Nek2;Nuf2;Pinx1;Plk1;Prc1;Prmt5;Psrc1;Ranbp1;Rrs1;Smc2;Spag5;Spdl1;Tacc3;Tpx2;Ttk;Ube2c;Usp16 | 13.952 |
| GO_RIBONUCLEOPROTEIN_COMPLEX | WT-TKO_down | 58 | 657 | 214 | 8100 | 3.3 | 3.87E-17 | 5.25E-14 | Cbx5;Celf5;Cwf19l2;Dcp2;Dhx29;Dkc1;Eif3a;Emg1;Ftsj3;Gar1;Hba-a1;Heatr1;Luc7l;Luc7l3;Mak16;Mphosph10;Mrpl19;Mrps27;Mrps5;Msi1;Myef2;Nat10;Ncl;Nhp2;Nip7;Nob1;Noc2l;Nol8;Nolc1;Nop14;Nop56;Nop58;Ppan;Ppie;Ppih;Ppwd1;Prpf4b;Rbm28;Riok1;Rpl13a;Rpp38;Rrp15;Rrp1b;Rrs1;Rsl1d1;Ruvbl1;Snrpa1;Snrpg;Tert;Top2a;Trim21;Tsr1;Txnl4a;Utp14a;Wdr12;Wdr3;Znhit6;Zrsr2 | 13.28 |
| REACTOME_METABOLISM_OF_RNA | WT-TKO_down | 59 | 657 | 231 | 8100 | 3.1 | 4.19E-16 | 5.12E-13 | Aaas;Cpsf2;Dcp2;Dcps;Ddx21;Dimt1;Dis3;Dkc1;Emg1;Exosc8;Ftsj1;Ftsj3;Gar1;Gemin5;Gnl3;Gtf2h4;Heatr1;Mphosph10;Nat10;Ncbp2;Ncl;Ndc1;Nhp2;Nip7;Nob1;Nol12;Nol9;Nop14;Nop56;Nop58;Nup133;Nup155;Nup85;Polr2l;Ppie;Ppih;Ppwd1;Prmt5;Pus7;Rbm28;Riok1;Rpl13a;Rpp38;Snrpa1;Snrpg;Tex10;Thg1l;Trit1;Trmt13;Tsr1;Txnl4a;Upf3b;Utp14a;Wdr12;Wdr18;Wdr3;Wdr4;Wdr43;Zrsr2 | 12.291 |
| GO_NCRNA_PROCESSING | WT-TKO_down | 49 | 657 | 169 | 8100 | 3.6 | 6.36E-16 | 7.06E-13 | Ddx18;Ddx21;Ddx27;Dimt1;Dis3;Dkc1;Emg1;Esf1;Exosc8;Ftsj1;Ftsj3;Gar1;Gtpbp4;Heatr1;Kti12;Lyar;Mdn1;Mphosph10;Nat10;Nhp2;Nob1;Nol8;Nol9;Nolc1;Nop14;Nop56;Nop58;Npm3;Pus7;Riok1;Rpp38;Rpusd2;Rrp15;Rrp1b;Rrp8;Rrs1;Rsl1d1;Tex10;Thg1l;Trit1;Trmt13;Tsr1;Utp14a;Wdr12;Wdr18;Wdr3;Wdr4;Wdr43;Znhit6 | 12.151 |
| REACTOME_CELL_CYCLE | WT-TKO_down | 72 | 657 | 331 | 8100 | 2.7 | 2.20E-15 | 2.24E-12 | Aaas;Atm;Aurka;Birc5;Bub1;Bub1b;Ccna2;Ccnb1;Ccp110;Cdc20;Cdc25c;Cdc45;Cdc7;Cdca5;Cdca8;Cdk1;Cenpa;Cenpe;Cenpf;Cenpi;Cenpw;Cep192;Cep290;Cntrl;Dbf4;Dkc1;E2f2;Ercc6l;Foxm1;Gar1;Gtse1;Hmmr;Hsp90aa1;Hsp90ab1;Incenp;Kif23;Mcm2;Mcm4;Mcm5;Mcm7;Mdm4;Mybl2;Myc;Ncapd2;Ncaph;Ndc1;Ndc80;Nde1;Nek2;Nhp2;Nuf2;Nup133;Nup155;Nup85;Pif1;Plk1;Rangap1;Rfc1;Rfc4;Rnf168;Rpa2;Ruvbl1;Smc2;Spdl1;Syce2;Tert;Top2a;Tpx2;Trp53bp1;Tuba1b;Ube2c;Zwilch | 11.649 |
| GO_CONDENSED_CHROMOSOME | WT-TKO_down | 37 | 657 | 107 | 8100 | 4.3 | 5.29E-15 | 4.98E-12 | Aurka;Birc5;Bub1;Bub1b;Ccnb1;Cdca5;Cenpa;Cenpe;Cenpf;Cenpw;Ddx11;Ercc6l;Fancd2;Hmgb2;Incenp;Knstrn;Mki67;Ncapd2;Ncaph;Ndc80;Nde1;Nek2;Nifk;Nuf2;Nup133;Nup85;Pinx1;Plk1;Rangap1;Rrs1;Smc2;Spag5;Spdl1;Syce2;Top2a;Trp53bp1;Zwilch | 11.303 |
| GO_ORGANELLE_FISSION | WT-TKO_down | 55 | 657 | 217 | 8100 | 3.1 | 6.32E-15 | 5.52E-12 | Aaas;Aspm;Atm;Aurka;Birc5;Bub1;Bub1b;Ccnb1;Cdc20;Cdc25c;Cdca5;Cdca8;Cenpe;Cenpf;Cep192;Cep85;Cep97;Cks2;Dlgap5;Fancd2;Incenp;Kif18b;Kif22;Kif23;Kifc1;Klhdc8b;Knstrn;Mki67;Mybl2;Ncapd2;Ncaph;Ndc1;Ndc80;Nde1;Nek2;Nuf2;Pinx1;Plk1;Prc1;Prmt5;Psrc1;Rad51b;Rad54l;Ranbp1;Rrs1;Smc2;Spag5;Spdl1;Syce2;Tacc3;Top2a;Tpx2;Ttk;Ube2c;Usp16 | 11.258 |
| GO_NCRNA_METABOLIC_PROCESS | WT-TKO_down | 51 | 657 | 195 | 8100 | 3.2 | 1.65E-14 | 1.29E-11 | Ddx18;Ddx21;Ddx27;Dimt1;Dis3;Dkc1;Emg1;Esf1;Exosc8;Ftsj1;Ftsj3;Gar1;Gtpbp4;Heatr1;Kti12;Lyar;Mdn1;Mphosph10;Nat10;Nhp2;Nifk;Nob1;Nol8;Nol9;Nolc1;Nop14;Nop56;Nop58;Npm3;Pus7;Riok1;Rpp38;Rpusd2;Rrp15;Rrp1b;Rrp8;Rrs1;Rsl1d1;Slfn9;Tex10;Thg1l;Trit1;Trmt13;Tsr1;Utp14a;Wdr12;Wdr18;Wdr3;Wdr4;Wdr43;Znhit6 | 10.889 |
